# Aperiodic neural activity distinguishes between phasic and tonic REM sleep

**DOI:** 10.1101/2024.08.08.606670

**Authors:** Yevgenia Rosenblum, Tamás Bogdány, Lili Benedikta Nádasy, Ilona Kovács, Ferenc Gombos, Péter Ujma, Róbert Bódizs, Nico Adelhöfer, Péter Simor, Martin Dresler

**Affiliations:** Radboud University Medical Centre, Donders Institute for Brain, Cognition and Behavior, Nijmegen, Netherlands; Institute of Psychology, ELTE Eötvös Loránd University, Budapest, Hungary; Doctoral School of Psychology, ELTE, Eötvös Loránd University, Budapest, Hungary; HUN-REN-ELTE-PPKE Adolescent Development Research Group, Faculty of Education and Psychology, Eötvös Loránd University, Budapest, Hungary; Pázmány Péter Catholic University, Department of General Psychology, Budapest, Hungary; Semmelweis University, Institute of Behavioural Sciences, Budapest, Hungary

**Author notes:** <u>Corresponding author:</u> Yevgenia Rosenblum. equal contribution (Nico Adelhöfer, Péter Simor, Martin Dresler).

## Abstract

**Introduction:** Traditionally categorized as a uniform sleep phase, rapid eye movement (REM) sleep exhibits substantial heterogeneity with its phasic and tonic constituents showing marked differences regarding neuronal network activity, environmental alertness and information processing. Here, we investigate how tonic and phasic states differ with respect to aperiodic neural activity, a marker of arousal levels, sleep stages, depth of sleep and sleep intensity. We also attempt to challenge the binary categorisation of REM sleep states by introducing graduality into their definition. Specifically, we quantify the intensity of phasic oculomotor events and investigate their temporal relationships with aperiodic activity.

**Method:** We analyzed 57 polysomnographic recordings from three open-access datasets of healthy young volunteers aged 21.7±1.4 years. REM sleep heterogeneity was assessed using either binary phasic-tonic categorization or quantification of eye movement (EM) amplitudes detected by electrooculography with the YASA algorithm. Slopes of the aperiodic power component measured by electroencephalography in the low (2–30Hz) and high (30–48Hz) frequency bands were calculated using the Irregularly Resampled Auto-Spectral Analysis. For statistical analyses, we used ANOVA, Spearman correlations and cross-correlations.

**Results:** The binary approach revealed that the phasic state is characterized by steeper low-band aperiodic slopes compared to the tonic state with the strongest effect observed over the frontal area. The phasic state also showed flatter high-band slopes with the strongest effect over central and parietal areas. The gradual approach confirmed this result further showing that higher EM amplitudes are linked to steeper low-band and flatter high-band aperiodic slopes. The temporal analysis within REM episodes revealed that aperiodic activity preceding or following EM events did not cross-correlate with EM amplitudes.

**Conclusion:** This study demonstrates that aperiodic slopes can serve as a reliable objective marker able to differentiate between phasic and tonic constituents of REM sleep and reflect the intensity of phasic oculomotor events for instantaneous measurements. However, EM events could not be predicted by preceding aperiodic activity and vice versa, at least not with scalp electroencephalography.

## Introduction

### REM sleep states

Nocturnal human sleep has a cycling nature where each cycle comprises an episode of non-rapid eye movement (NREM) sleep followed by an episode of REM sleep (Aserinsky & Kleitman, 1953; Dement & Kleitman 1957). REM sleep in turn cycles between phasic states characterized by bursts of eye movements and tonic states that consist of longer and more quiescent segments (Ermis et al., 2010; Simor et al., 2020). Phasic and tonic periods exhibit remarkable differences concerning mental experience, environmental alertness, spontaneous and evoked cortical activity, information processing, neuronal network and autonomic (e.g., heart rate, skin conductance, respiration) activity (Simor et al., 2018; 2020). For example, phasic states show internally driven sensorimotor processing detached from the surroundings and therefore are often tagged as “offline” periods (Fogel et al., 2007; Datta et al., 2013). Tonic states are more responsive to external stimuli as evidenced by awakening and arousal thresholds and thus are often tagged as “online” periods (Simor et al., 2020). Notably, rather than discrete states, phasic and tonic states are sometimes seen as “ends of a continuum” (Bueno-Junior et al., 2023). In this study, we investigate whether heterogeneity of REM sleep (defined in either discrete or non-discrete ways) is reflected by aperiodic neural activity, a marker of states of vigilance, sleep stages, sleep depth and sleep intensity.

### Aperiodic activity

Aperiodic neural activity is a distinct type of brain dynamic that reflects the overall frequency composition of the neural signal, which is not dominated by any specific frequency (Bódizs et al., 2024). Aperiodic dynamics follow a power-law function, where power decreases with increasing frequency (He, 2014). The steepness of this decay is approximated by the spectral exponent, which is equivalent to the slope of the spectrum when plotted in the log-log space (Gerster et al., 2022). Steeper (more negative) slopes indicate that the spectral power is relatively stronger in slower frequencies and relatively weaker in faster ones and *vice versa* (Bódizs et al., 2024).

In 2017, Gao et al. suggested that high-band (30 – 50Hz) aperiodic slopes reflect the balance between excitatory and inhibitory neural currents, which in turn defines a specific arousal state and a conscious experience of an organism. Following this report, aperiodic activity has received increased attention, and now is being seen as a window into diverse neural processes associated with cognitive tasks, age and disease (Voytek & Knight, 2015; Bódizs et al., 2021; Höhn et al., 2024). In the field of sleep research, it has been shown that aperiodic slopes differentiate between sleep stages such that high-band slopes are steeper during REM than NREM sleep, whereas low-band and broadband slopes show the opposite pattern (Miscovic et al., 2019; Lendner et al., 2020; Kozhemiako et al., 2022; Schneider et al., 2022). Given the ability of aperiodic activity to reflect many parameters of sleep, such as intensity, depth, stages and cycles of sleep (Reviewed in Bódizs et al., 2024), we hypothesize that it would further differentiate between REM sleep constituents.

### Aims and hypotheses

To assess whether REM sleep heterogeneity is reflected by aperiodic activity, we used both binary categorization and the gradual approach that quantified eye movement (EM) amplitudes. We hypothesized that the phasic state/higher EM amplitudes would be associated with steeper aperiodic slopes in the low-frequency band. This is based on the previous finding on increased delta oscillations in phasic vs tonic states (Simor et al., 2020) as such a difference implies a relative predominance of lower frequencies in phasic spectral power and thus, a steeper decay. Regarding the high-frequency band, we hypothesized that the phasic states would be linked to flatter aperiodic slopes based on a reported increase in gamma oscillations in phasic vs tonic states (Simor et al., 2020). Such a difference implies the relative predominance of higher frequencies in this band and thus, slower/flatter spectral decay. In addition, we assessed temporal relationships between oculomotor and aperiodic activities across a duration of single REM sleep episodes. Here, we used cross-correlations, assuming that its shape would reveal information concerning the possible direction (leading vs lagging) of the influence between the two time series.

## Methods

### Datasets

We used three independently collected open-source polysomnographic datasets of healthy young individuals overall comprising 57 participants aged 21.5 ± 1.4 years of age. Information about the studies, participants, polysomnographic devices as well as the links to the publicly available data is reported in Table 1. All studies were approved by the corresponding Ethics committee. All participants gave written informed consent.

**Table 1:** Study and device information for each dataset.

| Characteristic | Dataset 1 | Dataset 2 | Dataset 3 |
| --- | --- | --- | --- |
| Described in: | Simor et al., 2018;<br>Simor et al., 2021 | Van der Wijk et al., 2020;<br>Simor et al., 2021 | Simor et al., 2019 |
| Data availability: | Categorical: https://osf.io/2vptx<br>Continuous: non available |  | Categorical:<br>https://osf.io/9k5hb<br>Continuous: available but non-open access |
| Approved by | The Ethical Committee of the Semmelweis University | The United Ethical Review Committee for Research in Psychology, Hungary (EBKEB 2016/077) | The Ethical Committee of the Pázmány Péter Catholic University for Psychological Experiments |
| No. participants | 20 | 17 | 20 |
| Age, years | 21.72 ± 1.36 | 21.46 ± 1.43 | 21.19 ± 0.43 |
| Gender, female, % | 50 | 70 | 44 |
| Polysomnographic device | Brain-Quick BQ 132S; Micromed, Mogliano Veneto, Italy | Micromed SD LTM 32 Bs (Micromed S.p.A., Mogliano Veneto, Italy) | BQ KIT SD, LTM 128 EXPRESS (2 – 64 channels in master and slave mode, Micromed, Mogliano Veneto, Italy) |
| No. channels | 19 | 17 | 128 |
| Referenced to | Mathematically linked mastoids (A1, A2) |  | Fronto-centrally placed ground electrode |
| Sample rate, Hz | 1024 | 512 | 512 |

The participants slept wearing a polysomnographic device in a sleep laboratory with an adaptation night before the examination night. Only the second-night data was analyzed. Sleep was scored by independent experts according to the AASM standards (Iber, 2007). Only clean REM epochs were used for further analysis.

### REM sleep binary segmentation

We used publicly available data preprocessed as described elsewhere (Simor et al., 2018; 2019; 2021; Van der Wijk et al., 2020). Briefly, first, the Independent Component Analysis (ICA) was applied to the data to correct for oculomotor artifacts (Simor et al., 2021). Then, the data was semi-manually inspected for rapid EMs with a custom-made software tool for full-night sleep EEG analysis (FerciosEEGPlus, Ferenc Gombos 2008–2017). EMs were visually identified in non-overlapping 4s time windows based on the presence of EOG deflections of amplitude above 150μV and shorter than 500ms. A 4s-long segment was categorized as phasic if at least two consecutive eye movements were detected in adjacent 2s time windows. Segments were scored as a tonic when no eye movements occurred (EOG deflections below 25μV) in adjacent 2s time windows. To avoid contamination between the two microstates, segments were only selected if they were at least 8s apart from each other.

### Spectral power

Offline EEG data analyses were carried out with MATLAB (version R2021b, The MathWorks, Inc., Natick, MA), using the Fieldtrip toolbox (Oostenveld et al., 2011) and custom-made scripts. For each REM state of each participant, we averaged the preprocessed EEG signal over four topographical areas: frontal (Fz, F3, F4); central (Cz, C3, C4); parietal (Pz, P3, P4); and occipital (O1, O2) separately to reduce the number of statistical comparisons.

We calculated the total spectral power of every 4s EEG segment and differentiated it into its aperiodic (i.e., fractal) and oscillatory components using the Irregularly Resampled Auto-Spectral Analysis (IRASA; Wen & Liu, 2016), the approach embedded in the Fieldtrip toolbox. To implement the algorithm, we used the *ft_freqanalysis* function of the Fieldtrip toolbox as described elsewhere (Rosenblum et al., 2023a; 2023b). Then, the aperiodic power component was transformed to log-log coordinates and its slope was calculated to estimate the power-law exponent (the rate of spectral decay), using the function *logfit* (Lansey, 2020; available on https://osf.io/zhyf7).

The signal was analyzed in the low and high-frequency bands separately. The low band was defined as 2 – 30Hz, being a representative of the typical sleep frequency range (0.3 – 30Hz), with the exclusion of frequencies < 2Hz in order to control for the possibility that slow frequency activity during phasic periods is contaminated by eye movements whose potentials mainly extend over 0.3 – 2Hz (Tan et al., 2001).

The high-band (30 – 48Hz) was analyzed as this range has been used for reliable discrimination between wakefulness and REM sleep (Lendner et al., 2020). Likewise, the 30 – 50Hz band was previously proposed to indicate excitation-to-inhibition balance (Gao et al., 2017). Given that the open access data for Datasets 1 and 2 had been filtered in the 0.5 – 35Hz range (Simor et al., 2021), the high-band analysis was performed for Dataset 3 only.

To replicate previous findings, we also calculated the oscillatory power component by subtracting the aperiodic component from the total power and averaging it over conventional frequency bands. This analysis is reported in Supplementary Material 1.

We analyzed the data merged from three datasets into one large dataset as well as each dataset individually. For the former analysis, we transformed the aperiodic slopes into z-scores since when we pooled together the data acquired by different devices, we detected significant differences between Datasets 1 and 3 vs Dataset 2 (See Fig.1). Specifically, for each participant, we merged the values calculated for the 4 topographical areas of the phasic and tonic states into one vector and then performed z-transformation. After the z-transform, we averaged a given variable over each area over each REM state separately.

**Figure 1.**
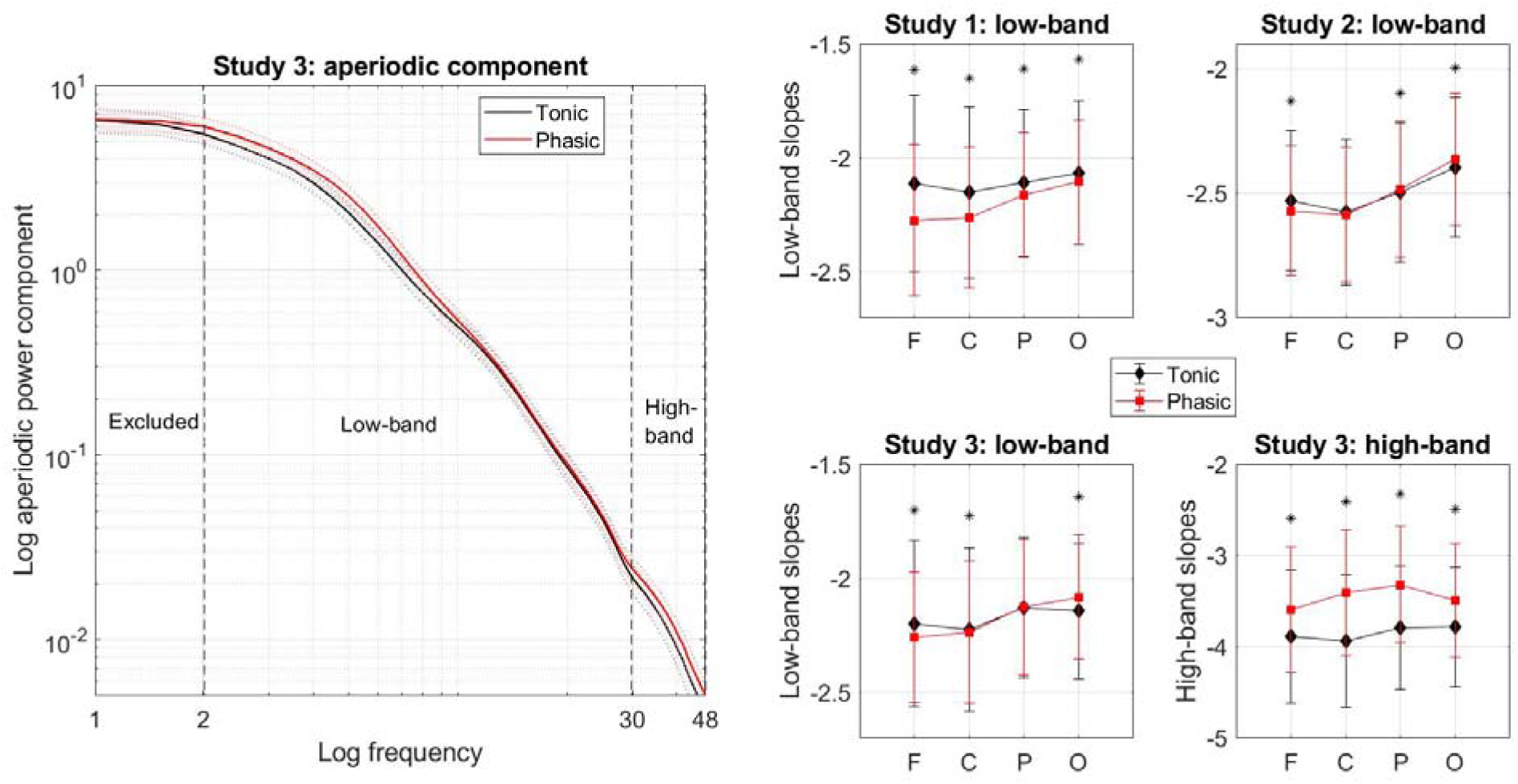
Aperiodic activity during tonic and phasic states. **Left:** Aperiodic component of spectral power averaged over frontal electrodes plotted as a function of frequency for tonic (black line) vs phasic (red line) states of REM sleep in Dataset 3. Shading indicates standard errors of the mean. The phasic state shows steeper decay of the aperiodic component in the low-frequency band (2 – 30Hz) and flatter decay in the high-frequency band (30 – 48Hz) compared to the tonic state. **Right:** Slopes of the aperiodic power component in the low band for Datasets 1 – 3 and in the high-band for Dataset 3 only averaged over the phasic vs tonic epochs over each topographical area separately. Overall, the tonic state (black diamonds) shows flatter low-band and steeper high-band aperiodic slopes compared to the phasic state (red squares) with some topographical variations in the strength and significance of the effect across the datasets. * – statistically significant difference between tonic and phasic states (p < 0.05), F – frontal, C – central, P – parietal, O – occipital.

Nevertheless, given that the raw values of spectral exponents/slopes have their own functional significance (as mentioned in more details in Discussion), we also analyzed each dataset individually while using raw values of aperiodic slopes. Raw slope values are reported in the Supplementary Excel File.

### Statistical analysis for the categorical approach

To compare aperiodic slopes during phasic vs tonic states, we used the ANOVA with the four-level “brain area” and two-level “REM sleep state” as within-subject factors. We performed ANOVAs for high and low-frequency bands separately with Benjamini-Hochberg’s adjustment to control for multiple (two) comparisons with a false discovery rate set at 0.05 and the α-level set in the 0.025 – 0.050 range. We applied Greenhouse-Geisser’s correction since Mauchly’s test revealed that the sphericity assumption was violated (ε < 0.75, p < 0.05). The assumptions of normality and homogeneity of variance were tested using the Q-Q plot and Levene’s homogeneity test, respectively.

Then, we performed *post hoc* analyses to compare binary phasic and tonic states for each topographical area separately using two-tailed Student’s paired t-tests. Effect sizes were calculated with Cohen’s d.

To assess the ability of the aperiodic slopes to discriminate between phasic and tonic states of REM sleep at the individual level, we calculated the area under the receiver operating characteristic (ROC) curve (AUC) for each topographical area. Benjamini-Hochberg’s adjustment was applied to control for multiple comparisons (8 tests corresponding to four areas for low and high bands separately) with a false discovery rate set at 0.05 and the α-level set in the 0.0063 (i.e., 0.05/8) – 0.0500 range. The sensitivity and specificity of the tests were evaluated using the median of all values (both phasic and tonic) of a given area as a cutoff. For the ROC and ANOVA, we used SPSS software (version 25; SPSS, Inc).

### EM quantification

To control for a possible bias of binary categorization of REM sleep into phasic and tonic states, we also used a gradual approach to define phasic oculomotor activity across the duration of REM sleep episodes. For this exploratory analysis, we used the continuous EOG data from Dataset 3 (n = 20) filtered in the 0.3 – 10Hz. We applied to it the *rem_detect* function of the YASA package for Python (Vallat & Walker, 2021) using its defaults (minimum amplitude: 50μV, duration: 0.3 – 1.2 seconds, rapid EMs frequency: 0.5 – 5Hz). As an outcome measure, we used the absolute peak amplitude of rapid EMs. This variable focuses on the EM morphology in relation to surrounding signal activity, ensuring that the identified peak is representative of an EM event rather than an isolated amplitude spike (Vallat & Walker, 2021).

We checked the EM detection by YASA via a visual inspection of the continuous EOG data. Likewise, we manually calculated the peak value of rapid EMs by identifying the maximum amplitude within a predefined 4s window based on raw amplitude measurements. The coefficient of correlation between the amplitude calculated by YASA and that calculated manually was 0.87. An in-depth analysis of one participant further showed that the percentage of agreement between the YASA algorithm and visual inspection (calculated as true positives + true negatives) equalled 91%. The percentage of rapid EMs missed by the YASA (false negatives) equalled 0.25%. The percentage of rapid EMs missed by manual coding (false positives) equalled 8.69%. Then, we correlated EM amplitudes with aperiodic slopes to obtain Spearman correlation coefficients.

### Temporal relationships between EMs and aperiodic slopes

To assess temporal relationships between oculomotor and aperiodic activities across the duration of a single REM sleep episode, we cross-correlated between the time series of EM amplitudes defined with continuous EOG (as described in the previous section) and the time series of aperiodic slopes calculated using continuous preprocessed EEG. Namely, preprocessing included filtering and the removal from the EEG signal independent component(s) related to EMs as described in Simor et al. (2019). Then, we calculated aperiodic slopes as described above. In order to approximate the distribution of the measured values to the normal distribution, both time series were ranked using the *tiedrank* function and correlation coefficients were transformed to Fisher z-scores. Confidence intervals of 95% were calculated to infer statistical significance. Cross-correlations were performed for the lags (i.e., temporal intervals between the two time series) lying between −5 and 5 minutes with a step of 4 seconds for low and high frequency bands for each topographical area and for each REM sleep episode separately.

A previous EOG study noted that EM density fluctuates at about 2-min intervals (Ktonas et al., 2003). Therefore, here, we included only episodes longer than 10 minutes to obtain the time series of sufficient length that, theoretically, would contain several periods of phasic bursts, and thus sufficient statistical power. Likewise, this selection increased the homogeneity of the analyzed episodes since early-night REM sleep episodes (which have been mainly excluded after this selection) are usually both considerably shorter and qualitatively different from the late-night ones because of circadian and homeostatic effects (Simor et al., 2023). This procedure resulted in 40 REM sleep episodes from 19 participants (in one participant, all REM sleep episodes were shorter than 10 minutes) that on average lasted for 17.7 ± 9.4 minutes.

Having performed cross-correlation at the individual episode level, next, we evaluated its episode-centered effect sizes to better understand the dynamics across all REM sleep episodes. For this, we counted the number of significant correlations between the two time series and divided it by the total number of all (significant and non-significant) performed correlations. Finally, we assessed the population prevalence of the person-centered effect sizes with the Bayesian prevalence. This method estimates the proportion of the population that would show the effect if they were tested in this experiment (Ince et al., 2022). As an output, it provides the maximum *a posterior* estimate – the most likely value of the population parameter and the highest posterior density intervals – the range within which the true population value lies with the specified probability level of 96%. To perform this analysis, we used an online web application available at https://estimate.prevalence.online.

## Results

### Categorical approach

Overall, the phasic state of REM sleep showed steeper (i.e., more negative) slopes of the aperiodic power component in the low (2 – 30Hz) band compared to the tonic state with some topographical variation across the datasets. Repeated-measures ANOVA performed for the pooled dataset (n = 55) revealed a significant effect of the area (F(51,4) = 47), state (F(53,1) = 11) as well as the state-area interaction (F(51,3) = 44; all p<0.001). The ROC analysis revealed that the low-band slopes measured over the frontal (AUC = 0.79 ± 0.015, p < 0.001) and central (AUC = 0.66 ± 0.015, p = 0.004) areas can discriminate between the two states.

*Post hoc* analysis performed for each dataset separately further showed that for Dataset 1, steeper phasic slopes could be observed over all areas with an anterior-to-posterior gradient of the strength of effect sizes with the largest difference measured over the frontal area. For Dataset 2, the effect could be observed over the frontal area only. For Dataset 3, the effect could be observed over the frontal and central areas. Interestingly, for Datasets 2 and 3, the occipital area showed the opposite direction of the effect: steeper aperiodic slopes during the tonic as compared to phasic state (Fig.1, Table 2).

**Table 2:**
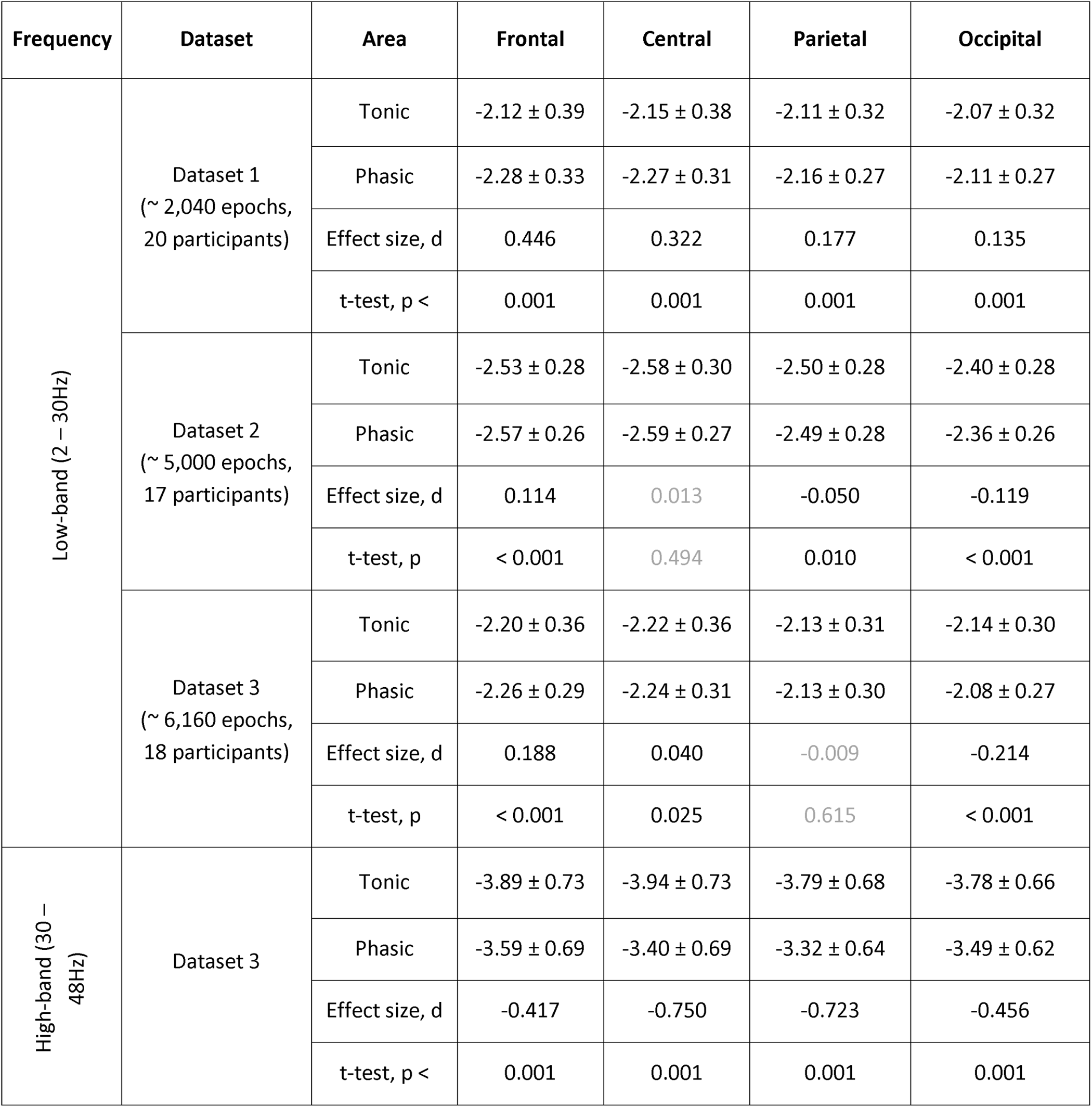
Aperiodic slopes.

| Frequency | Dataset | Area | Frontal | Central | Parietal | Occipital |
| --- | --- | --- | --- | --- | --- | --- |
| Low-band (2 – 30Hz) | Dataset 1<br>(~ 2,040 epochs,<br>20 participants) | Tonic | -2.12 ± 0.39 | -2.15 ± 0.38 | -2.11 ± 0.32 | -2.07 ± 0.32 |
|  |  | Phasic | -2.28 ± 0.33 | -2.27 ± 0.31 | -2.16 ± 0.27 | -2.11 ± 0.27 |
|  |  | Effect size, d | 0.446 | 0.322 | 0.177 | 0.135 |
|  |  | t-test, p < | 0.001 | 0.001 | 0.001 | 0.001 |
|  | Dataset 2<br>(~ 5,000 epochs,<br>17 participants) | Tonic | -2.53 ± 0.28 | -2.58 ± 0.30 | -2.50 ± 0.28 | -2.40 ± 0.28 |
|  |  | Phasic | -2.57 ± 0.26 | -2.59 ± 0.27 | -2.49 ± 0.28 | -2.36 ± 0.26 |
|  |  | Effect size, d | 0.114 | 0.013 | -0.050 | -0.119 |
|  |  | t-test, p | < 0.001 | 0.494 | 0.010 | < 0.001 |
|  | Dataset 3<br>(~ 6,160 epochs,<br>18 participants) | Tonic | -2.20 ± 0.36 | -2.22 ± 0.36 | -2.13 ± 0.31 | -2.14 ± 0.30 |
|  |  | Phasic | -2.26 ± 0.29 | -2.24 ± 0.31 | -2.13 ± 0.30 | -2.08 ± 0.27 |
|  |  | Effect size, d | 0.188 | 0.040 | -0.009 | -0.214 |
|  |  | t-test, p | < 0.001 | 0.025 | 0.615 | < 0.001 |
| High-band (30 – 48Hz) | Dataset 3 | Tonic | -3.89 ± 0.73 | -3.94 ± 0.73 | -3.79 ± 0.68 | -3.78 ± 0.66 |
|  |  | Phasic | -3.59 ± 0.69 | -3.40 ± 0.69 | -3.32 ± 0.64 | -3.49 ± 0.62 |
|  |  | Effect size, d | -0.417 | -0.750 | -0.723 | -0.456 |
|  |  | t-test, p < | 0.001 | 0.001 | 0.001 | 0.001 |

In addition, in Dataset 3 (the only dataset where the high-band (30 – 48Hz) analysis was possible to perform, n = 18), the phasic state showed flatter (more positive) high-band slopes compared to the tonic state over all areas with moderate effect sizes. The repeated-measures ANOVA revealed a significant effect of the area (F(15,3) = 12), state (F(17,1) = 194) as well as the state-area interaction (F(15,3) = 8; all p < 0.001). The ROC analysis revealed that the high-band slopes measured over the central and parietal areas showed the highest discriminative ability (AUC ≥ 0.96 ± 0.015, p < 0.001) with sensitivity and specificity as high as 0.91 and 0.80, respectively.

### Gradual approach: Between-episode analysis

Beside the binary REM sleep categorization described above, we also used a more gradual approach to assess REM sleep heterogeneity by quantifying EM amplitudes. Here, we performed correlations at the between-episode (this section) and within-episode (next section) levels.

For between-episode analysis, we pooled together the epochs with EM events (defined as EOG amplitude > 50μV) that were longer than one minute. This resulted in 7,554 four-second epochs from 20 participants (Dataset 3). We found that EM amplitudes correlated negatively with the low-band aperiodic slopes (calculated from the EEG of the corresponding epochs) over the frontal, central and parietal areas and positively with the low-band aperiodic slopes over the occipital area and with the high-band aperiodic slopes over all areas (upper row in Fig.2 A).

**Figure 2.**
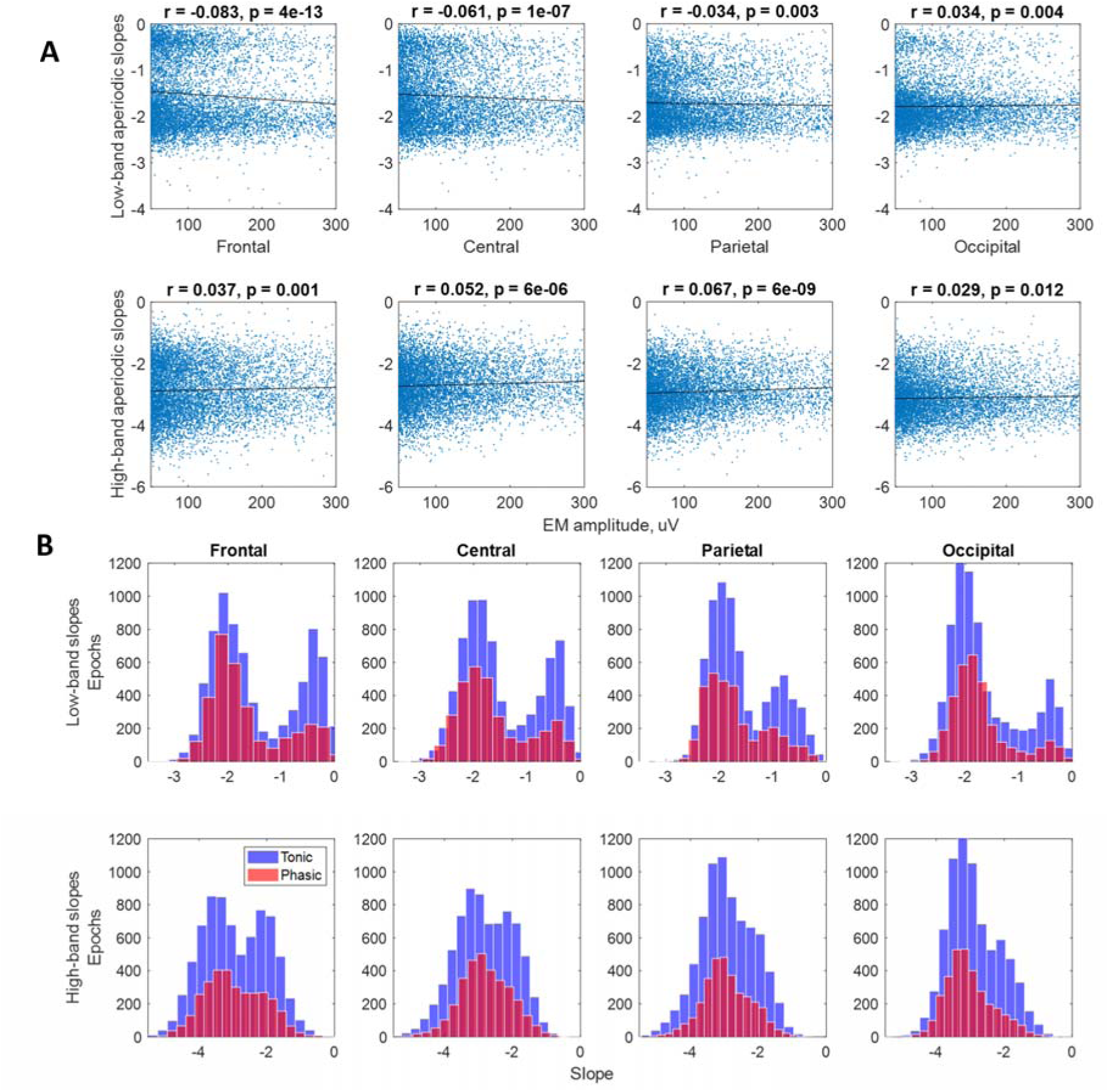
Continuous approach. Aperiodic slopes in the low (2 – 30Hz, upper row) and high (30 – 48Hz, lower row) bands over frontal, central, parietal and occipital electrodes from the continuous data from Dataset 3 pooled for all participants (20 participants, 7554 epochs). A: Aperiodic slopes and EM amplitudes. Correlations between EM amplitudes and aperiodic slopes where each dot represents a 4s epoch, r – Spearman’s correlation coefficient, EM – eye movements. **B: Slope frequency distribution.** Aperiodic slopes show bimodal distribution: e.g., the frontal low-band slope distribution has a major mode centered at −2.1 (the slope value typical for REM sleep), and a minor mode centered at −0.4 (the slope values typical fo wakefulness), which probably reflects sleep arousals naturally weaved into the texture of REM sleep. X-axis exhibits slope values.

Interestingly, we observed that aperiodic slopes derived from the continuous data distributed bimodally (Fig.2 B). For example, the frontal low-band slope distribution had a major mode (i.e., the larger local peak) centered at −2.1 (the slope value typical for REM sleep) and the minor mode centered at −0.4 (a value typical for the wake, see also Discussion, section *Spectral exponent range interpretation*). For comparison, in Supplementary Material 1, we also show slope distribution of the categorical data from the same dataset, which is unimodal (Fig.S1 B). Slope distribution for each participant individually is presented in Fig.S1 C.

Since the data was carefully cleaned for artifacts and only the epochs that fully satisfy the AASM criteria for REM sleep were considered, we presume that the minor mode reflects sleep arousals. Spontaneous sleep arousals are authentic elements of undisturbed sleep microstructure defined as transient accelerations in sleep EEG rhythms that last for ≥ 3 seconds (AASM). Given that arousals are weaved into the texture of REM sleep (Halász et al., 2004), we did not exclude them from the analysis.

The finding on the bimodal distribution of aperiodic slopes, however, raised a question of whether the correlations between aperiodic slopes and EM amplitudes a consequence of this bimodal distribution are. To control for the built-in heterogeneity of the continuous REM sleep data, we further stratified all slopes into two groups that would be distributed unimodally. The cutting points were chosen both theoretically and empirically. The first cutting point was equal to −1, where slopes < −1 were considered typical for sleep and sleepiness (including wake after sleep onset) and slopes > −1 were considered typical for evident wakefulness (Bódizs et al., 2024). The second cutting points were data-driven, i.e., we cut the data at the antimodes −1.3 and −1.5.

We found that the direction, topography and significance of the correlations between EM amplitudes and slopes < −1 remained the same as for all slopes (i.e., before the stratification). Namely, EM amplitudes correlated negatively with the low-band aperiodic slopes over the frontal, central and parietal areas and positively with the low-band aperiodic slopes over the occipital area (Fig.S2, Supplementary Material 1).

### Gradual approach: Within-episode analysis

Next, we assessed temporal relationships between aperiodic activity and EMs within each REM sleep episode using cross-correlations. An example time series of EM amplitudes and aperiodic slopes for a representative REM sleep episode is shown in Fig.3 A. All REM episodes are presented in Supplementary Material 2 and on https://osf.io/zhyf7.

**Figure 3.**
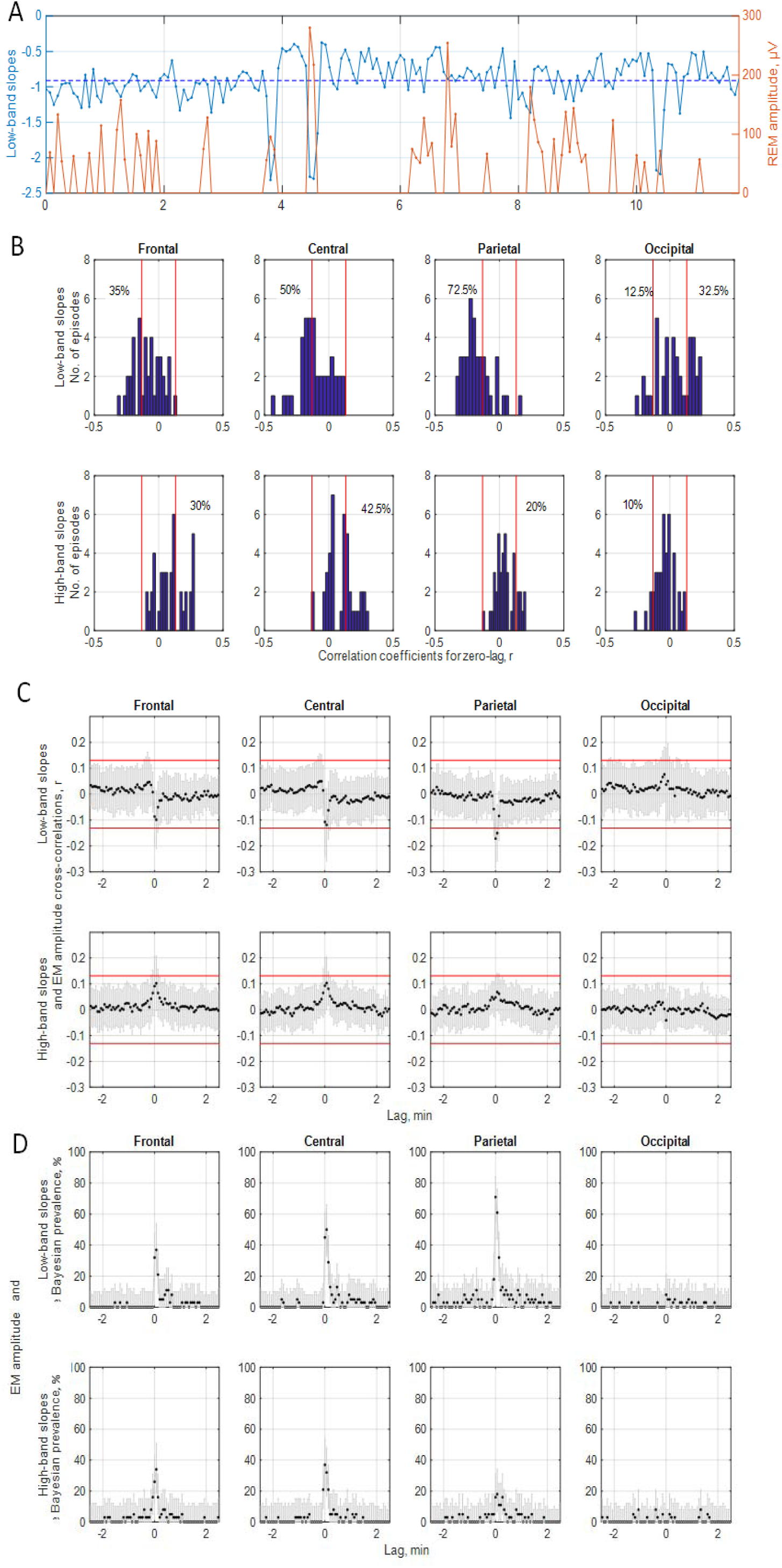
EM amplitudes and aperiodic slopes. **A: Representative time series. Top:** The time series of EM amplitudes and low-band parietal aperiodic slopes calculated for each 4s epoch during a 15.4-minute REM sleep episode. The horizontal dashed blue line shows the slope mean. Higher EM amplitudes are associated with steeper slopes at the time of the event (zero lag, r = −0.31, p < 0.001). **B: Histograms.** Frequency distribution of Spearman coefficients obtained while correlating between the time series of EM amplitudes and aperiodic slopes with the time lag of zero in each REM sleep episode individually (n = 40). The vertical red lines mark the confidence interval of 95%. The absolute values to the left and right of these lines indicate statistically significant correlations. **C: Average coefficients of cross-correlations with standard deviations.** Cross-correlations were calculated between the ranked time series of EM amplitudes and low-band **(top)** or high-band **(bottom)** aperiodic slopes averaged over frontal, central, parietal and occipital electrodes for each REM sleep episode individually. Correlation coefficients were Fisher z-transformed and averaged over all episodes. Negative and positive lags mean that aperiodic slope time series are leading and lagging, respectively. The horizontal red lines mark the confidence interval of 95%, and the absolute values above these lines indicate statistical significance (|r| > 0.1, p < 0.05). The only significant correlation is that for the parietal low-band aperiodic slopes at zero lag (the time of EM event). **D: Bayesian population prevalence of effect sizes.** The Bayesian population prevalence (in %) of the episode-centered effect sizes (black dots), i.e., the most likely value of the population parameter, and the highest posterior density intervals (gray lines), the range within which the true population value lies with the probability level of 96%. Episode-centered effect sizes were obtained by counting the number of significantly negative (for low-band, top) or positive (for high-band, bottom) correlations between time series of aperiodic slopes and EM amplitudes and dividing it by the total number of all correlations (both significant and non-significant, n = 40).

We found that for zero-lags, time series of EM amplitudes and low-band aperiodic slopes correlated negatively over the parietal area (|r| averaged over all episodes > 0.1, p < 0.05, Fig.3 C). At the individual episode level, this correlation was observed in 72.5% of the analyzed episodes (Fig.3 B and D). High-band aperiodic slopes correlated with EM amplitudes in 10 – 42.5% of the episodes only depending on the topographical area (Fig.3 B and D). Table 3 reports the exact proportions, their Bayesian population prevalence and average Spearman correlation coefficients for each topographical area.

**Table 3:** EM amplitudes and aperiodic slopes: within episode correlations.

| Band | 2 – 30Hz |  |  |  | 30 – 48Hz |  |  |  |
| --- | --- | --- | --- | --- | --- | --- | --- | --- |
| Area | F | C | P | O | F | C | P | O |
| REM sleep episodes showing a significant effect, % | 35 | 50 | 72.5 | 12.5 | 30 | 42.5 | 20 | 0 |
| Bayesian estimate prevalence, % | 32 | 47 | 71 | 8 | 26 | 39 | 16 | — |
| Bayesian density interval, % | 17 – 48 | 31 – 64 | 55 – 84 | 0 – 21 | 13 – 43 | 24 – 56 | 5 – 31 | — |
| Averaged correlation coefficient ± std, r | -0.086 ± 0.011 | -0.107 ± 0.128 | **-0.171 ± 0.110** | 0.037 ± 0.140 | 0.091 ± 0.114 | 0.093 ± 0.108 | 0.048 ± 0.078 | -0.041 ± 0.082 |

For time lags ≠ 0, aperiodic activity preceding or following EM events did not cross-correlate with EM amplitudes (|r| averaged over all episodes < 0.1, p > 0.05, Fig.3 C). At the individual episode level, less than 15% of all REM sleep episodes showed significant correlations for time lags ≠ 0 (p < 0.05, Fig.3 D). This indicates that rapid EMs could not be predicted by the magnitude of aperiodic slopes and vice versa.

## Discussion

This study explored how REM sleep states differ with respect to aperiodic neural activity using either binary (phasic-tonic) categorization or gradual quantification of the intensity of oculomotor events to reflect REM sleep heterogeneity. Both approaches showed similar results for instantaneous measurements at the time of an EM event, namely, negative correlations between the EM amplitudes/phasic states and aperiodic slopes. Nevertheless, the temporal analysis within individual REM sleep episodes revealed that EMs could not be predicted by preceding aperiodic activity.

### Slope value meaning

REM sleep is rather heterogeneous with its tonic constituents being depicted as more wake-like or intermediate states between wakefulness and phasic REM sleep concerning several characteristics, including environmental alertness and external information processing (Simor et al., 2020). The current study supports this point of view from a new angle, that of aperiodic neural activity.

Aperiodic activity is best described by the spectral exponent α that reflects the steepness of the decay of the power-law function 1/f^α^ (He, 2014). In the log-log space, exponents are equivalent to slopes, the variable reported in this study. Crucially, specific ranges of spectral exponents define specific features of the underlying time series (Bódizs et al., 2024) as follows:

Exponents recorded during wake tend to be closer to fractional values around −1 (f^−1^ or, equivalently, 1/f) or slightly above as derived from the broadband fitting. An exponent of −1 is a sign of antipersistent dynamics characterized by substantial differences between time scales and negative correlations between successive increments of time series (e.g., amplitude decreases are followed by their increases and *vice versa*). Nevertheless, the nominal exponent values during wake vary substantially and could be as low as −2, especially for wake after sleep onset (Schneider et al., 2022).

The power-law function depicting NREM sleep exhibits fractional exponents between −3 and −2.5 (f^−3^ — f^−2.5^, see Table 3 in Bódizs et al. (2024) for benchmarks). These exponents are the sign of persistent dynamics, where serial increments correlate positively (i.e., increases within the time series are followed by increases and *vice versa*) and overall neural dynamics differ less when observed across different time scales.

REM sleep signals show fractional exponents of around −2 (f^−2^), which is characteristic of the time series with rather independent successive increments. Importantly, α = −2 is seen as a critical value delimiting evident wakefulness and sleep or sleepiness (Bódizs et al., 2024).

In three different datasets of the current study, average exponent/slope values recorded during REM sleep ranged from −2.0 to −2.6 within the fitting band of 2 – 30Hz. This is in line with Schneider et al. (2022) who reported REM sleep exponent values of −2.4 for the fitting band of 2 – 48Hz in young adults. Our study further specified that tonic slopes showed average slopes ranging from −2.1 to −2.5 while phasic slopes presented with values from −2.3 to −2.6 (Table 1, see also Fig.S1 showing histograms). Thus, concerning the spectral exponent range, tonic states are intermediate between phasic REM sleep and wakefulness. This is in coherence with a reported wake-like nature of the tonic state regarding frequency-specific cortical synchronization (Simor et al., 2018) or auditory stimulation effects as quantified by event-related potentials (Takahara et al., 2006).

Moreover, our study revealed that both tonic and phasic aperiodic slopes are distributed bimodally with a major mode centered slightly below −2 and a minor mode centered between −1 and 0 (Fig.2 B). While the major mode values are typical for REM sleep, the minor mode values are more wake-like. We presume that the minor mode reflects transient sleep arousals, the key elements of REM sleep weaved into its microstructure to ensure sleep reversibility and the sleeper’s connection with the external world (Halász et al., 2004). The observed numbers reveal some overlap with Fig.4 in Miskovic et al. (2019) showing that REM-related slopes are distributed from −2 to −1 for the fitting band of 0.5 – 35Hz.

Interestingly, it has been suggested that the 1/f^α^ function with α around −1 represents the “memory” of the EEG signal or local field potentials, i.e., time series’ ability to integrate the effects of their own history (Keshner, 1982). Fig.2 B here reveals that slopes around −1 are more common in tonic than phasic states. Seen in this framework, a longer timescale of the tonic EEG signal could mean that it has stronger integrative properties over both more recent and the distant past. A shorter timescale of the phasic signal may indicate a stronger effect of the recent compared to distant past on actual amplitude values.

Seen in the context of the spectral exponent framework, our findings suggest that the REM sleep signal alternates between states with higher vs lower persistency. Episodes with higher signal persistency (which correspond to an “offline” mode in previous studies) might be needed to protect or preserve sleep (Lina et al., 2019) or promote coordination of neural activity (Montgomery et al., 2008), whereas episodes with higher signal anti-persistency (an “online” mode) might be needed to reinstate to some extent the environmental alertness (Simor et al., 2020).

### Topography of signal persistency and complexity

It has been suggested that time series showing higher anti-persistency are associated with a more disorganized/randomized network that favors environmental responsiveness during wakefulness and possibly reflect high turbulence/fluctuations in neuronal activity (Lina et al., 2019). Interestingly, aperiodic dynamics showing more anti-persistency (flatter aperiodic slopes) are often described as more complex. A recent study showed that tonic states present with higher signal complexity compared to phasic states (Lu et al., 2024). We replicated this finding and further showed that Lempel-Ziv complexity correlated with low-band aperiodic slopes over all areas in all participants (Supplementary Material 1, Fig.S4, Table S1). This is in coherence with previous reports on correlations between aperiodic slopes and LZC during non-REM, REM sleep and rest (Medel et al., 2020; Höhn et al., 2024; Rosenblum et al., 2023b).

Seen in the context of REM sleep states, higher anti-persistency/complexity of the low-band signal recorded during tonic states over frontal, central and parietal areas could be attributed to higher levels of conscious content that require more complex brain activity. This is again in line with the more wake-like and “online” mode of the tonic state (Simor et al., 2020) as well as with the nature of dream reports obtained after tonic periods. Specifically, “tonic” dreams have been characterized by the presence of thinking, recognizing, interpreting, being aware, comparing and/or explaining, overall, being more thought-like than the visual “phasic” dreams (Molinari et al., 1969).

Interestingly, here, low-band aperiodic slopes were flatter over the occipital vs anterior areas during the phasic state only. Moreover, occipital slopes were flatter in phasic compared to tonic states in two out of three datasets, opposite to that measured over more anterior areas. One possible interpretation is that the phasic signal quality differs as a function of a brain area. This resembles the established link between the posterior cortical area-specific decrease in slow-wave activity and the presence of conscious experience in the form of dreams (Siclari et al., 2017). We hypothesize that states showing conscious experience would exhibit flatter aperiodic slopes, the variable that, unfortunately, had not been assessed by Siclari et al. (2017). Intriguingly, in mice, neural activity in the occipital cortical regions (including visual areas) gradually increases during NREM-REM sleep transitions and stays high throughout the entire REM episode (Wang et al., 2022).

Another explanation for the observed topographical gradient follows the suggestion that REM sleep homeostasis might share some mechanisms with NREM sleep (Marzano et al., 2010). REM sleep homeostasis is rather underexplored, yet one study has revealed that it has a similar direction and topography of changes in low-frequency activity with NREM sleep (Marzano et al., 2010). Namely, NREM sleep homeostatic pressure (corresponding to higher signal persistency) is higher in frontal than occipital areas (Cajochen et al., 1999; Bódizs et al., 2024).

The observed topographical gradient might also reflect local asynchronies in sleep regulation and delays in signal propagation, resulting in a gradual appearance of a given REM sleep state. In other words, it is possible that the occipital area has an earlier entry into the tonic state (i.e., local tonic state) than the anterior ones. This assumption is in line with the fact that a given brain state can have temporal and spatial features of other brain states (Wang et al., 2022). Of note, the local nature of REM sleep states presumed above has not been reported in literature yet. For now, we infer it from the generally accepted view on the local control of other sleep features, which becomes global only when a large and widespread number of brain areas are synchronously involved (Peter-Derex et al., 2023). Future studies are needed to explore local and global features of REM sleep states.

### Continuous heterogeneity of REM sleep

Though binary REM sleep categorization is broadly used in literature, one should keep in mind that phasic and tonic states might, alternatively, represent “ends of a continuum” (Bueno-Junior et al., 2023). Another view is that REM sleep is “the sequential occurrence of different activity patterns during prolonged tonic activations punctuated by phasic bursts” (Tononi et al., 2024). An earlier view states that REM sleep is unified in its tonic background characteristics (such as low-voltage mixed frequency EEG and EMG suppression) with superimposed phasic events occurring heterogeneously (Molinari et al., 1969). Bueno-Junior et al. (2023) understand REM sleep as a heterogeneous structure where some signals show binary states (e.g., facial and oculomotor activity in both mice and humans), whereas others show continuous pattern (e.g., theta frequency in mice and respiration rate in humans). Intriguingly, the authors further suggest that binary and continuous features co-occur on an infra-slow timescale (< 0.1 Hz) (Bueno-Junior et al., 2023). Following this, here, besides the tonic-phasic dichotomy, we also assessed REM sleep heterogeneity by quantifying EM amplitudes.

The findings obtained using this approach confirmed and extended those obtained using the categorical one for the instantaneous measurements. For example, steeper low-band parietal aperiodic slopes at the time of an EM event were associated with higher EM amplitudes in > 70% of the analyzed REM sleep episodes. At the same time, the analysis of temporal, intra-REM sleep episode relationships between aperiodic and oculomotor activities revealed that EMs could not be predicted by preceding aperiodic activity or vice versa.

A possible interpretation of our findings is that aperiodic and oculomotor activities are regulated by a common (for example, brainstem) pacemaker. Notably, rapid EMs coincide with the ponto-geniculo-occipital circuitry, which could not be measured in humans with non-invasive techniques (Simor et al., 2020; however, see Wehrle et al., 2005). Our findings, therefore, suggest that aperiodic slopes can serve as a reliable biomarker (possibly even the cortical signature indicating subcortical activity) able to differentiate between phasic and tonic states of REM sleep, however, without an ability for temporal predictions.

The temporal aspects of the phasic activity of REM sleep, its neural correlates, and functional significance are far from being established. Among others, it has been suggested that EMs represent time points at which neural activity and associated visual-like processing is updated, possibly reflecting a change of the visual imagery in dreams (Andrillon et al., 2015). Likewise, EMs have been linked to consolidation of procedural memory involving novel cognitive strategies and problem-solving skills, where EMs correlate with theta and sensorimotor rhythms (van den Berg et al., 2023). Another view is that EM density reflects neural excitation as discussed in more detail in the next section.

### Excitation-to-inhibition balance and aperiodic activity

Besides the low-band (2 – 30Hz) aperiodic slopes, here, we also measured the high-band (30 – 48Hz) ones, inspired by the theory stating that they might reflect the balance between excitatory and inhibitory neural currents (Gao et al., 2017), which in turn defines a specific arousal state and a conscious experience of an organism. Seen in the light of this theory, our findings on flatter high-band aperiodic slopes during phasic vs tonic states suggest that the phasic state is associated with a relative shift towards neural excitation. This finding is also in line with the known increase in gamma-band power during the phasic state (Simor et al., 2019). Increases in both high-band aperiodic activity and gamma power seen during phasic states probably have the same neural origin, reflecting wakefulness-like increases in cortical excitability. This may produce short bursts of consciousness, resulting in dreams with vivid visual imagery and high emotional and perceptual content (Simor et al., 2020; Usami et al., 2017).

Our findings showing that higher EM amplitudes coincide with flatter high-band aperiodic slopes (that reflect a shift towards neural excitation) are also in line with the view on EM density as a reflection of the build-up of a pressure to awaken across a night. Indeed, it has been shown that EM density increases across sleep cycles (while sleep pressure, need and depth decrease) and is higher at the circadian time of higher arousal (van den Berg et al., 2023).

### Limitations, strengths and conclusions

The major limitation of this study is its cross-sectional observational design and correlational approach, and thus an inability to shed light on the mechanism underlying the observed effects. Likewise, we analyzed the data of healthy young participants in a restricted age range (mean: 21.7 ± 1.4 years) and it is unclear to what extent our results generalize to the older population.

The strengths of this study include its sample size, scripts and data sharing and self-replication in three independent datasets, affirming the overall robustness of the findings.

In conclusion, our study demonstrates that aperiodic neural activity can serve as a neural correlate and a reliable marker able to differentiate between phasic and tonic constituents of REM sleep and reflects the intensity of phasic oculomotor events, providing further evidence on the heterogeneity of REM sleep.

## Contributors

YR wrote the manuscript and integrated the data and its analysis. All authors analyzed, discussed and interpreted the data, contributed to, reviewed, revised, and approved the final draft of the paper. All authors had final responsibility for the decision to submit for publication.

## Data sharing

For the categorical part of the study, the EEG data is open-access under https://osf.io/2vptx and https://osf.io/9k5hb. Aperiodic slopes for each epoch, for each participant for each dataset as well as EM amplitudes for Dataset 3 are reported in Supplementary Excel File and under https://osf.io/zhyf7.

## Acknowledgements

MD is supported by the Vidi fellowship by the Dutch Research Council (NWO), RB is supported by the Culture and Innovation Ministry of Hungary (TKP2021-EGA-25, TKP2021-NKTA-47).

