## Supplementary Material 1 for "Aperiodic neural activity distinguishes between phasic and tonic REM sleep"

* - equal contribution.

### Supplementary Material 1

### Frequency distribution of aperiodic slopes

A

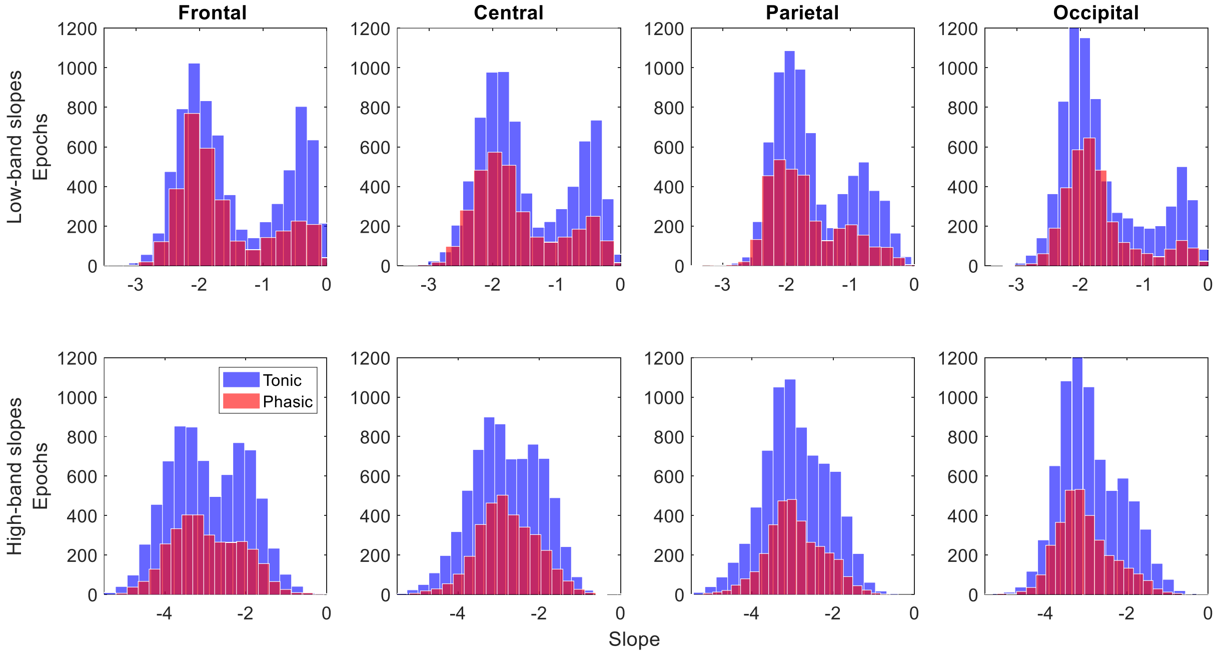

B

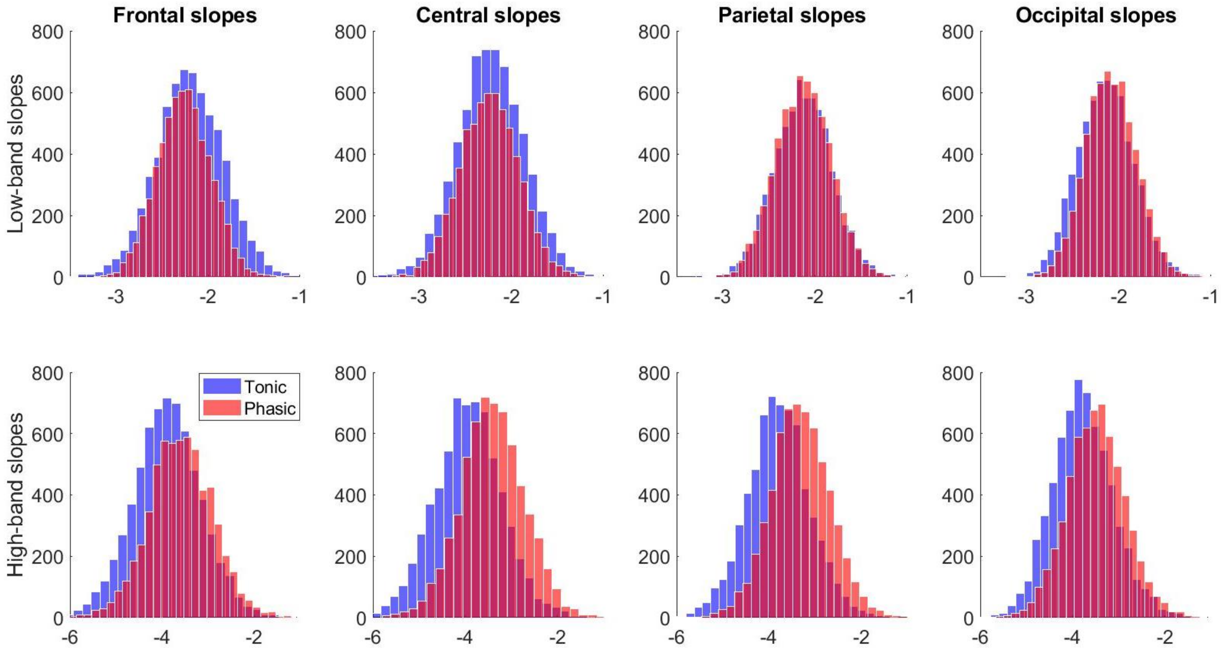

C

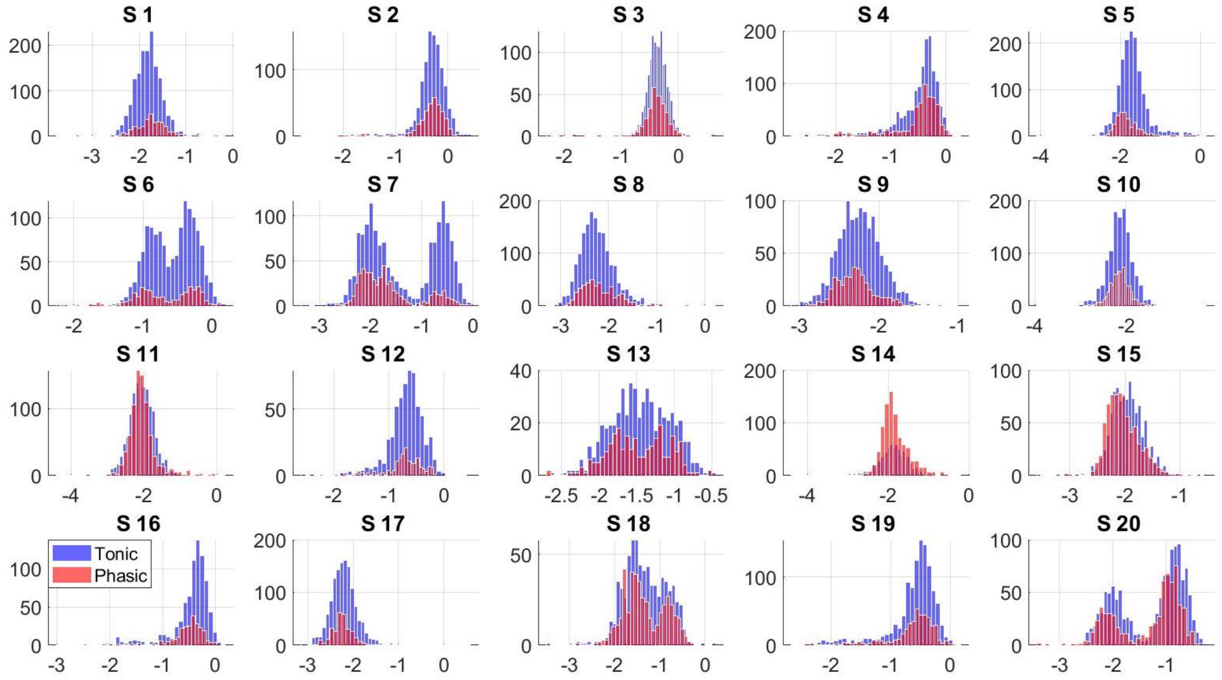

**Supplemental Figure 1. Slope frequency distribution. A – B:** Distribution of aperiodic slopes in the low (2 – 30 Hz, upper row) and high (30 – 48Hz, lower row) bands over frontal, central, parietal and occipital electrodes from the continuous **(A,** the same as Fig.2B**)** and categorical **(B)** data from Dataset 3 pooled for all participants. Continuous data shows bimodal while categorical data shows unimodal distribution. Frontal low-band slope distribution has a major mode centered at -2.1 (the slope values typical for REM sleep), and a minor mode centered at -0.4 (the slope values typical for wakefulness), which probably reflects sleep arousals naturally weaved into the texture of REM sleep. **C:** Distribution of the low-band frontal slopes of the continuous data for each participant individually. X-axis exhibits low-band frontal slope values.

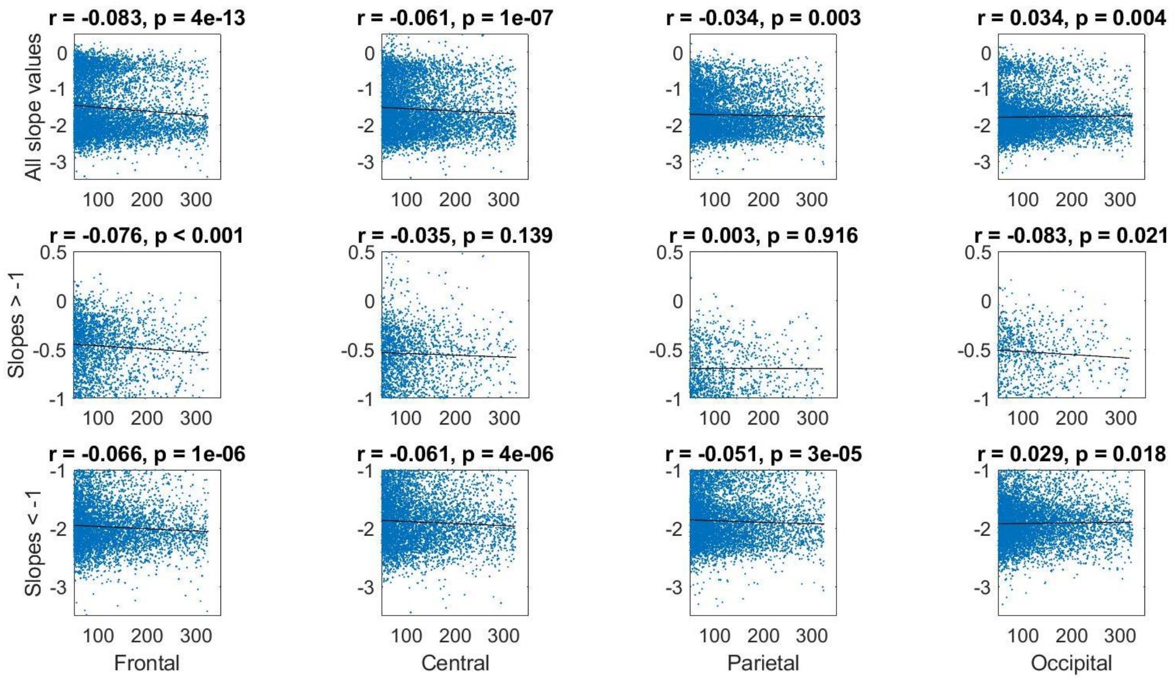

**Supplemental Figure 2. Aperiodic slopes and EM amplitudes.** Correlations between the low-band aperiodic slopes for each topographical area and EM amplitudes are shown separately for all slopes (first row, bimodal distribution), slopes > -1 (values typical for the wake, second row, almost unimodal distribution) and slopes < -1 (values typical for sleep, third row, almost unimodal distribution). Each dot represents a four-second epoch from the continuous data of Dataset 3 (20 participants, 7554 epochs), x-axis exhibits the EM amplitudes in µV, r – Spearman's correlation coefficient, EM – eye movements.

### Oscillations

To replicate previous findings, we also calculated the oscillatory power component by subtracting the aperiodic component from the total power. Then, the power was averaged over delta (2–4Hz), theta (4–8Hz), alpha (8–12Hz), and beta (12–30Hz) frequency bands.

In line with the literature, we found that the phasic state shows lower oscillatory power in the 12–30Hz band (beta) compared to the tonic state with a large effect size ($p < 0.001$. Cohen's $d = -1.1$). Oscillatory power in the rest of the frequency bands was comparable.

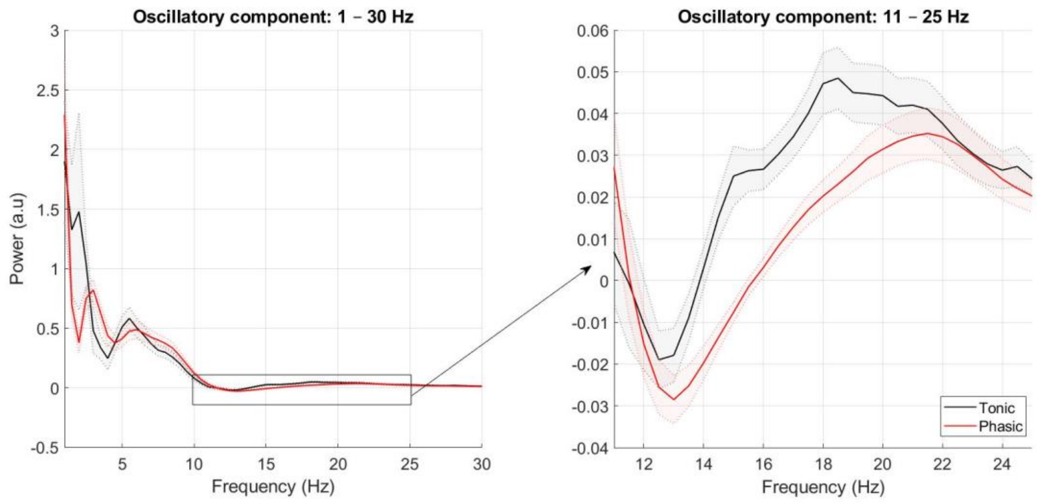

**Supplemental Figure 3. Oscillatory frontal power components.** Spectral power was calculated in the 1 – 48 Hz band over frontal electrodes, differentiated to its aperiodic and oscillatory components and plotted as a function of frequency for tonic (black) and phasic (red) states of REM sleep. Shading indicates standard errors. The phasic state shows lower oscillatory power in the 12 – 25 Hz band (beta) compared to the tonic state.

**Lempel-Ziv complexity**

Lempel-Ziv complexity (LZC, Lempel & Ziv, 1976) is another well-known measure of arousal levels and sleep stages. LZC determines the regularity and compressibility of a signal in the time domain by scanning the symbolic sequences for new patterns and increasing the complexity count every time a new sequence is detected (Lempel & Ziv, 1976). Unpredictable, irregular signals are reflected by higher complexity and vice versa. Higher LZC values indicate more complex data.

LZC was calculated as described in Rosenblum et al. (2023 b) either in the low (2–30 Hz) or high (30–48Hz) bands and averaged over each topographical area separately. All values were transformed to the z-scores as described for aperiodic slopes in the Main text. We found that the phasic state has decreased low-band (2–30Hz) LZC as compared to the tonic state in all topographical areas while high-band (30–48Hz) LZC was comparable in both states.

To test if changes in the LZC can be explained by changes in the EEG power, we employed the normalization procedure as in Rosenblum et al. (2023 b) and found that all differences disappeared after phase randomization normalization.

In addition, we correlated LZC with aperiodic slopes within each participant and found that low-band – but not high-band – slopes and raw LZC correlated positively over all areas in all participants ($r = 0.41 – 0.60$, $p < 0.001$, 200 – 600 epochs per participant per state, Supplemental Table 1). This is in line with previous reports on correlations between aperiodic slopes and LZC during non-REM, REM sleep and rest (Medel et al., 2020; Höhn et al., 2024; Rosenblum et al., 2023 b).

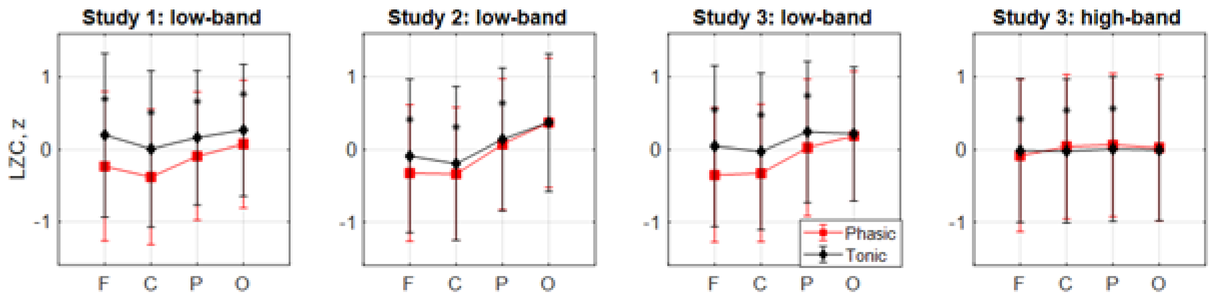

**Supplemental Figure 4. LZC.** Raw LZCs in the 2 – 30 Hz for Datasets 1 – 3 **(from left to right)** and in the 30 – 48 Hz band for Dataset 3 **(right)** were z-normalized and averaged over the phasic and tonic states of REM sleep over each topographical area separately. The tonic state (black squares) shows higher low-band LZC compared to the phasic state (red diamonds) with the anterior-posterior gradient where the largest differences are observed over the frontal area. * – significant difference between tonic and phasic states, F – frontal, C – central, P – parietal, O – occipital, LZC – Lempel-Ziv complexity.

**Supplemental Table 1: Lempel-Ziv complexity**

| Frequency | Dataset | Area | Frontal | Central | Parietal | Occipital |
|---|---|---|---|---|---|---|
| Low-band (2 – 30Hz) | Dataset 1 (~ 2,040 epochs, 20 participants) | Tonic, z | 0.20 | 0.01 | 0.16 | 0.27 |
| | | Phasic, z | -0.23 | -0.38 | -0.09 | 0.07 |
| | | Effect size, d | 0.40 | 0.38 | 0.28 | 0.21 |
| | | t-test, p < | 0.001 | 0.001 | 0.001 | 0.001 |
| | Dataset 2 (~ 5,000 epochs, 17 participants) | Tonic, z | -0.09 | -0.19 | 0.14 | 0.37 |
| | | Phasic, z | -0.33 | -0.34 | 0.07 | 0.37 |
| | | Effect size, d | 0.24 | 0.14 | 0.07 | 0.00 |
| | | t-test, p < | 0.001 | 0.001 | 0.001 | 0.896 |
| | Dataset 3 (~ 6,160 epochs, 18 participants) | Tonic, z | 0.04 | -0.03 | 0.24 | 0.21 |
| | | Phasic, z | -0.35 | -0.33 | 0.03 | 0.18 |

| | | | | | | |
|---|---|---|---|---|---|---|
| | | Effect size, d | 0.39 | 0.30 | 0.22 | 0.03 |
| | | t-test, p < | 0.001 | 0.001 | 0.001 | 0.060 |
| | Merged Dataset | ROC, AUC | **0.81** | **0.83** | **0.78** | 0.60 |
| | | ROC, p < | 0.001 | 0.001 | 0.001 | 0.084 |
| | Correlations between LZC and aperiodic slopes (all p < 0.05) | Phasic, r | 0.581 | 0.507 | 0.425 | 0.477 |
| | | Tonic, r | 0.599 | 0.458 | 0.470 | 0.410 |
| High-band (30 – 48Hz) | Dataset 3 | Tonic | -0.02 | -0.02 | 0.01 | -0.01 |
| | | Phasic | -0.08 | 0.04 | 0.06 | 0.02 |
| | | Effect size, d | 0.07 | -0.06 | -0.05 | -0.03 |
| | | t-test, p < | 0.001 | 0.001 | 0.003 | 0.127 |

*Bold font – significant differences between tonic and phasic states that passed the correction for multiple comparisons, r > 0.7 are considered as strong correlation scores, values lower than 0.3 are considered as weak, r-values in the range of 0.3 – 0.7 are considered as moderate scores. Correlation coefficients were calculated within each participant (200 – 600 epochs per participant) individually and then averaged over all participants, r – Pearson's correlation coefficient, F – frontal, C – central, P – parietal, O – occipital. AUC – area under the curve, ROC – receiver operating characteristic.*

**Electrocardiography (ECG)**

Given that recently, it has been shown that "cortically" measured aperiodic activity could be (at least partly) attributed to cardiac activity captured by surface electrodes via volume conduction (Schmidt et al., 2023), we also analyzed aperiodic activity over the ECG channel. The procedure was the same as for the EEG channels described in Methods. The ECG signal was recorded as part of the EEG recordings and was available for Datasets 1 and 2.

We found comparable cardiac low-band (2 – 30Hz) aperiodic slopes during tonic and phasic states (Study 1: $-1.84 \pm 0.30$ vs $-1.83 \pm 0.32$; Study 2: $-1.99 \pm 0.44$ vs $-2.01 \pm 0.46$). In addition, to confirm that the aperiodic ECG signal relates to heart rate we calculated the mean heart rate (i.e., 60 s/R-R interval s). The R peaks were detected by the *findpeaks* function and checked through the visual inspection for each 4s epoch. Then, we correlated the aperiodic cardiac slopes with heart rate averaged over each epoch using Pearson's correlations within each participant individually. Significant correlations were observed in about 37.5% of all participants only.
