## Supplementary Material 2 for "Aperiodic neural activity distinguishes between phasic and tonic REM sleep"

**Eye movement (EM) amplitudes and aperiodic slopes for each REM sleep episode** (40 episodes from 19 participants, Dataset 3, continuous data).

**Top:** The time series of EM amplitudes and low-band parietal aperiodic slopes calculated for each 4-second epoch of one continuous REM sleep episode. The horizontal dashed blue line shows the mean value of the slopes. Spearman correlation coefficient r (time lag = 0) and an associated p-value are also reported.

**Bottom:** Cross-correlation between the ranked time series of EM amplitudes and aperiodic slopes shown above with time lags from -5 to +5 min. Negative and positive lags mean that aperiodic slope time series are leading and lagging, respectively. The horizontal red lines mark the confidence interval of 95%, and the absolute values above these lines indicate statistical significance ($|r| > 0.1$, $p < 0.05$).

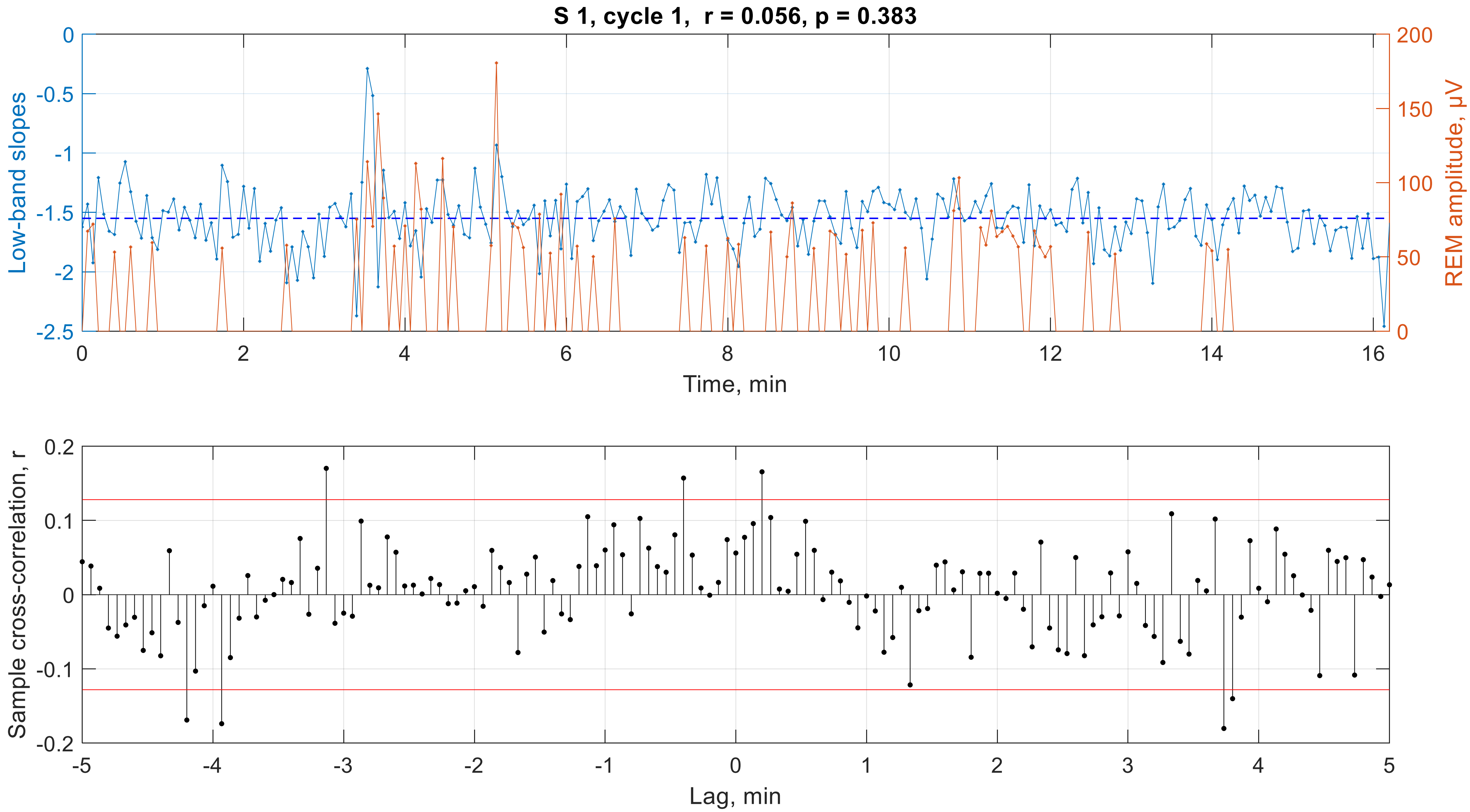

S 1, cycle 1, r = 0.056, p = 0.383
Low-band slopes
REM amplitude, μV
Time, min
0
-0.5
-1
-1.5
-2
-2.5
200
150
100
50
0
2
4
6
8
10
12
14
16
Sample cross-correlation, r
Lag, min
0.2
0.1
-0.1
-0.2
-5
-4
-3
-2
-1
1
3
5

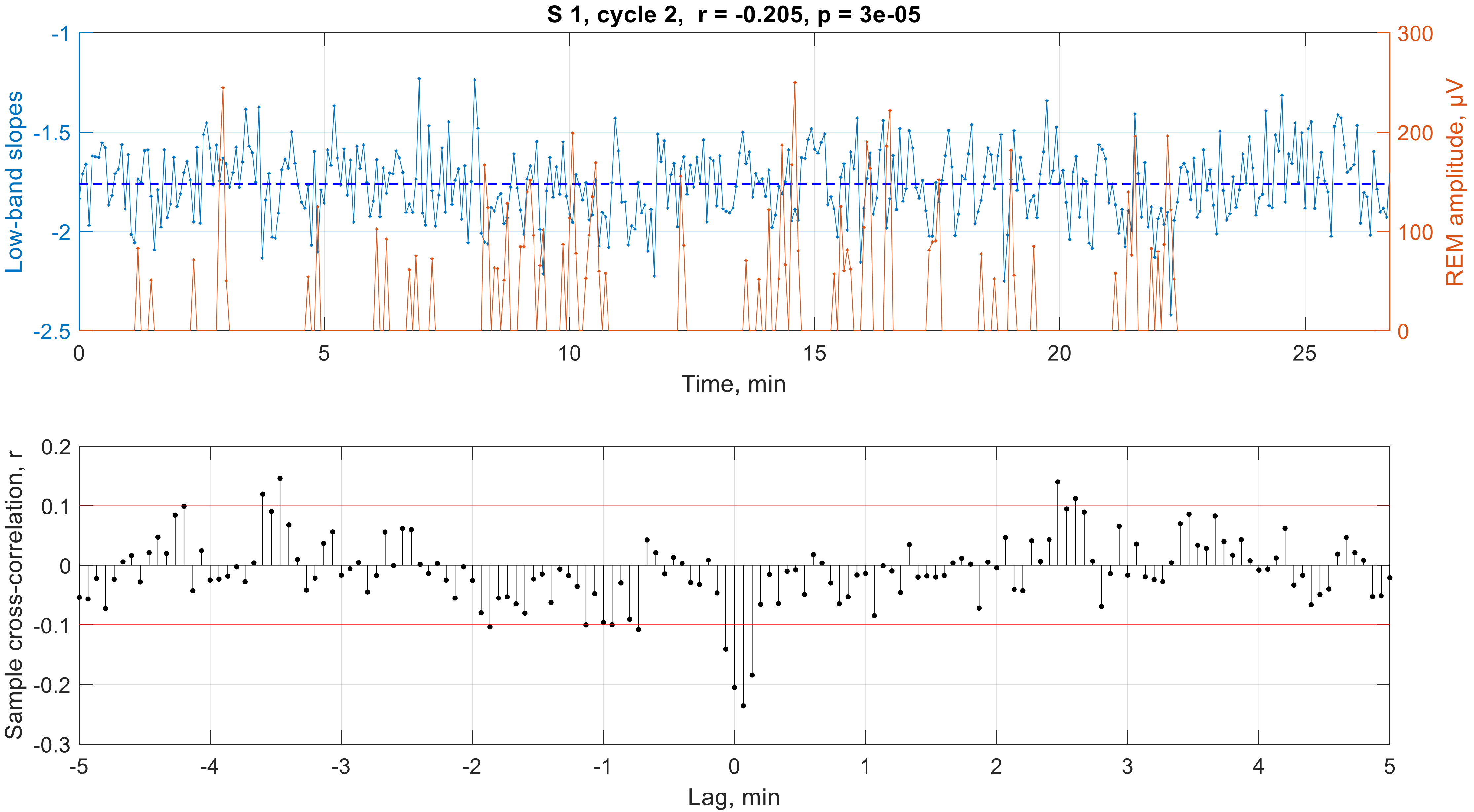

S 1, cycle 2,  r = -0.205, p = 3e-05
Low-band slopes
REM amplitude, µV
Time, min
-1
-1.5
-2
-2.5
300
200
100
0
0
5
10
15
20
25
Sample cross-correlation, r
Lag, min
0.2
0.1
0
-0.1
-0.2
-0.3
-5
-4
-3
-2
-1
0
1
2
3
4
5

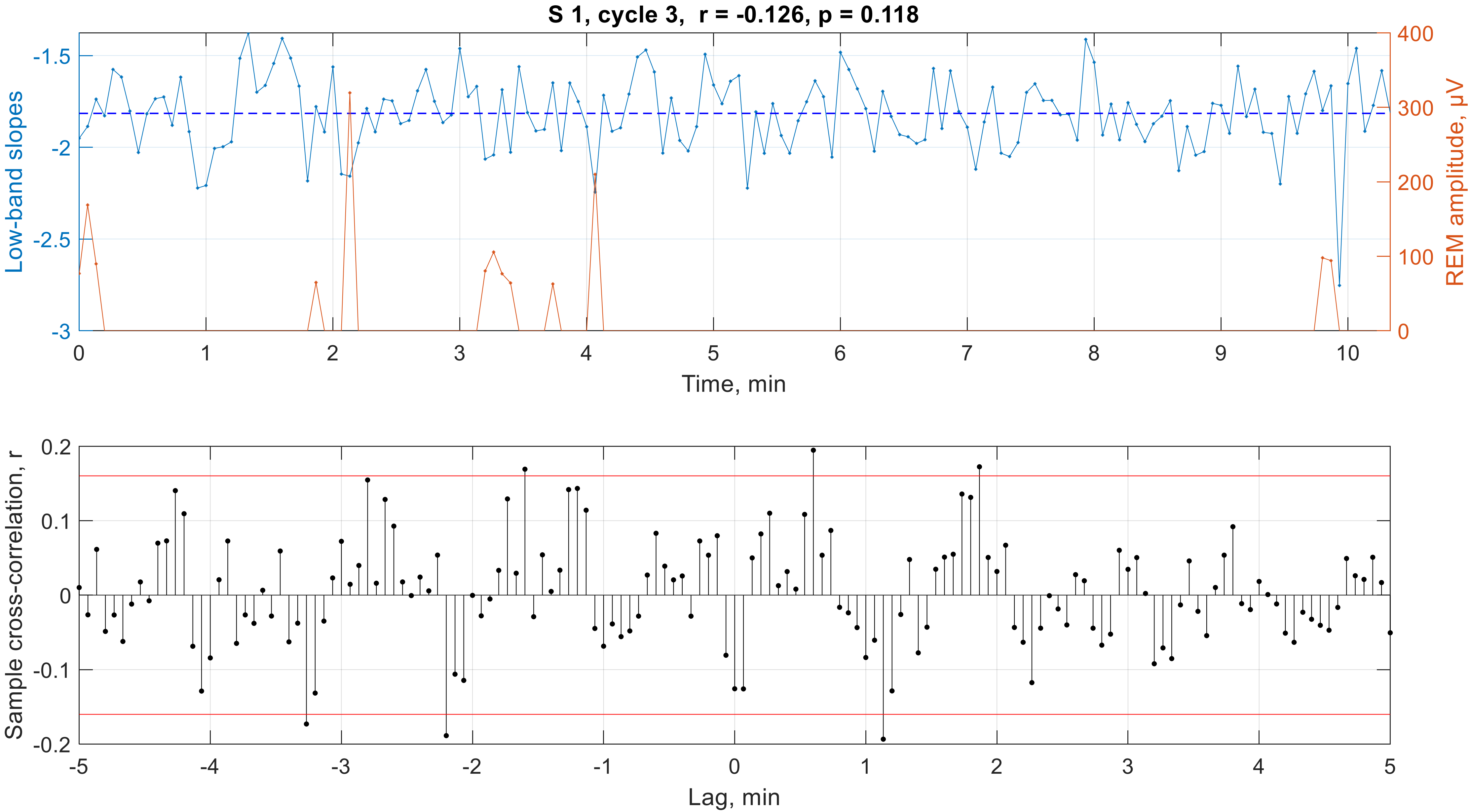

S 1, cycle 3, r = -0.126, p = 0.118
Low-band slopes
REM amplitude, μV
Time, min
Sample cross-correlation, r
Lag, min

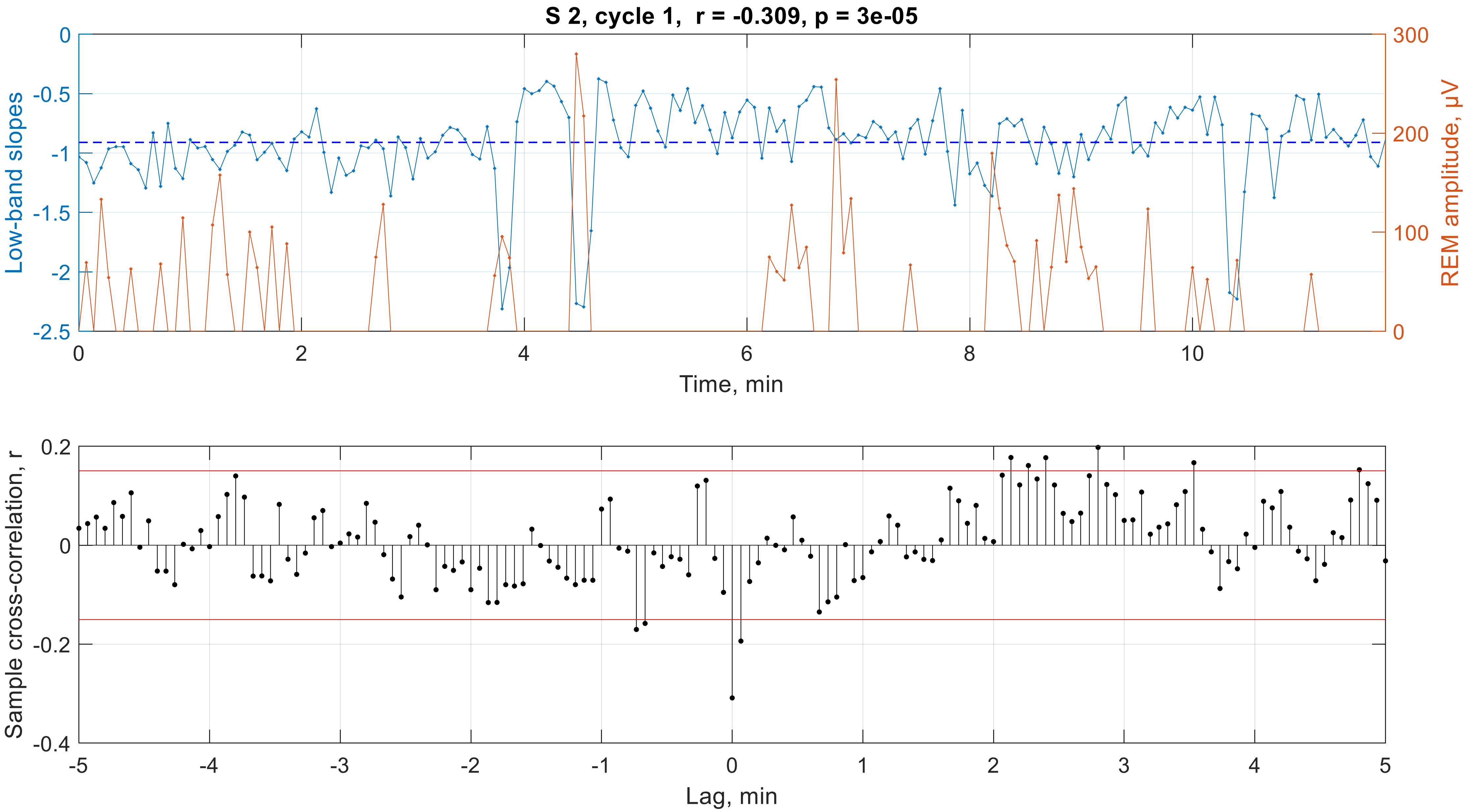

S 2, cycle 1, r = -0.309, p = 3e-05
Low-band slopes
REM amplitude, μV
Time, min
Sample cross-correlation, r
Lag, min

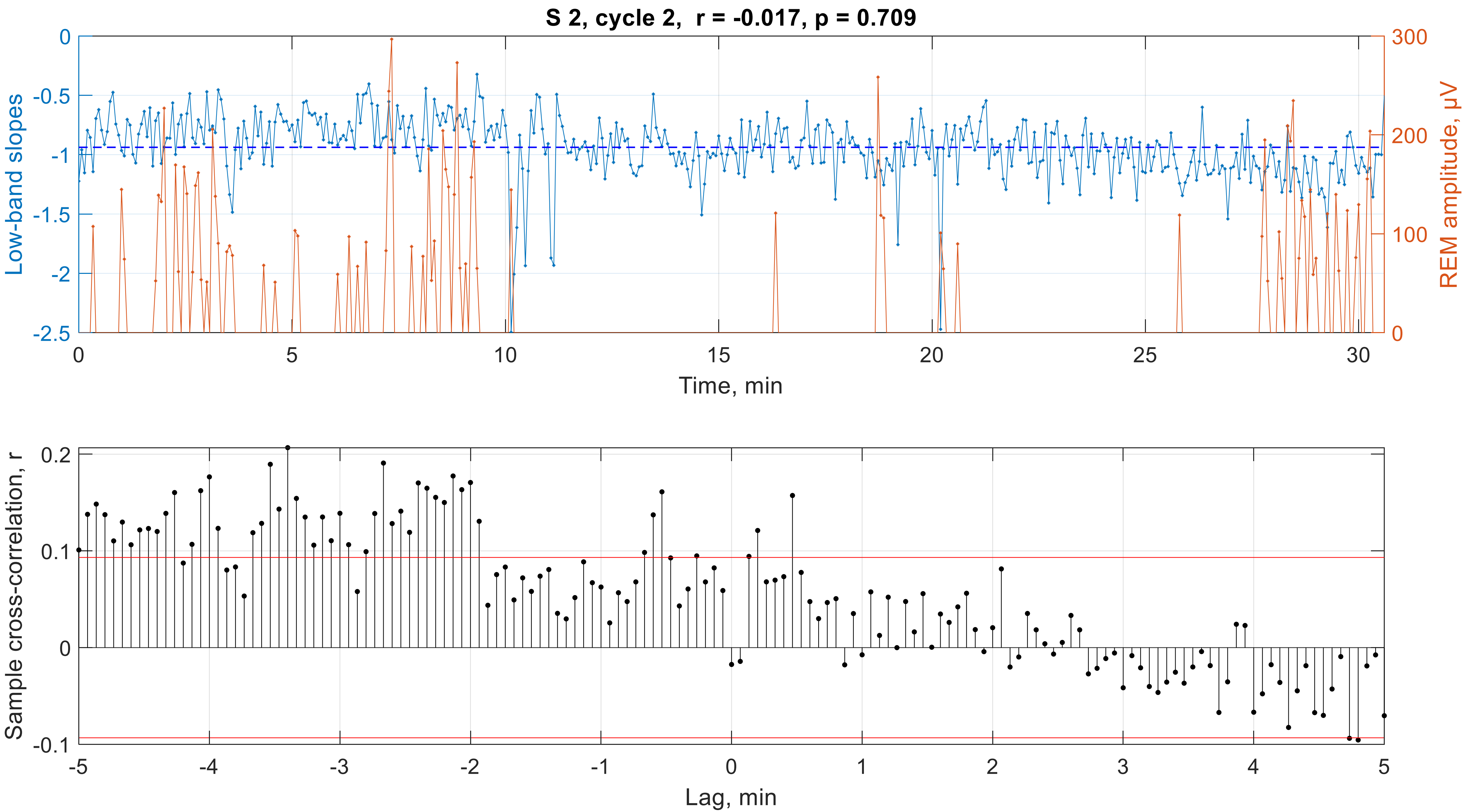

S 2, cycle 2, r = -0.017, p = 0.709
Low-band slopes
REM amplitude, μV
Time, min
Sample cross-correlation, r
Lag, min

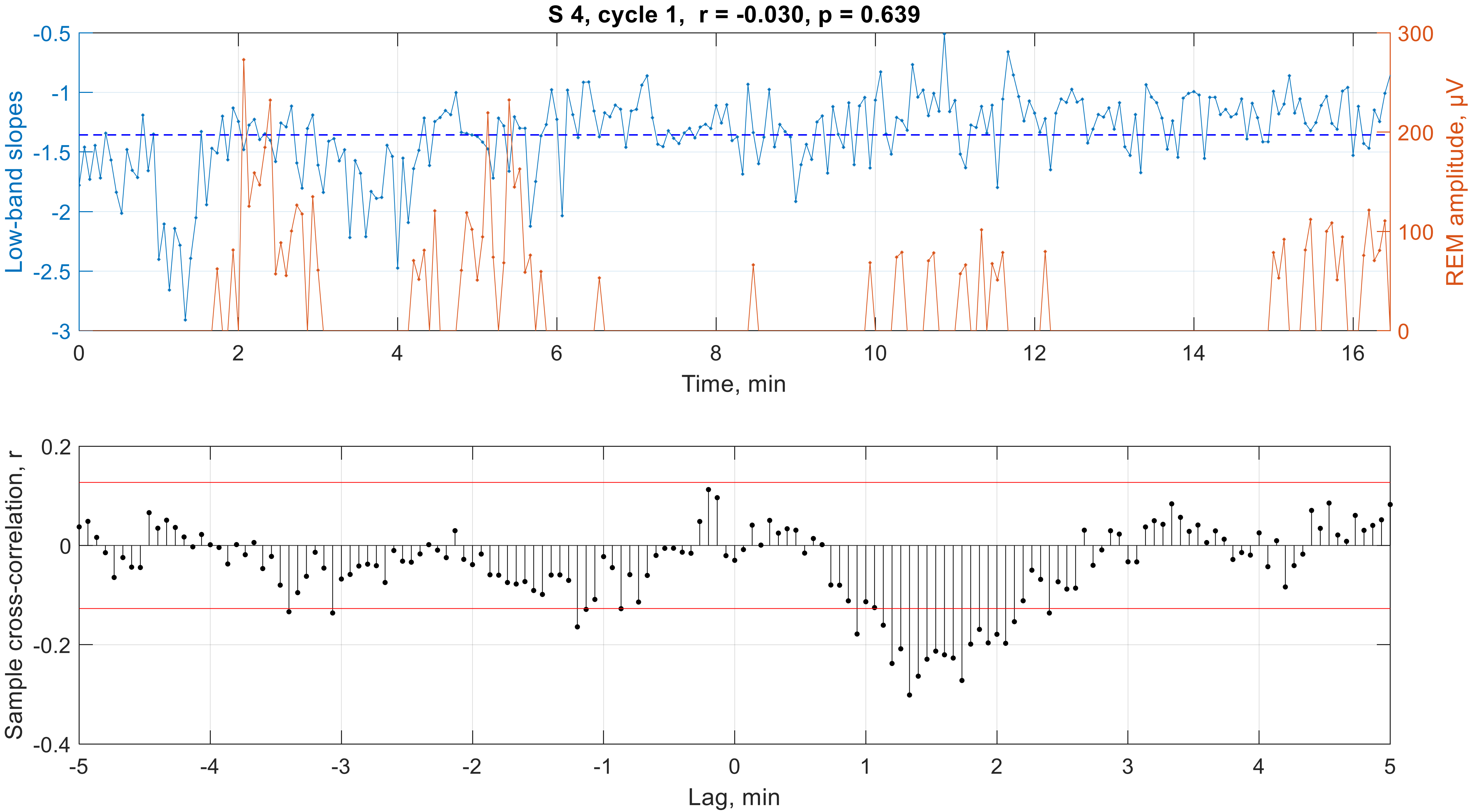

S 4, cycle 1, r = -0.030, p = 0.639
Low-band slopes
REM amplitude, μV
Time, min
Sample cross-correlation, r
Lag, min

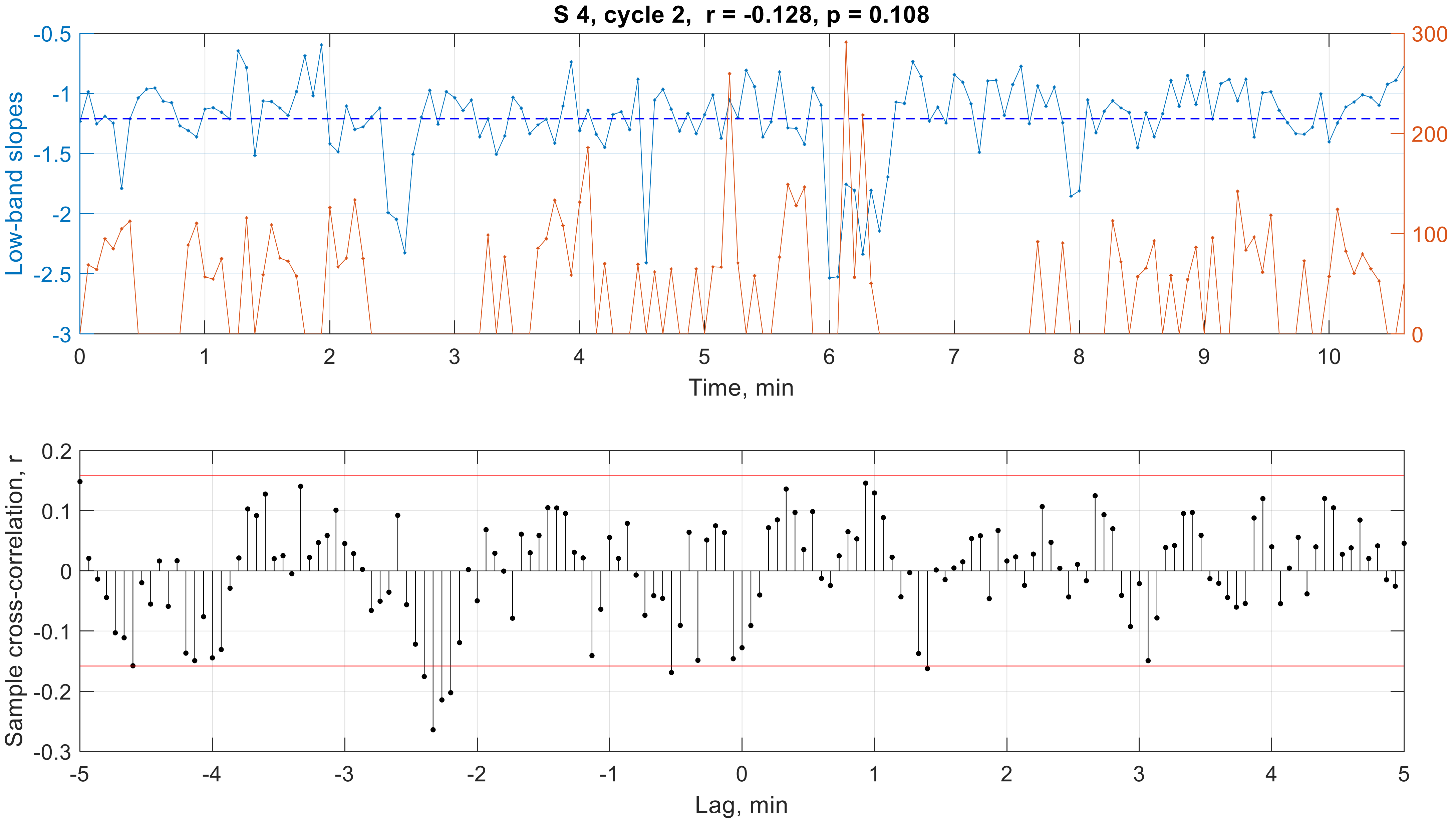

S 4, cycle 2, r = -0.128, p = 0.108
Low-band slopes
REM amplitude, μV
Time, min
Sample cross-correlation, r
Lag, min

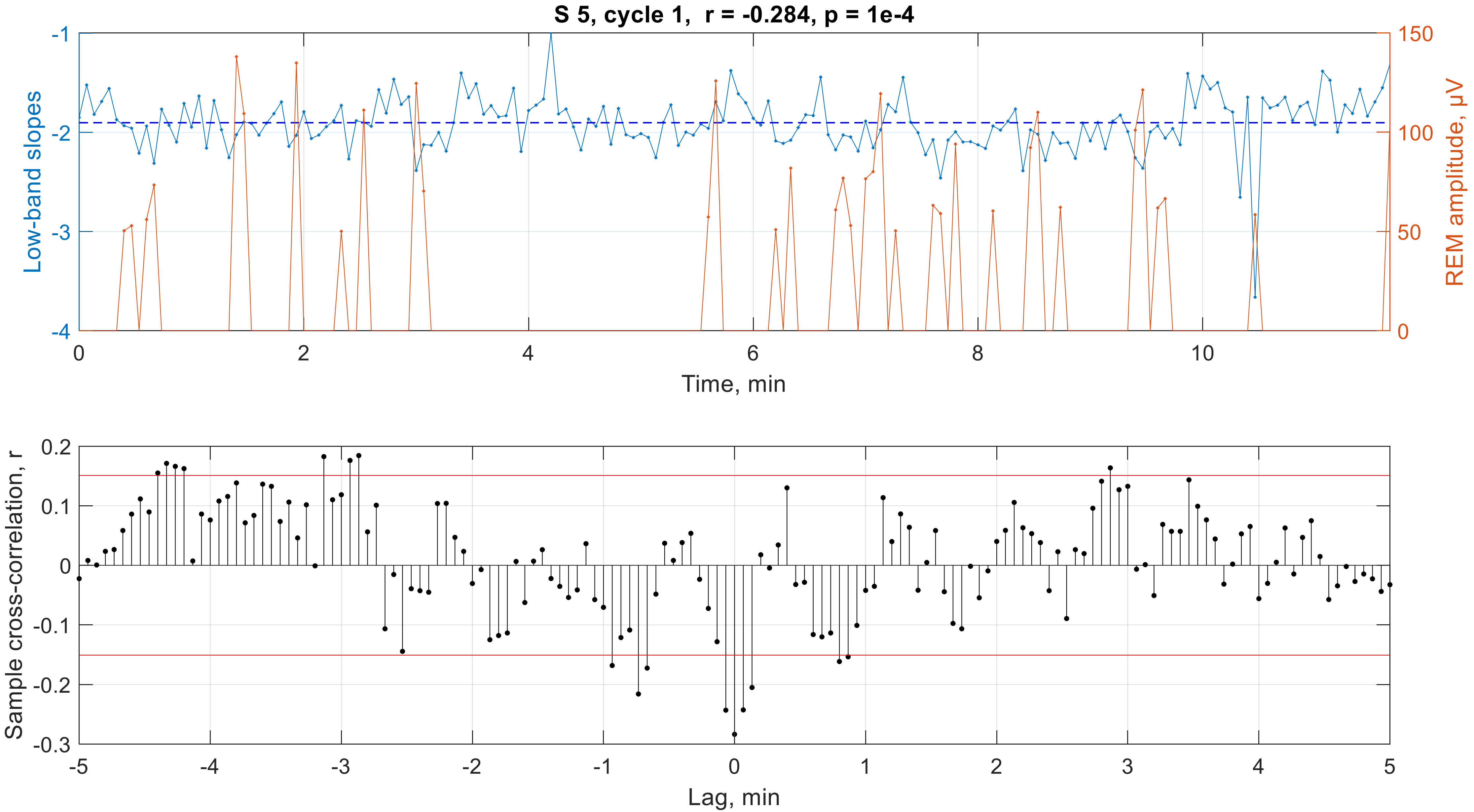
S 5, cycle 1, r = -0.284, p = 1e-4
Low-band slopes
REM amplitude, μV
Time, min
Sample cross-correlation, r
Lag, min

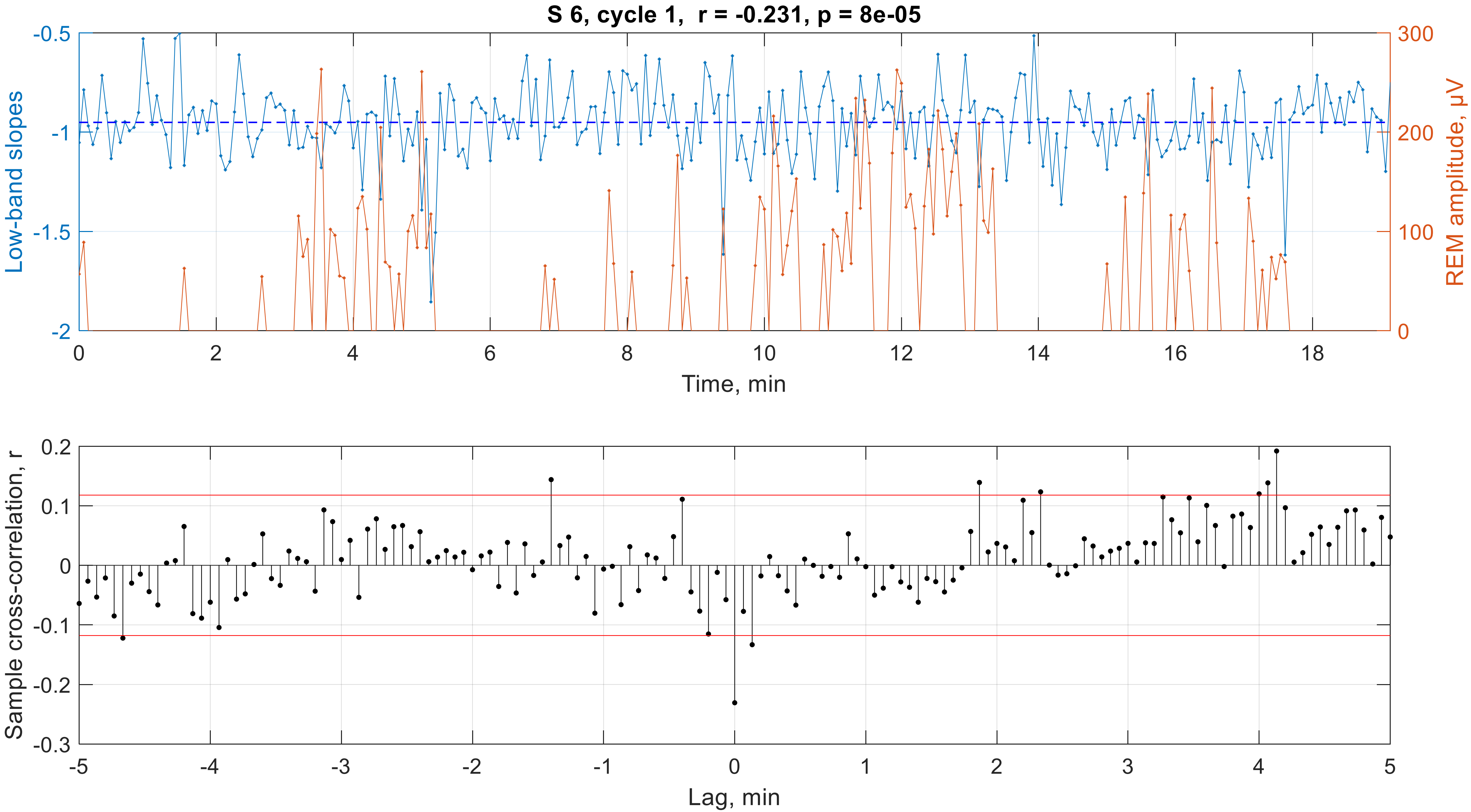

S 6, cycle 1, r = -0.231, p = 8e-05
Low-band slopes
REM amplitude, μV
Time, min
-0.5
-1
-1.5
-2
300
200
100
0
0
2
4
6
8
10
12
14
16
18
Sample cross-correlation, r
Lag, min
0.2
0.1
0
-0.1
-0.2
-0.3
-5
-4
-3
-2
-1
0
1
2
3
4
5

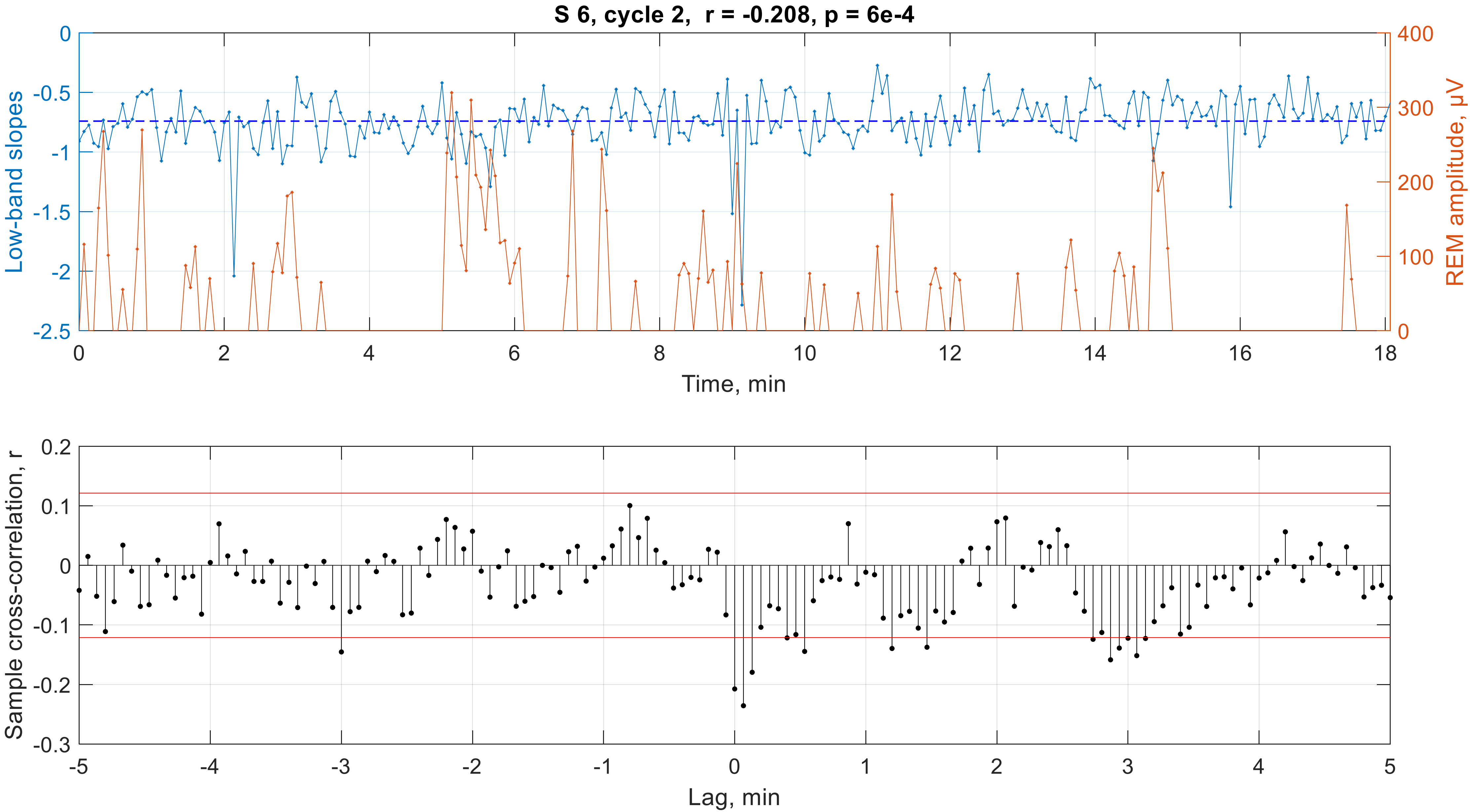

S 6, cycle 2, r = -0.208, p = 6e-4
Low-band slopes
REM amplitude, μV
Time, min
Sample cross-correlation, r
Lag, min

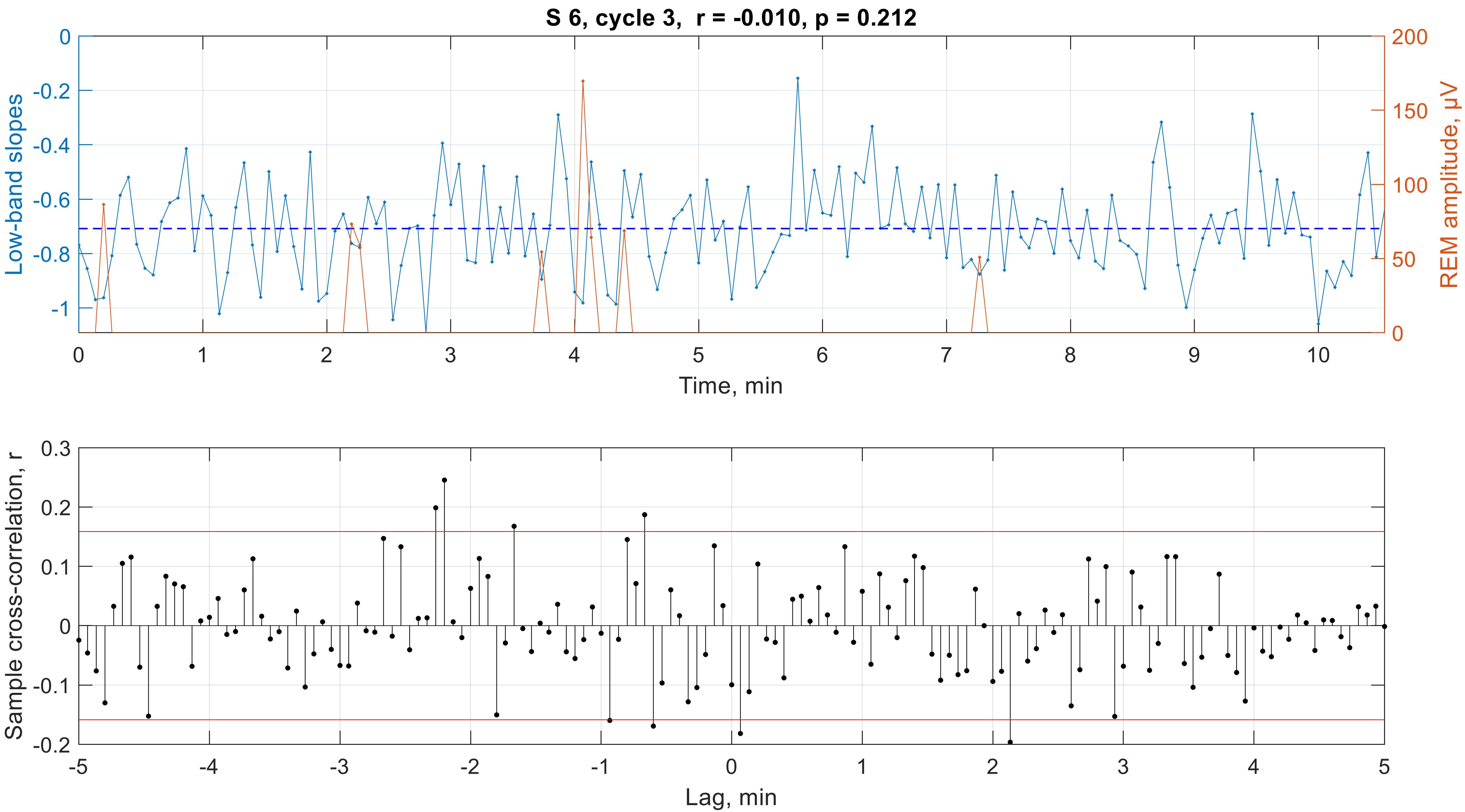

S 6, cycle 3,  r = -0.010, p = 0.212
Low-band slopes
REM amplitude, μV
Time, min
0
-0.2
-0.4
-0.6
-0.8
-1
200
150
100
50
0
0
1
2
3
4
5
6
7
8
9
10
Sample cross-correlation, r
Lag, min
0.3
0.2
0.1
0
-0.1
-0.2
-5
-4
-3
-2
-1
0
1
2
3
4
5

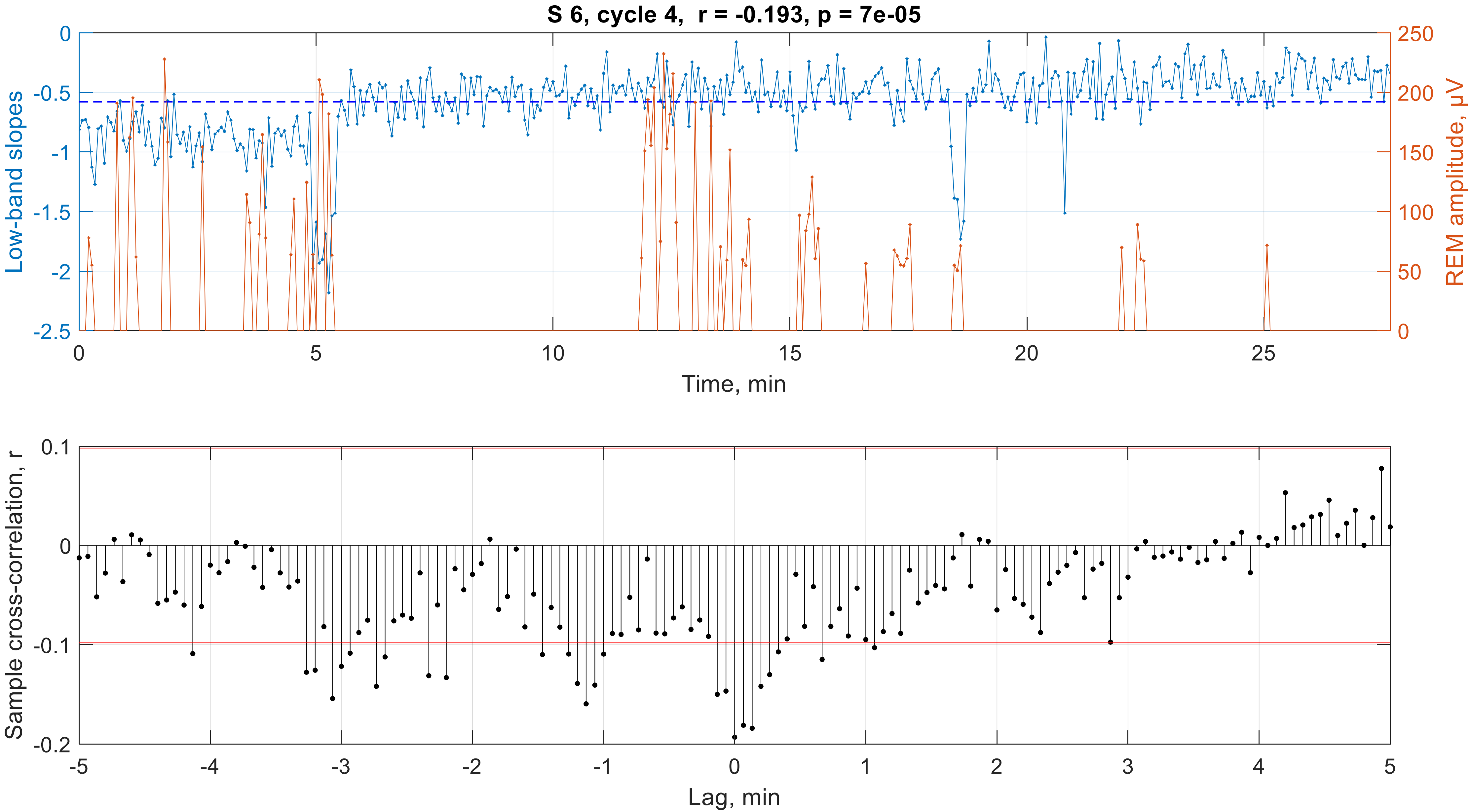
S 6, cycle 4,  r = -0.193, p = 7e-05
Low-band slopes
REM amplitude, μV
Time, min
0
-0.5
-1
-1.5
-2
-2.5
250
200
150
100
50
0
0
5
10
15
20
25
Sample cross-correlation, r
Lag, min
0.1
0
-0.1
-0.2
-5
-4
-3
-2
-1
0
1
2
3
4
5

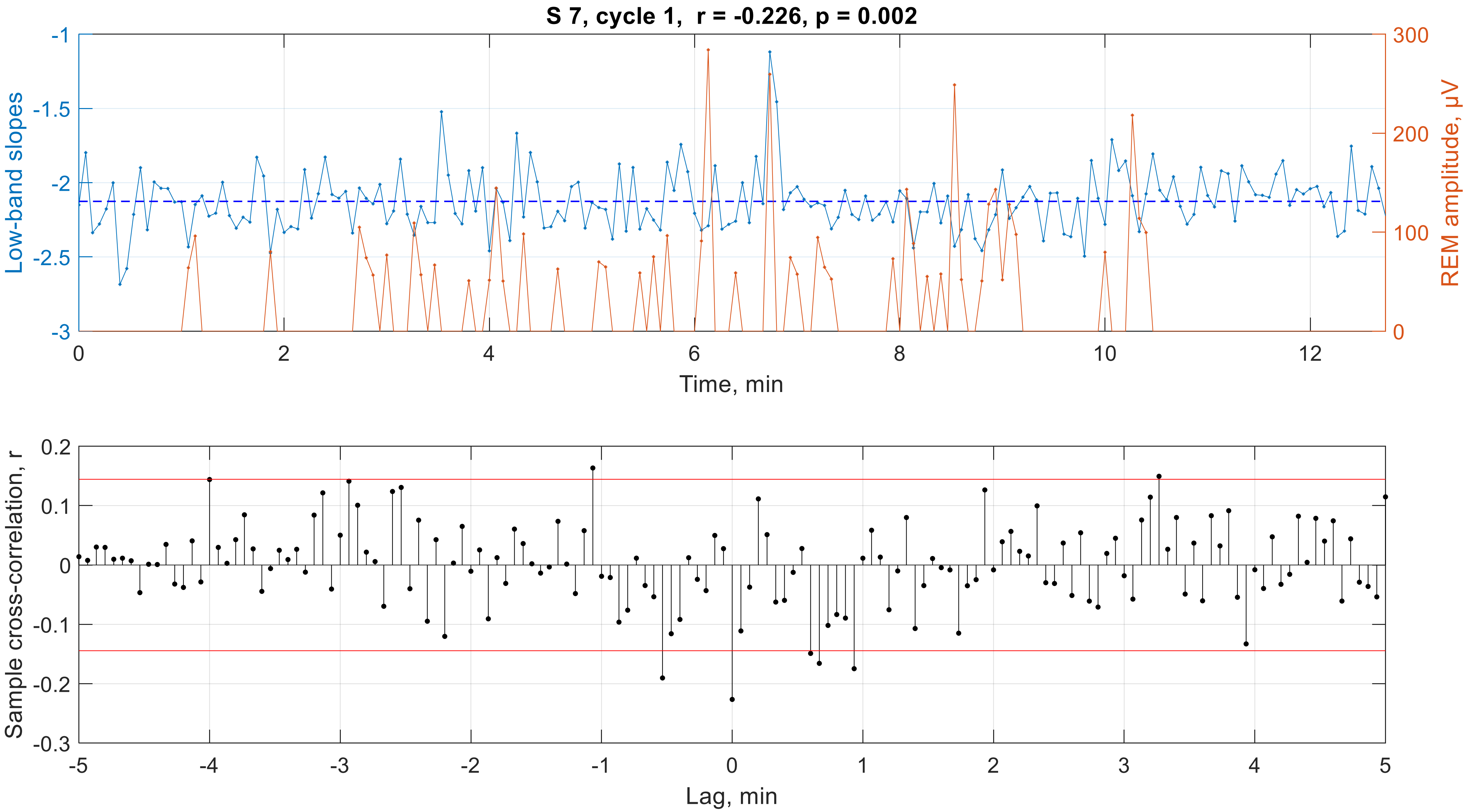

S 7, cycle 1, r = -0.226, p = 0.002
Low-band slopes
REM amplitude, µV
Time, min
-1
-1.5
-2
-2.5
-3
300
200
100
0
0
2
4
6
8
10
12
Sample cross-correlation, r
Lag, min
0.2
0.1
0
-0.1
-0.2
-0.3
-5
-4
-3
-2
-1
0
1
2
3
4
5

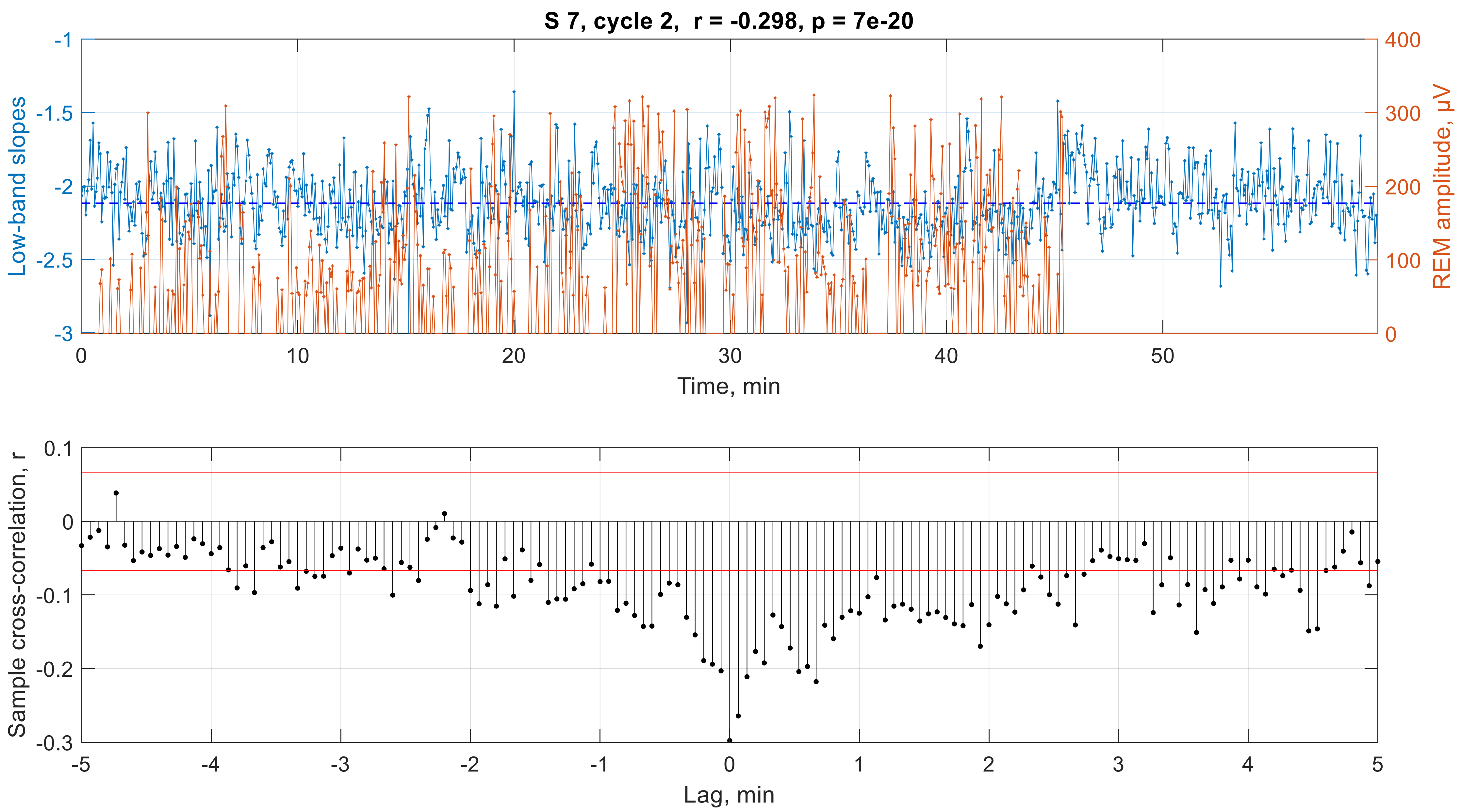
S 7, cycle 2, r = -0.298, p = 7e-20
Low-band slopes
REM amplitude, μV
Time, min
-1
-1.5
-2
-2.5
-3
400
300
200
100
0
0
10
20
30
40
50
Sample cross-correlation, r
Lag, min
0.1
0
-0.1
-0.2
-0.3
-5
-4
-3
-2
-1
0
1
2
3
4
5

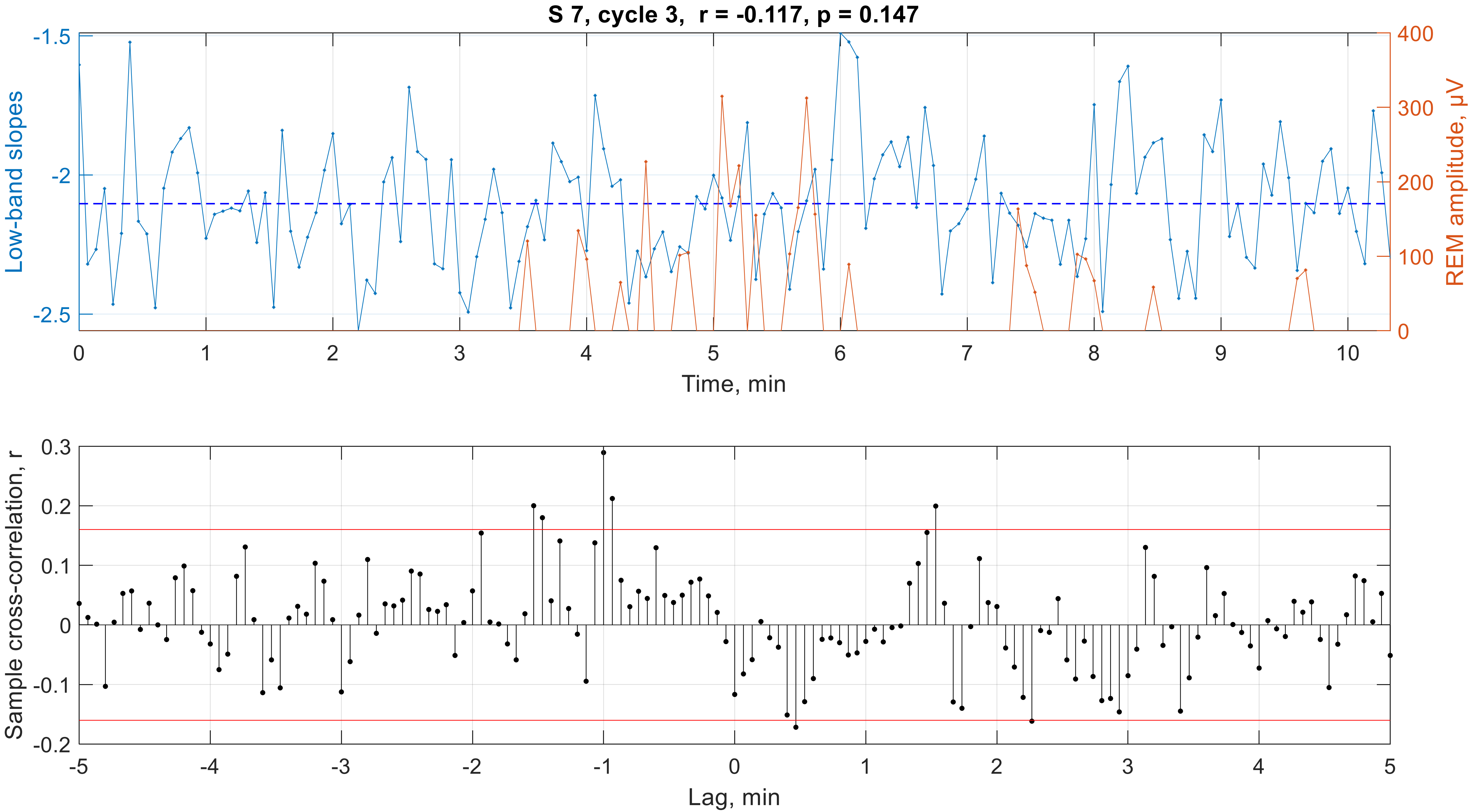

S 7, cycle 3, r = -0.117, p = 0.147
Low-band slopes
REM amplitude, μV
Time, min
Sample cross-correlation, r
Lag, min

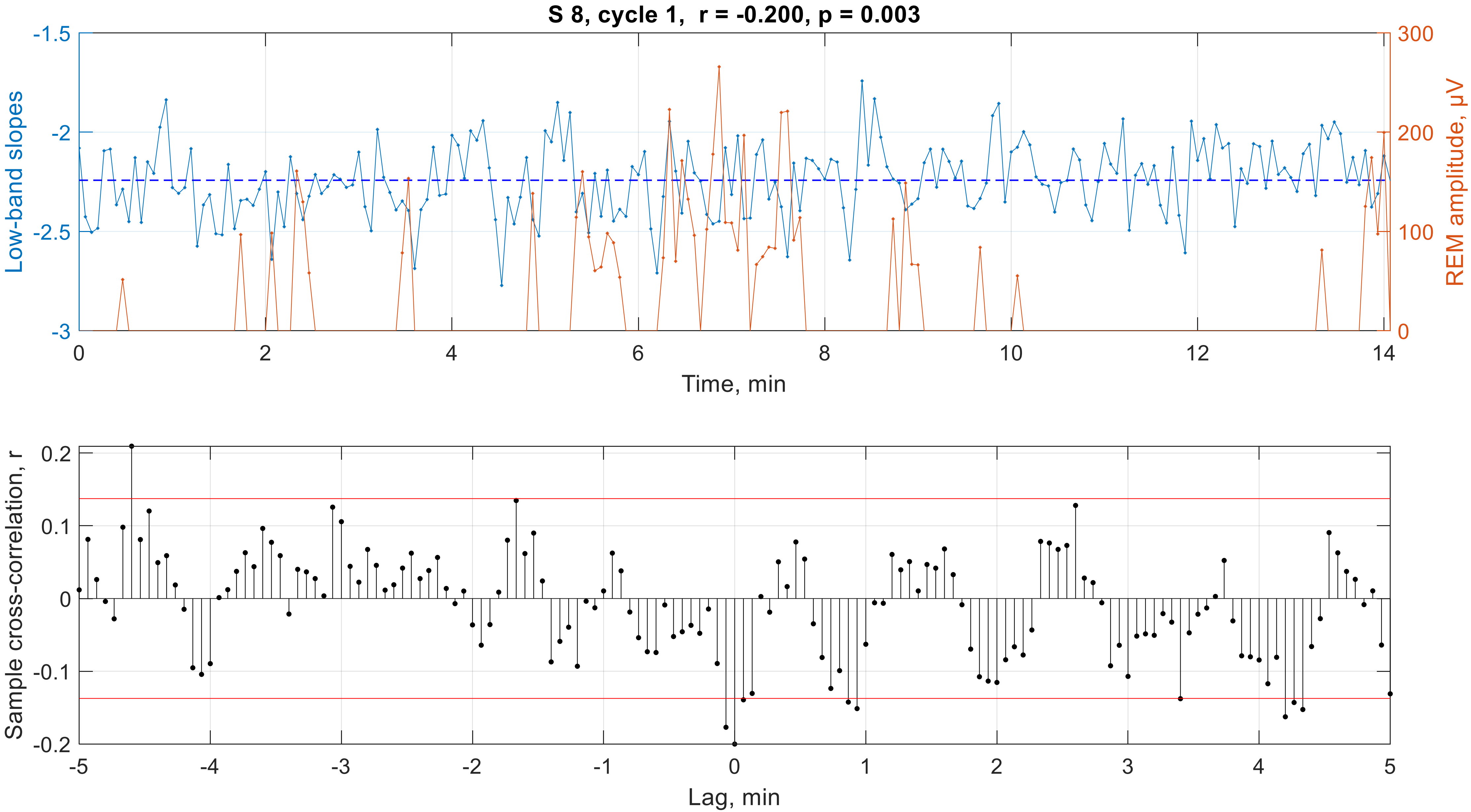

S 8, cycle 1,  r = -0.200, p = 0.003
Low-band slopes
REM amplitude, µV
Time, min
-1.5
-2
-2.5
-3
300
200
100
0
0
2
4
6
8
10
12
14
Sample cross-correlation, r
Lag, min
0.2
0.1
0
-0.1
-0.2
-5
-4
-3
-2
-1
0
1
2
3
4
5

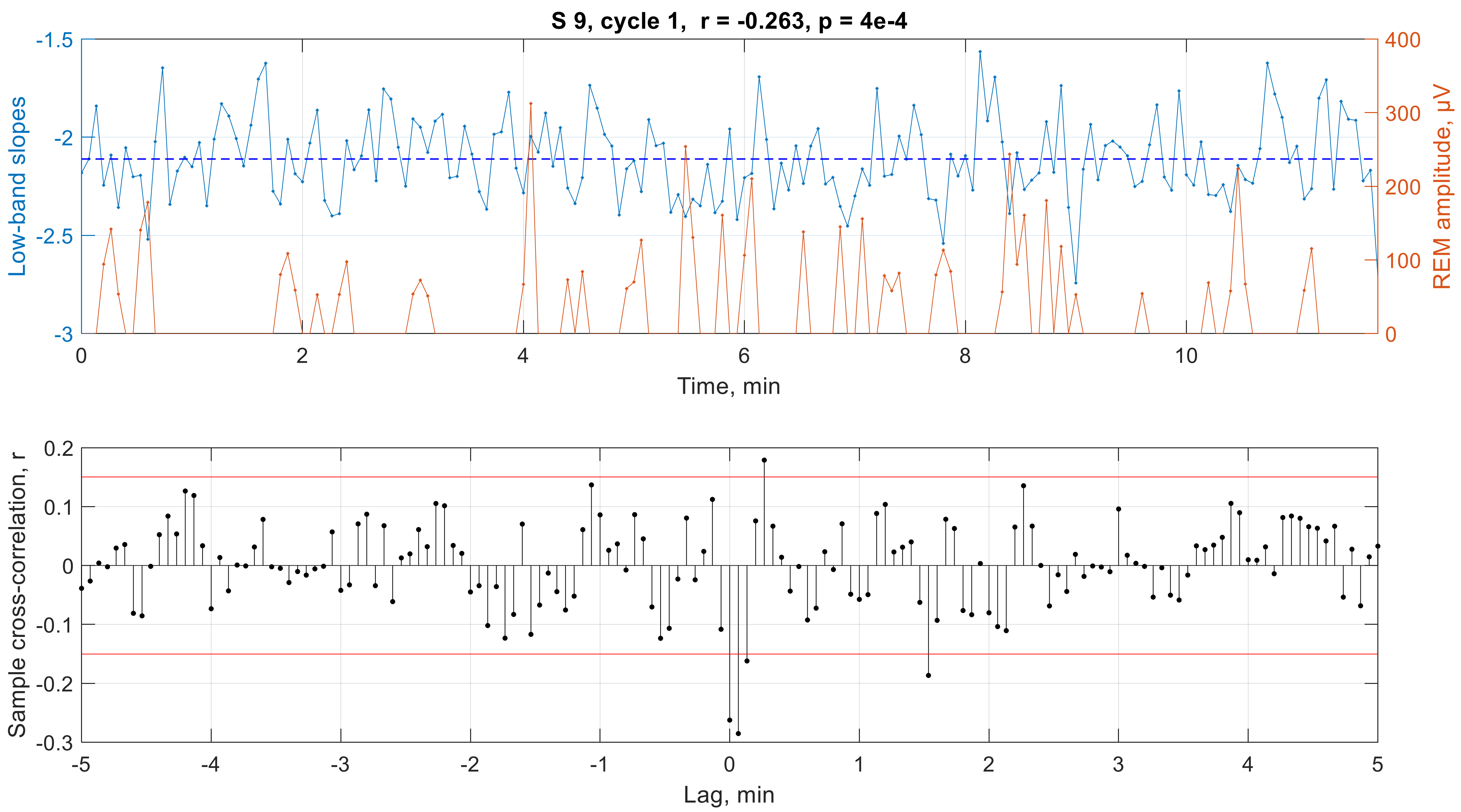
S 9, cycle 1, r = -0.263, p = 4e-4
Low-band slopes
REM amplitude, μV
Time, min
Sample cross-correlation, r
Lag, min
-1.5
-2
-2.5
-3
400
300
200
100
0
0
2
4
6
8
10
0.2
0.1
0
-0.1
-0.2
-0.3
-5
-4
-3
-2
-1
0
1
2
3
4
5

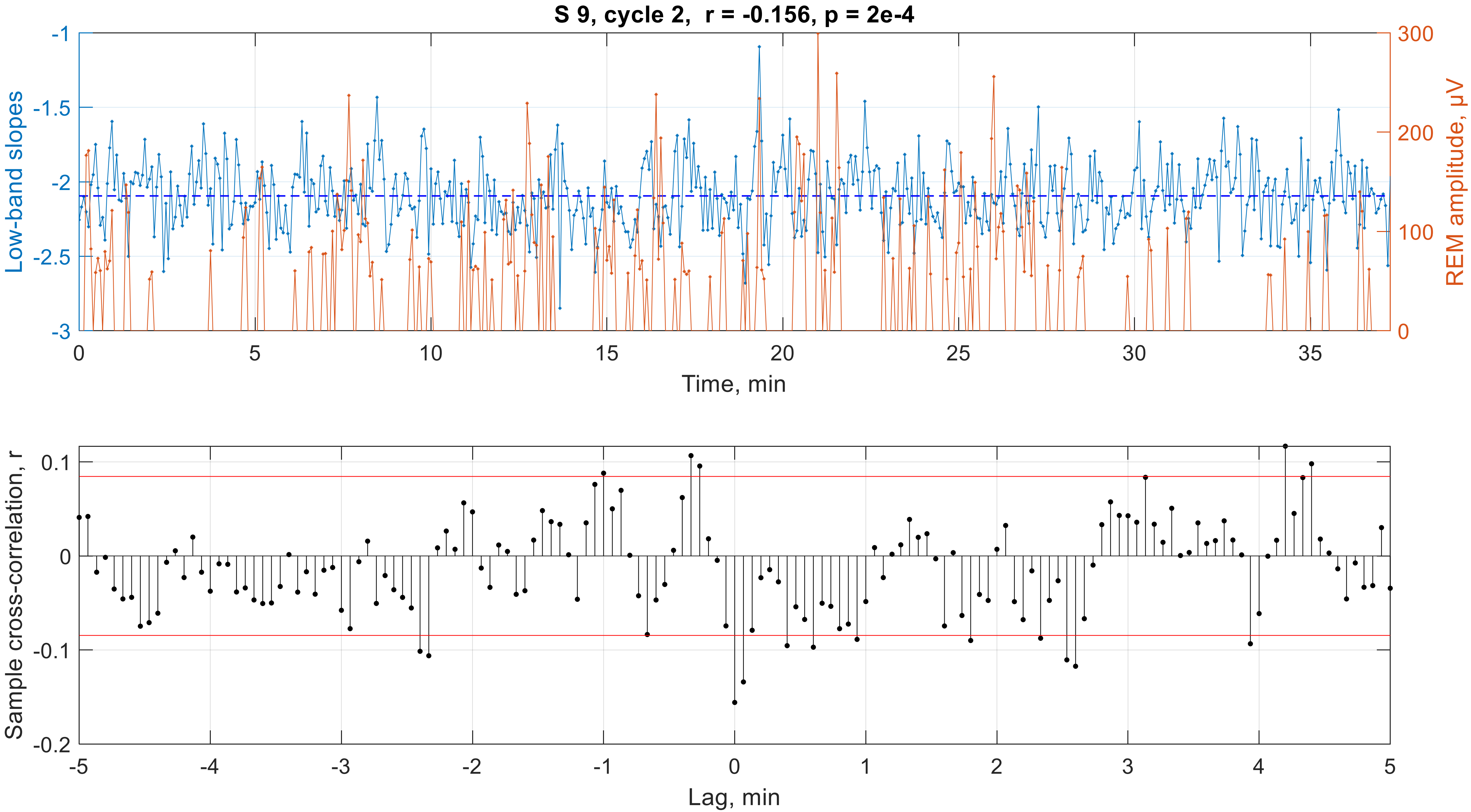

S 9, cycle 2, r = -0.156, p = 2e-4
Low-band slopes
REM amplitude, μV
Time, min
-1
-1.5
-2
-2.5
-3
300
200
100
0
0
5
10
15
20
25
30
35
Sample cross-correlation, r
Lag, min
0.1
0
-0.1
-0.2
-5
-4
-3
-2
-1
0
1
2
3
4
5

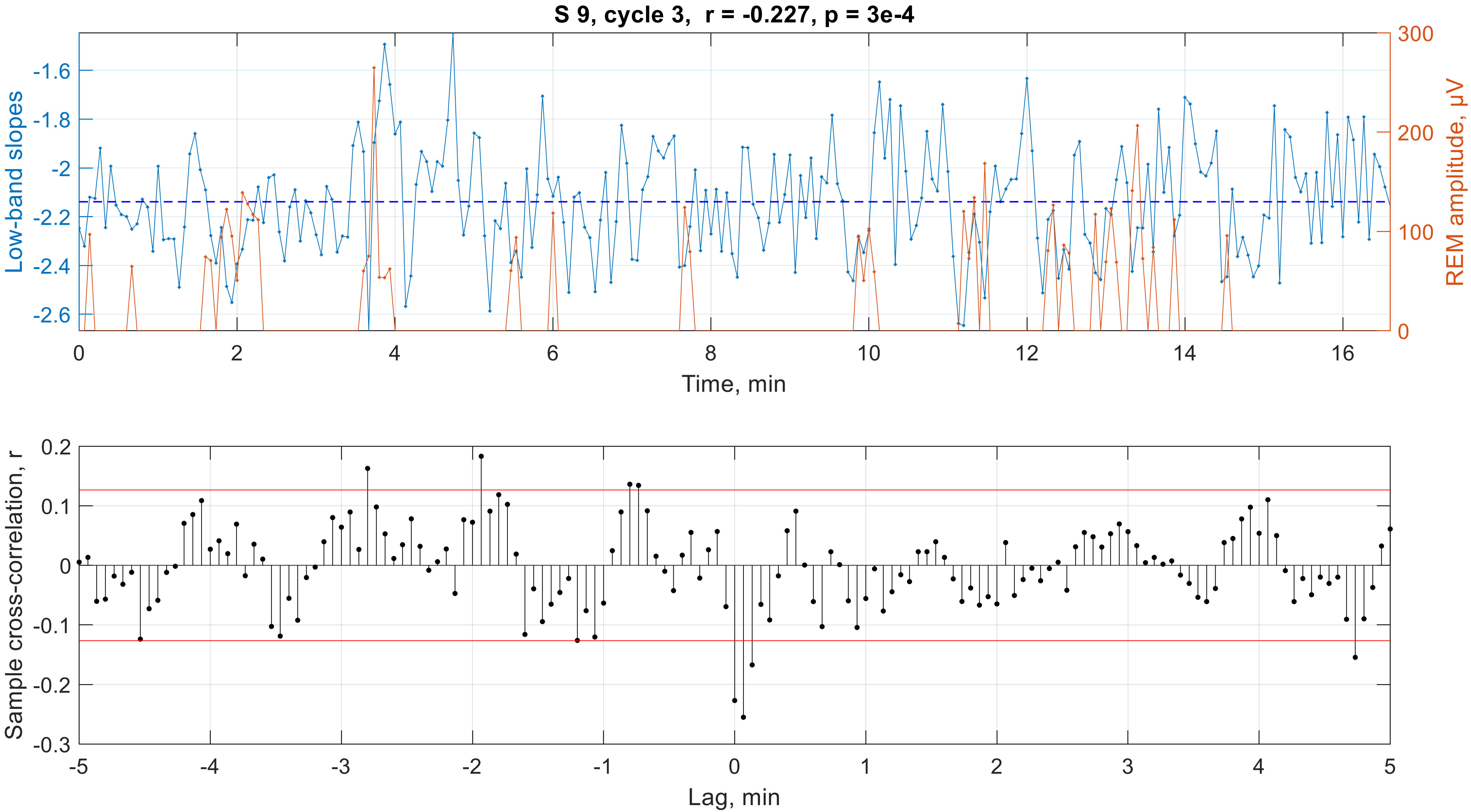

S 9, cycle 3, r = -0.227, p = 3e-4
Low-band slopes
REM amplitude, μV
Time, min
Sample cross-correlation, r
Lag, min

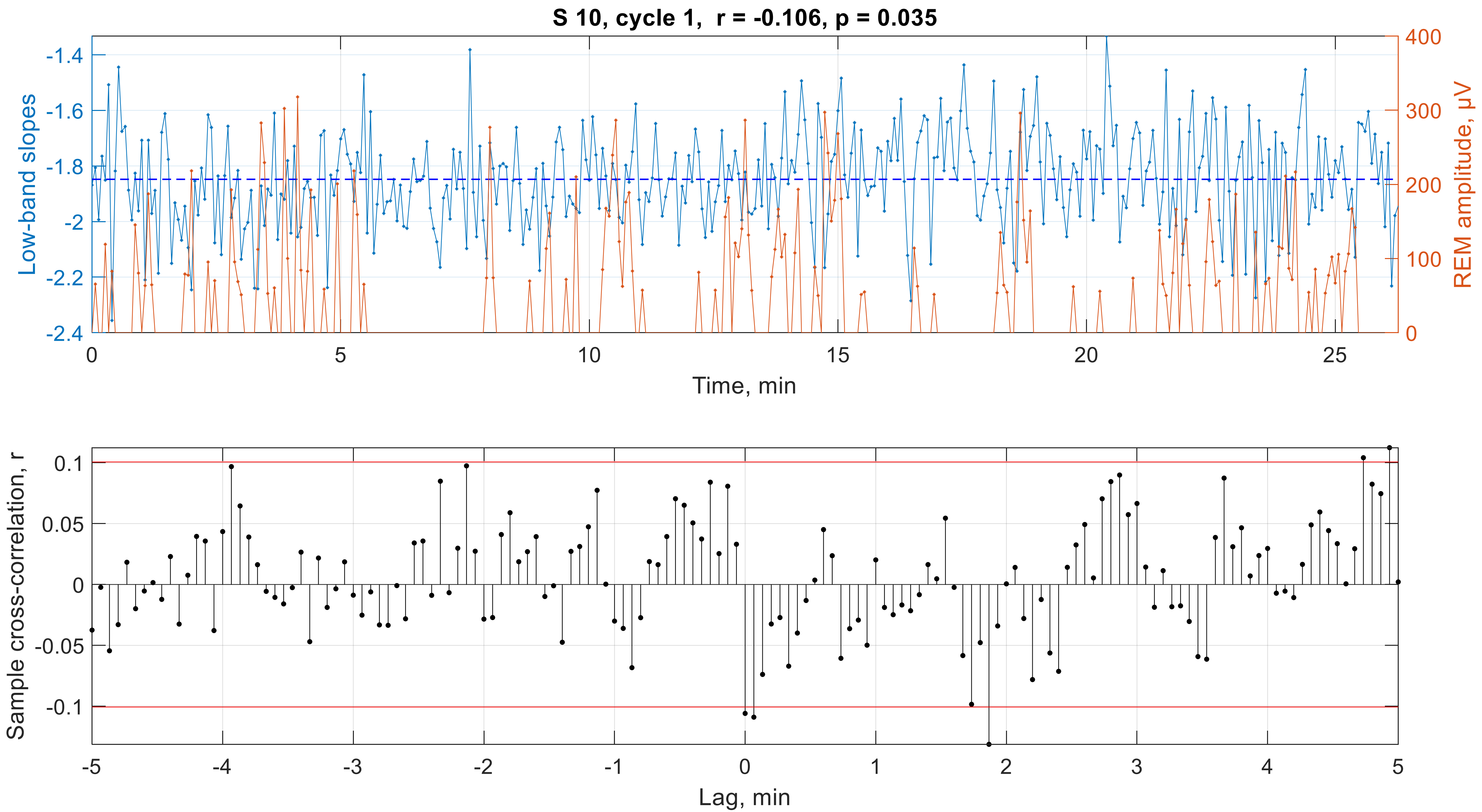

S 10, cycle 1, r = -0.106, p = 0.035
Low-band slopes
-1.4
-1.6
-1.8
-2
-2.2
-2.4
REM amplitude, μV
400
300
200
100
0
0
5
10
15
20
25
Time, min
Sample cross-correlation, r
0.1
0.05
0
-0.05
-0.1
-5
-4
-3
-2
-1
0
1
2
3
4
5
Lag, min

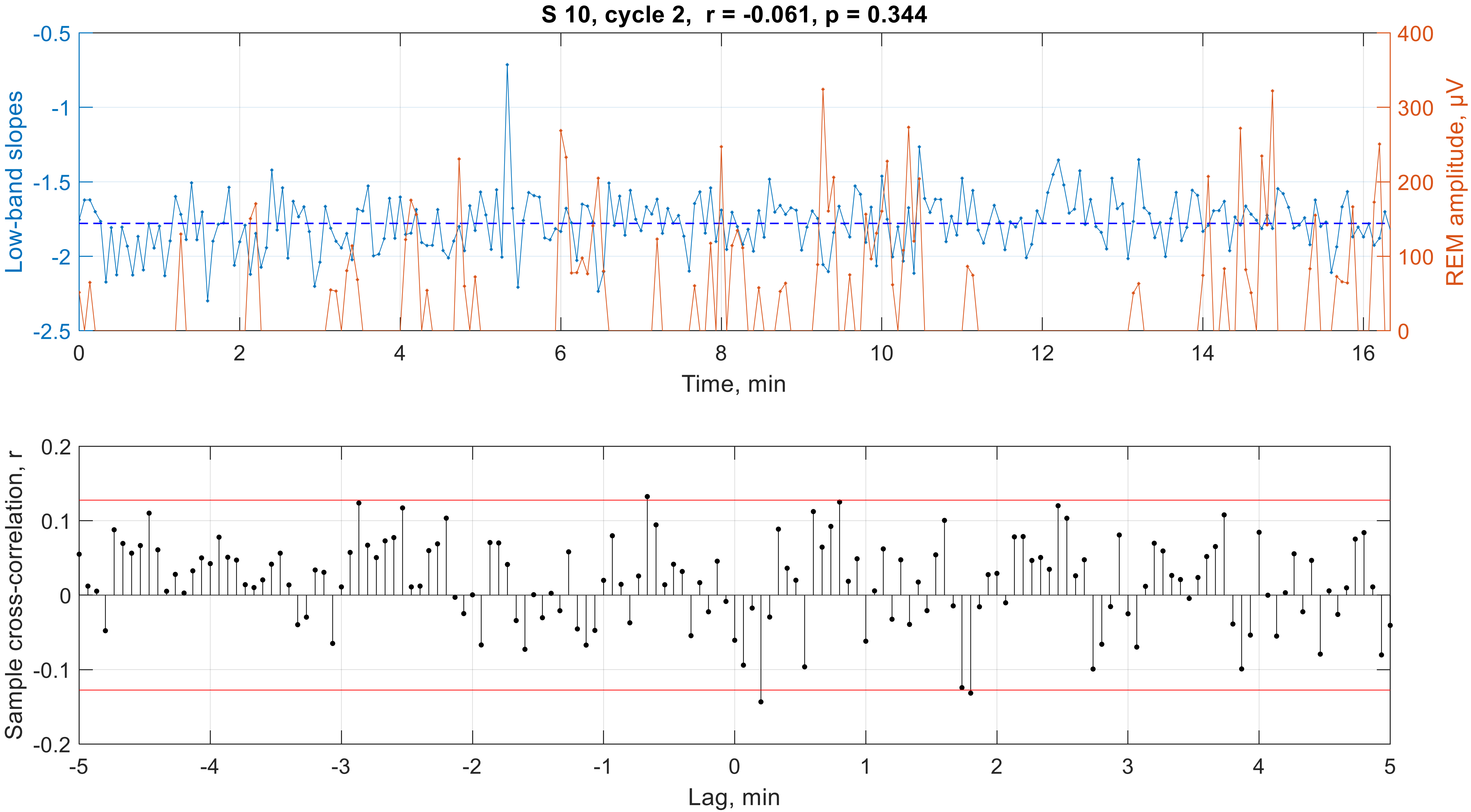
S 10, cycle 2, r = -0.061, p = 0.344
Low-band slopes
REM amplitude, μV
Time, min
Sample cross-correlation, r
Lag, min

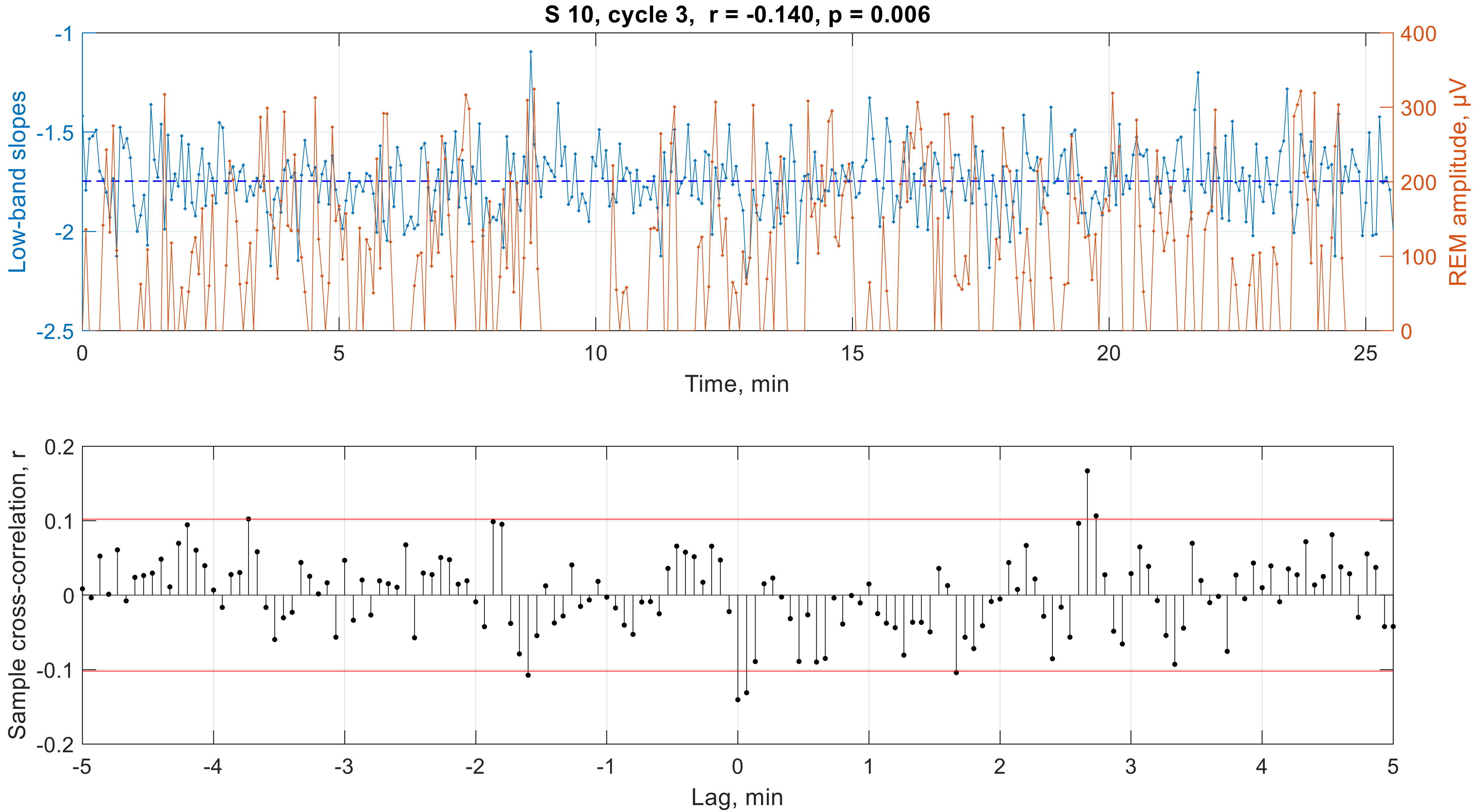

S 10, cycle 3, r = -0.140, p = 0.006
Low-band slopes
REM amplitude, μV
Time, min
Sample cross-correlation, r
Lag, min

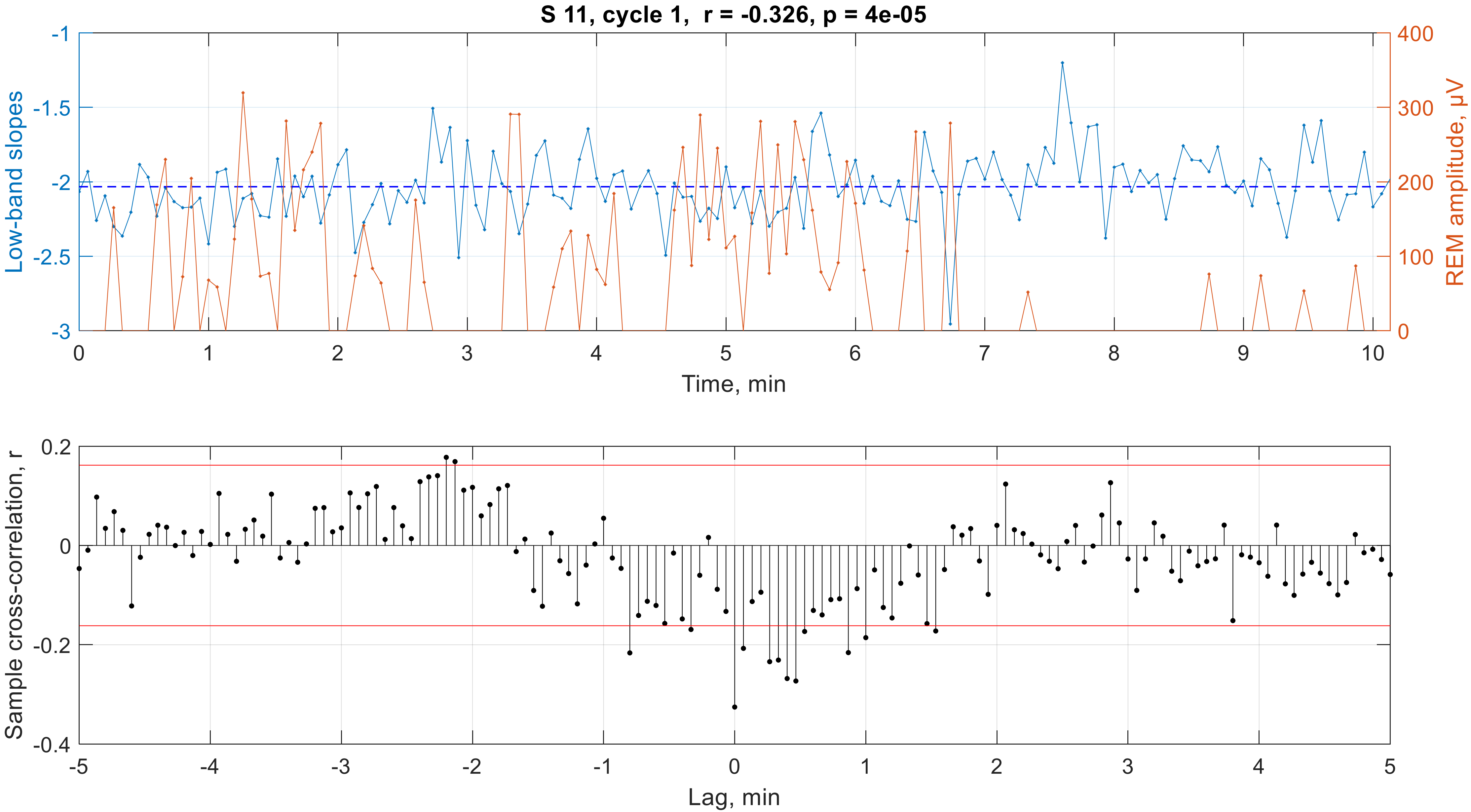

S 11, cycle 1, r = -0.326, p = 4e-05
Low-band slopes
REM amplitude, μV
Time, min
Sample cross-correlation, r
Lag, min

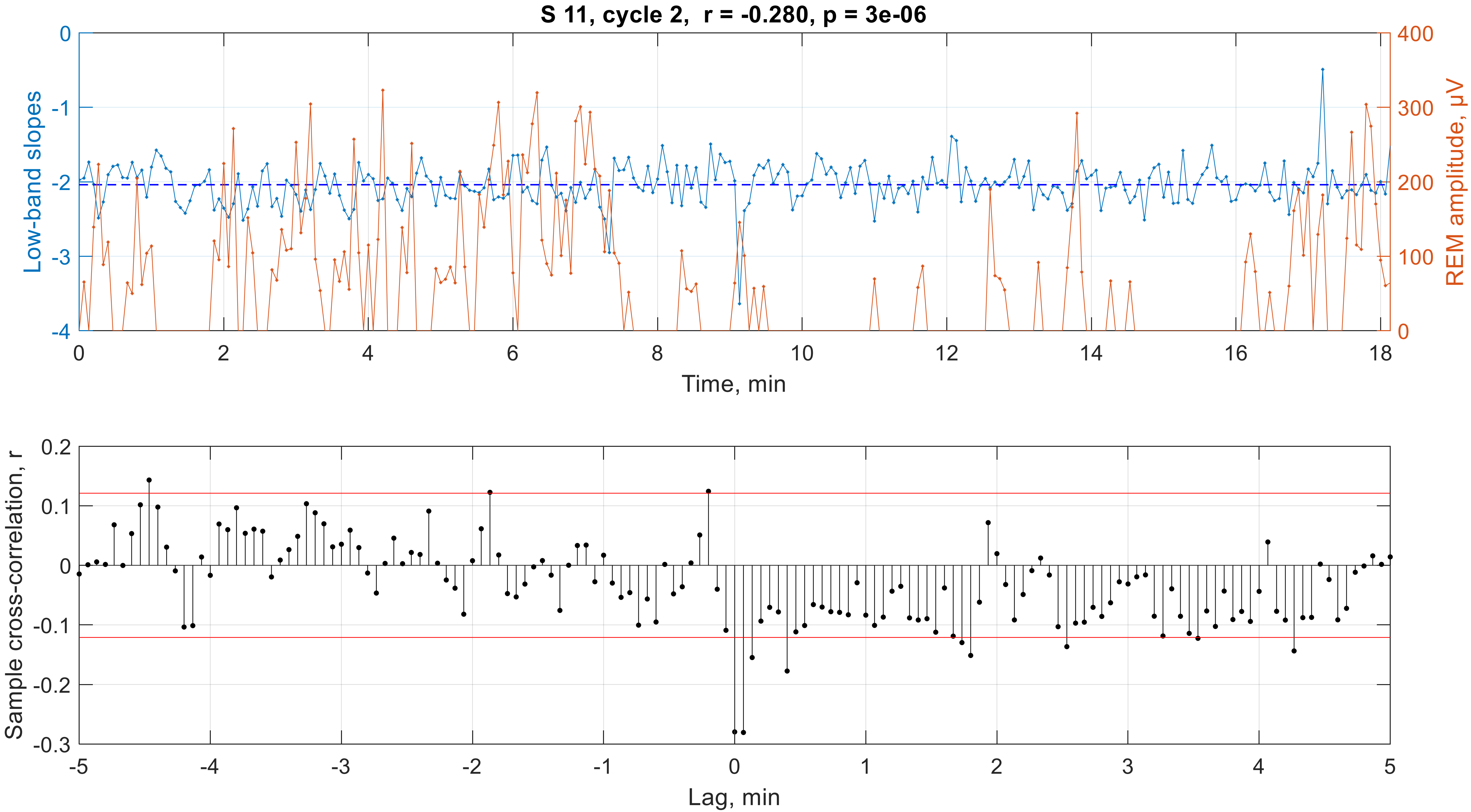

S 11, cycle 2, r = -0.280, p = 3e-06
Low-band slopes
REM amplitude, μV
Time, min
0
-1
-2
-3
-4
400
300
200
100
0
2
4
6
8
10
12
14
16
18
Sample cross-correlation, r
Lag, min
0.2
0.1
-0.1
-0.2
-0.3
-5
-4
-3
-2
-1
1
3
5

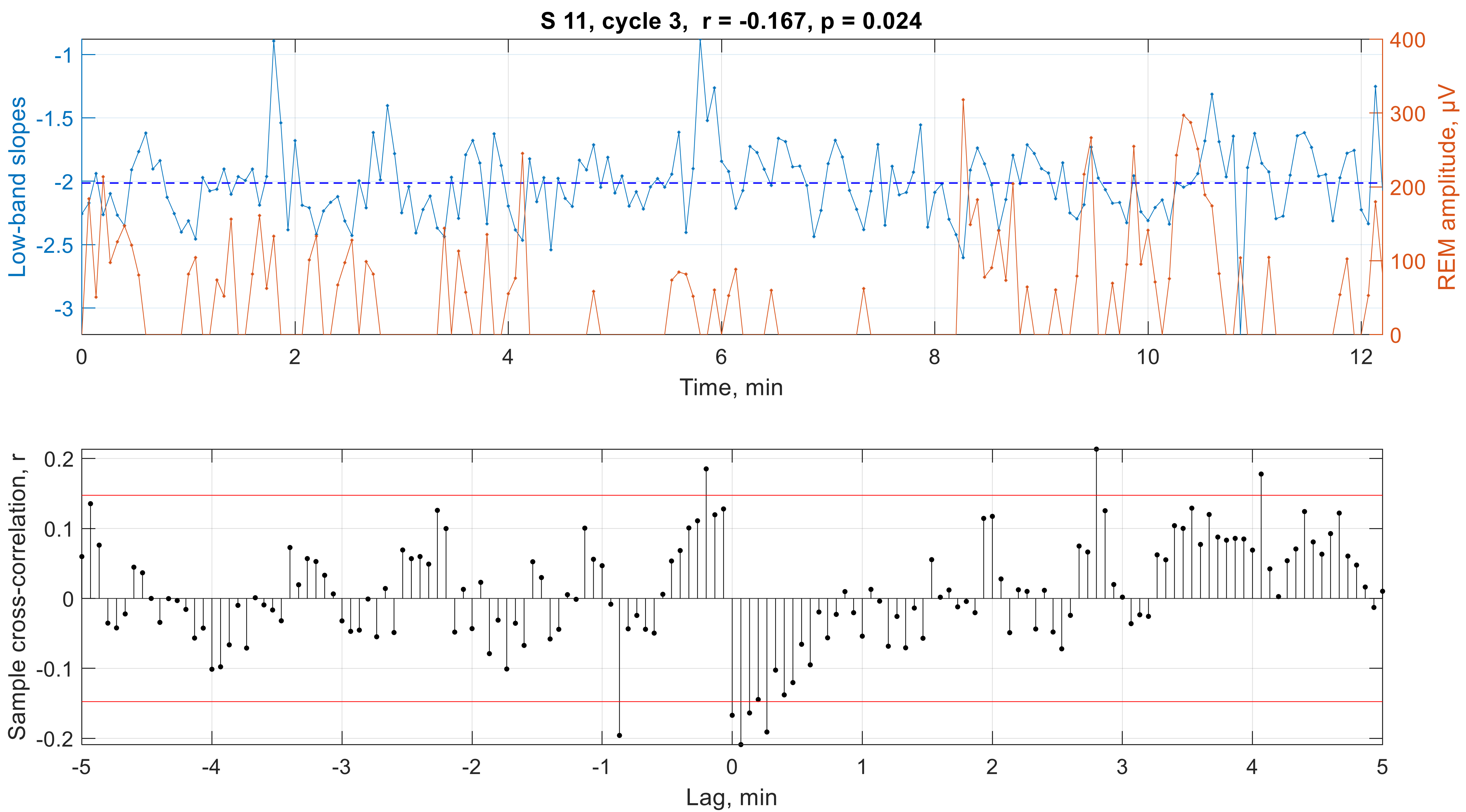

S 11, cycle 3, r = -0.167, p = 0.024
Low-band slopes
REM amplitude, μV
Time, min
Sample cross-correlation, r
Lag, min

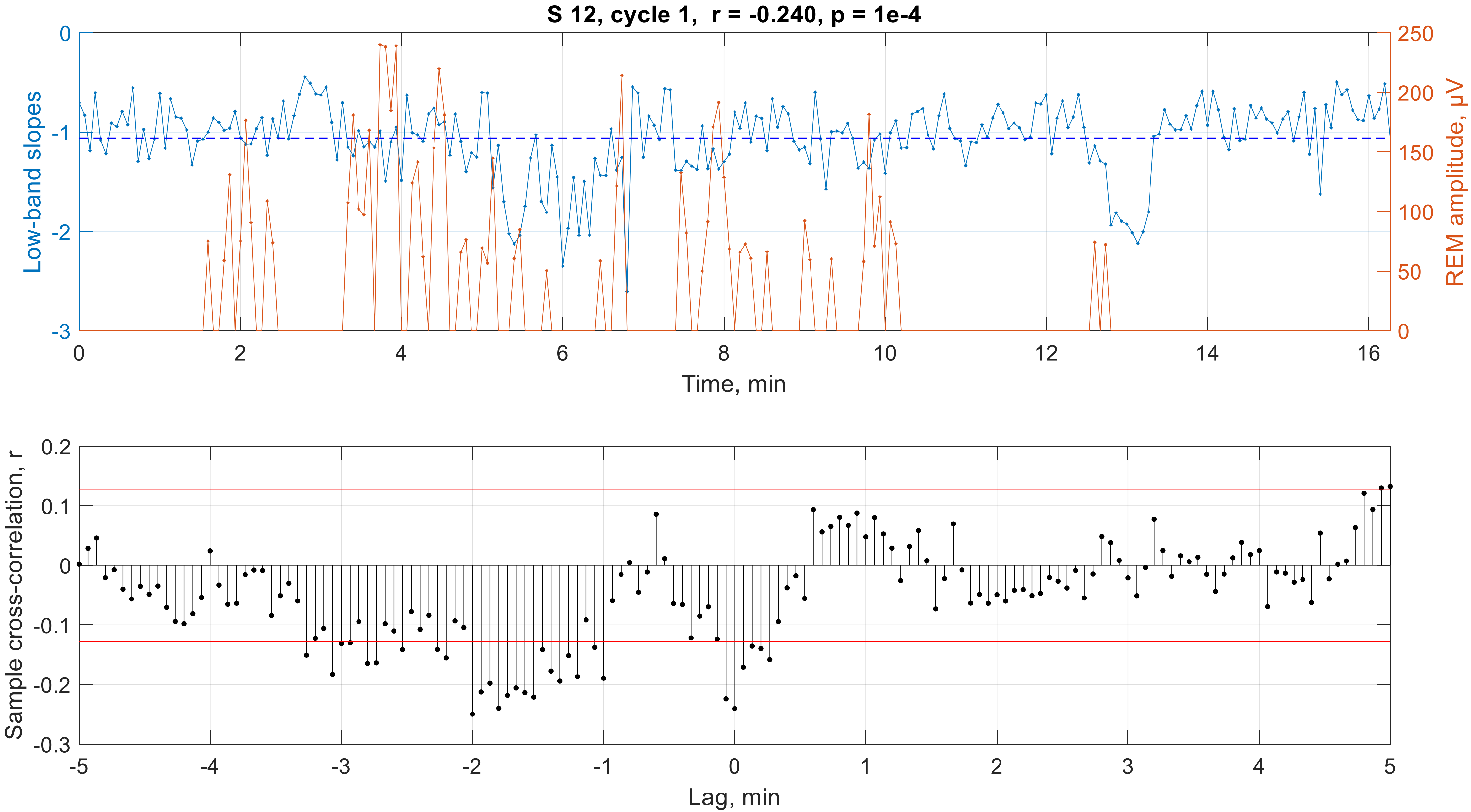

S 12, cycle 1, r = -0.240, p = 1e-4
Low-band slopes
REM amplitude, μV
Time, min
0
-1
-2
-3
250
200
150
100
50
0
2
4
6
8
10
12
14
16
Sample cross-correlation, r
Lag, min
0.2
0.1
-0.1
-0.2
-0.3
-5
-4
-3
-2
-1
1
3
5

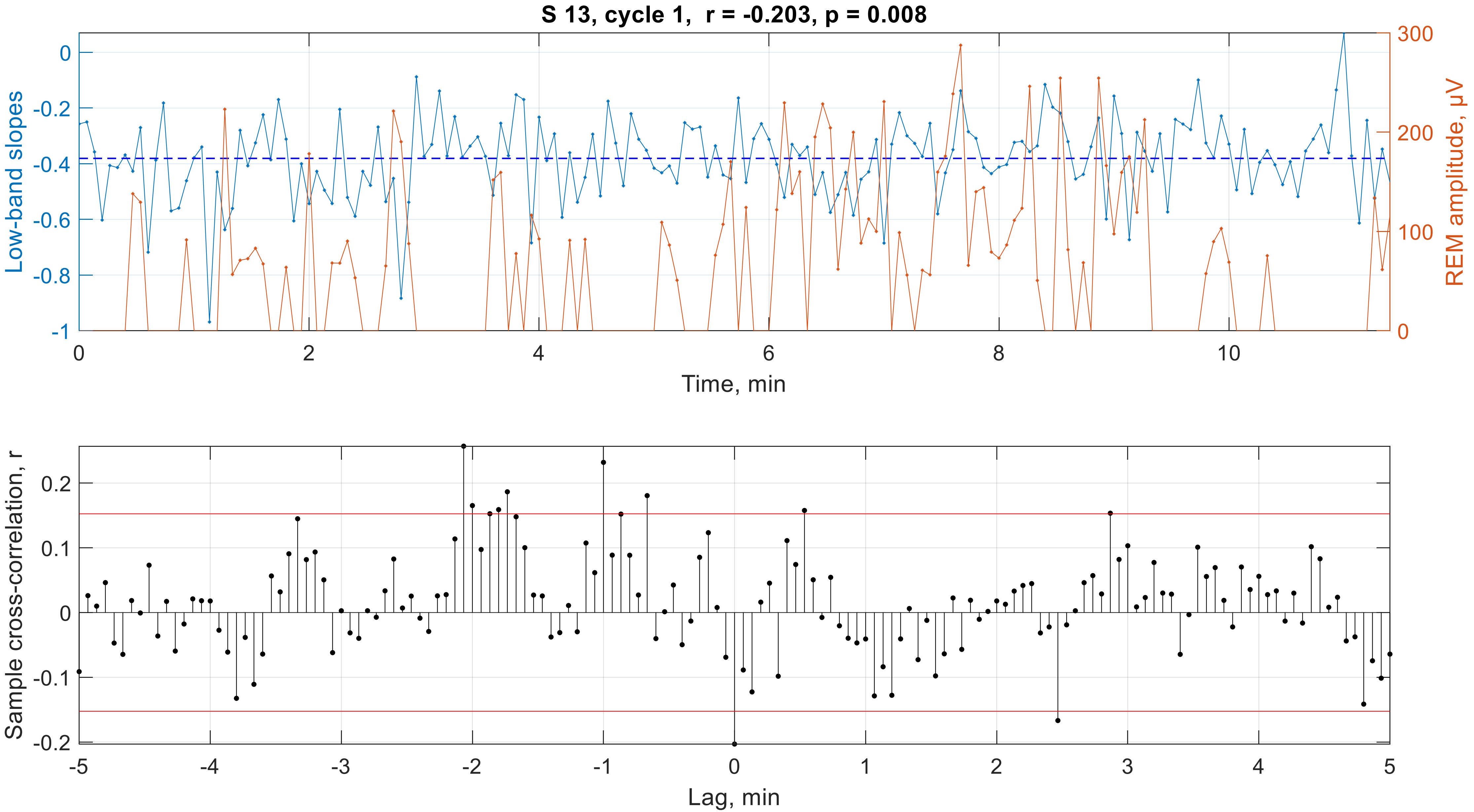
S 13, cycle 1, r = -0.203, p = 0.008
Low-band slopes
REM amplitude, μV
Time, min
0
-0.2
-0.4
-0.6
-0.8
-1
300
200
100
0
2
4
6
8
10
Sample cross-correlation, r
Lag, min
0.2
0.1
0
-0.1
-0.2
-5
-4
-3
-2
-1
0
1
2
3
4
5

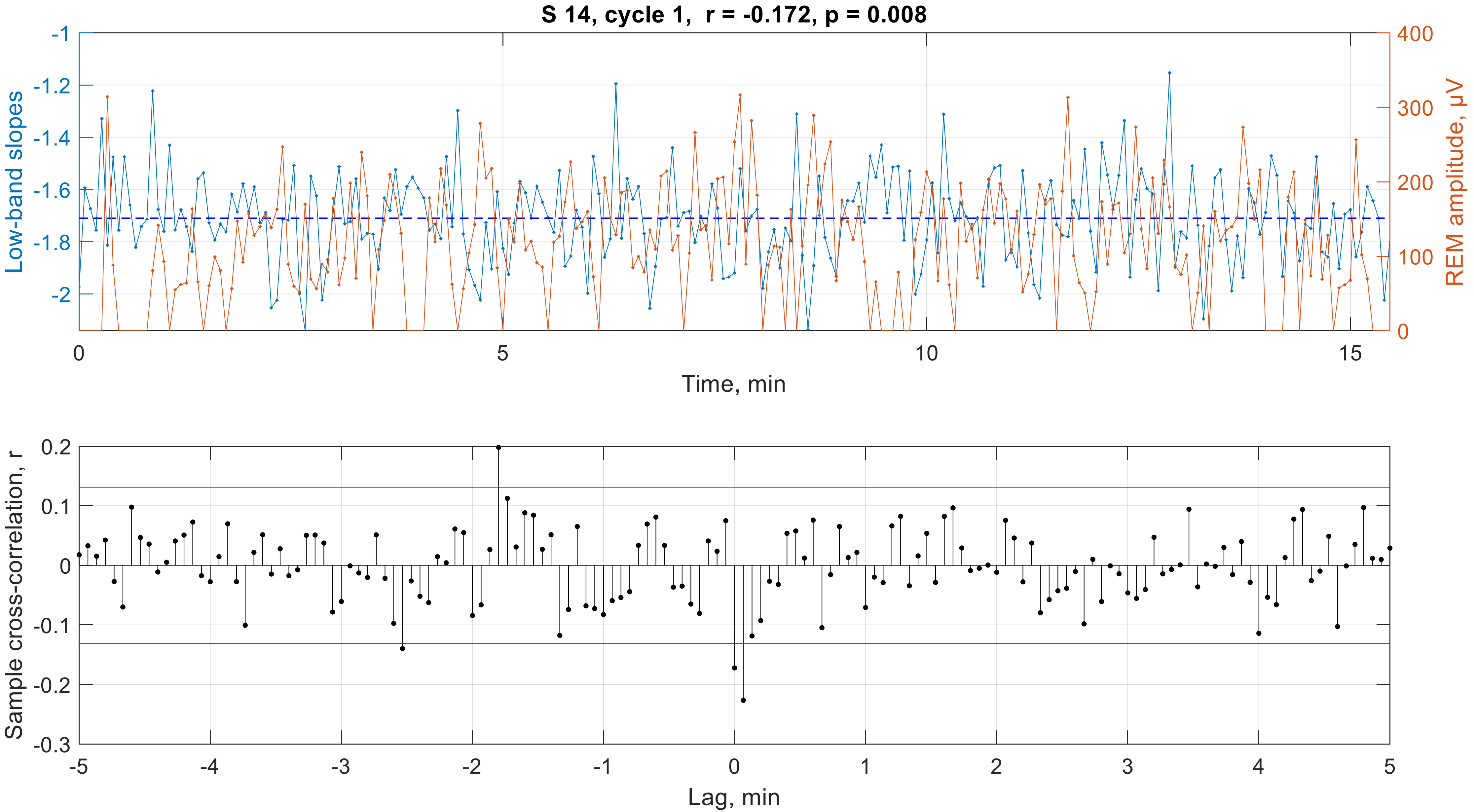

S 14, cycle 1, r = -0.172, p = 0.008
Low-band slopes
REM amplitude, μV
Time, min
Sample cross-correlation, r
Lag, min

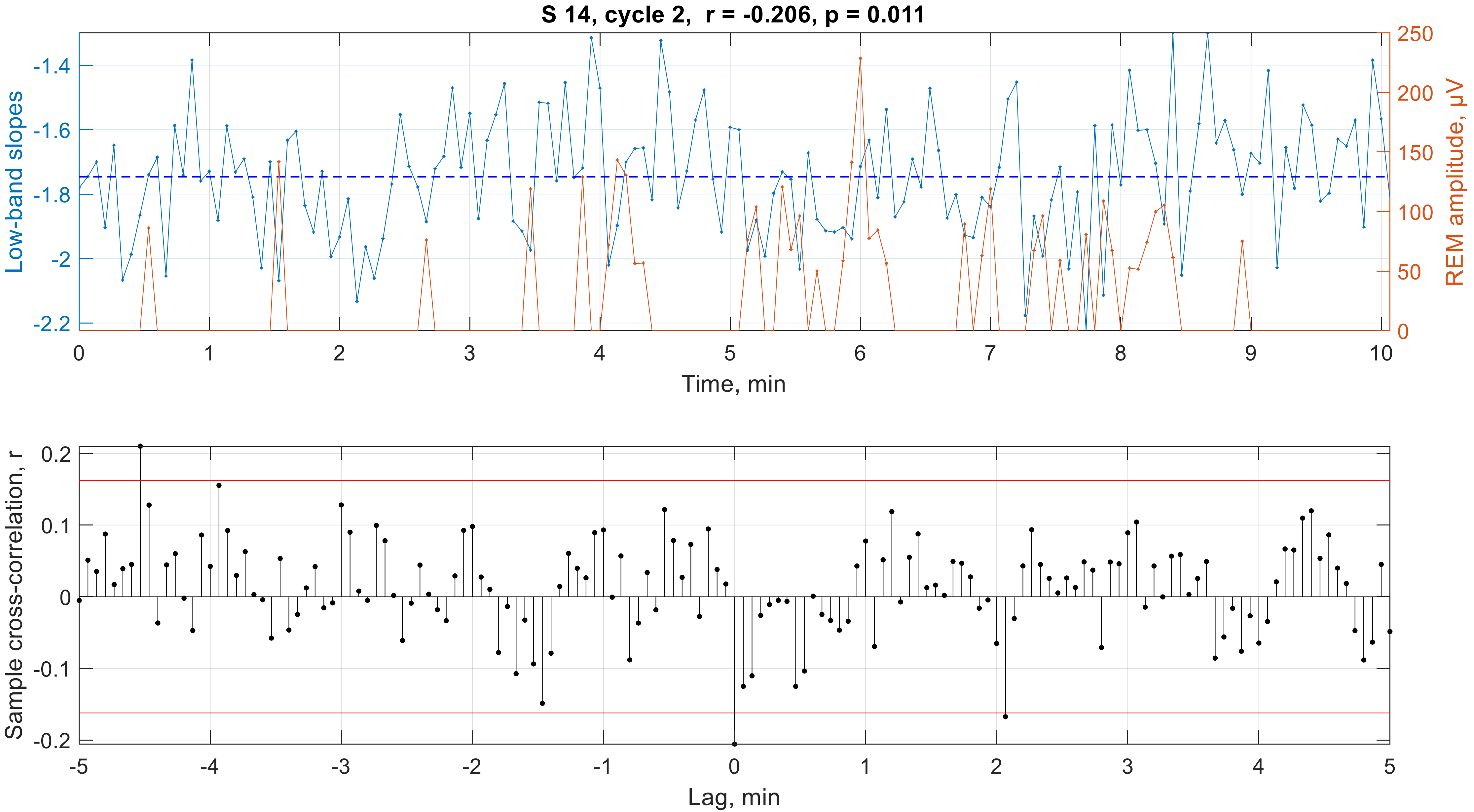
S 14, cycle 2, r = -0.206, p = 0.011
Low-band slopes
REM amplitude, μV
Time, min
Sample cross-correlation, r
Lag, min

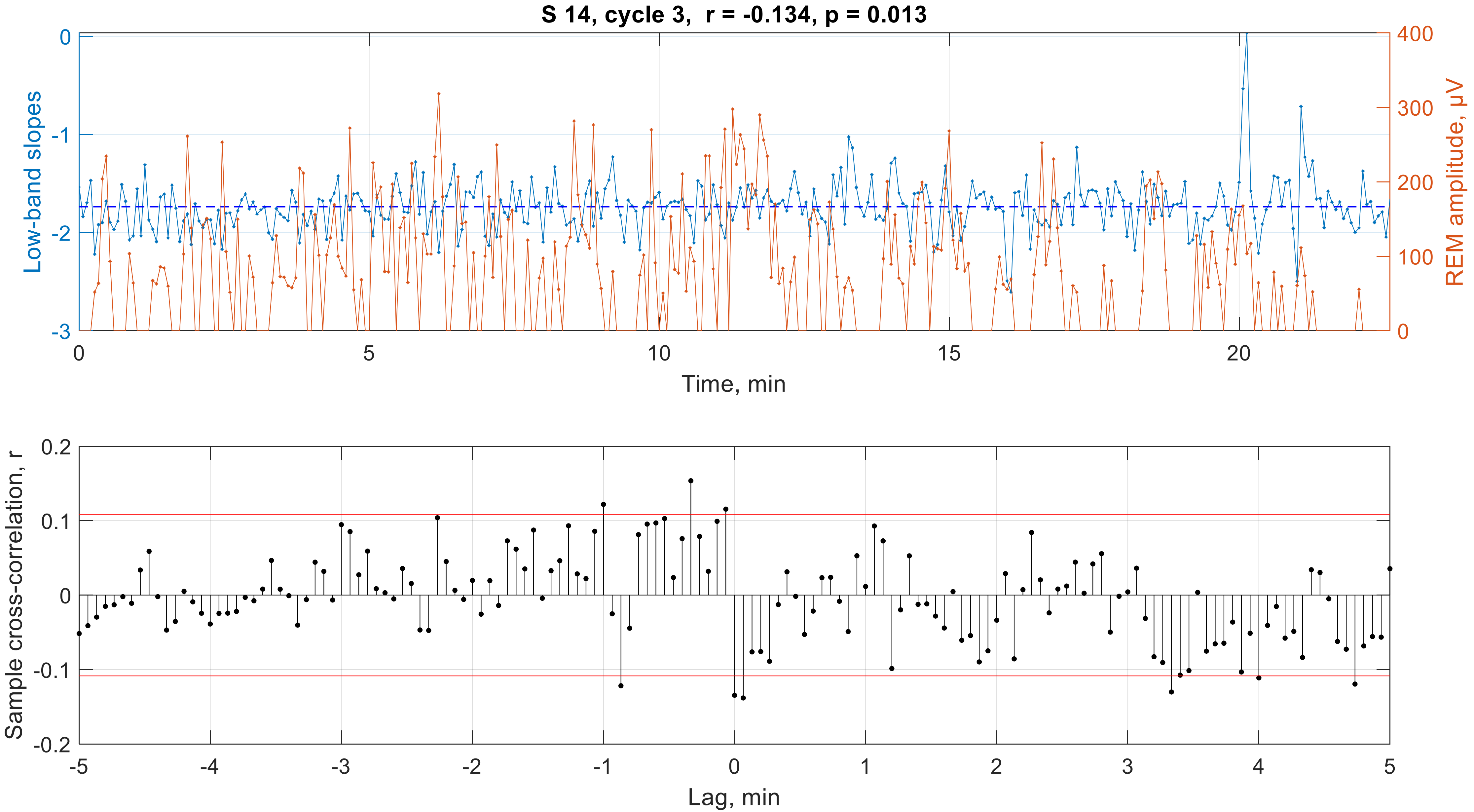
S 14, cycle 3, r = -0.134, p = 0.013
Low-band slopes
REM amplitude, μV
Time, min
Sample cross-correlation, r
Lag, min

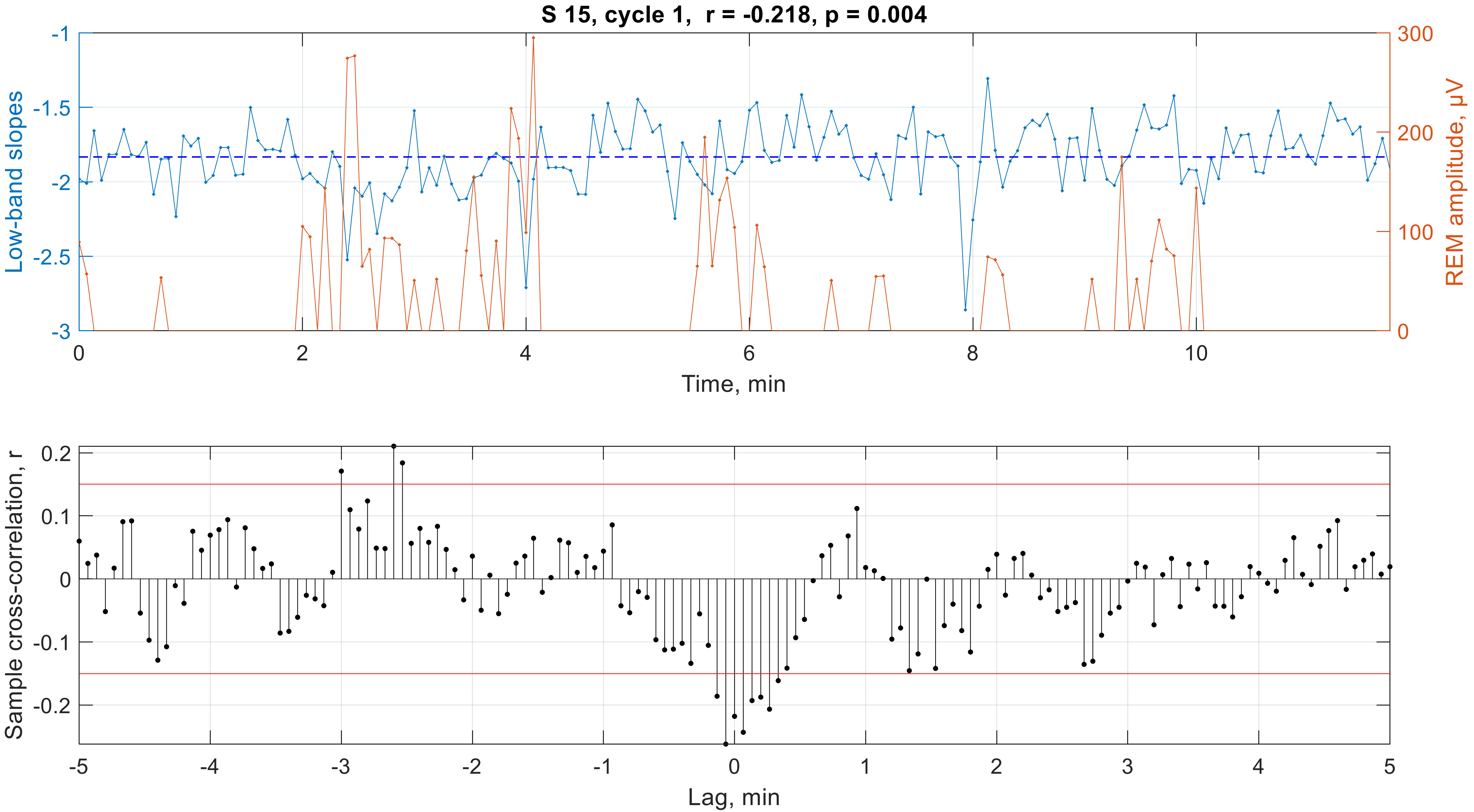

S 15, cycle 1,  r = -0.218, p = 0.004
Low-band slopes
REM amplitude, μV
Time, min
Sample cross-correlation, r
Lag, min

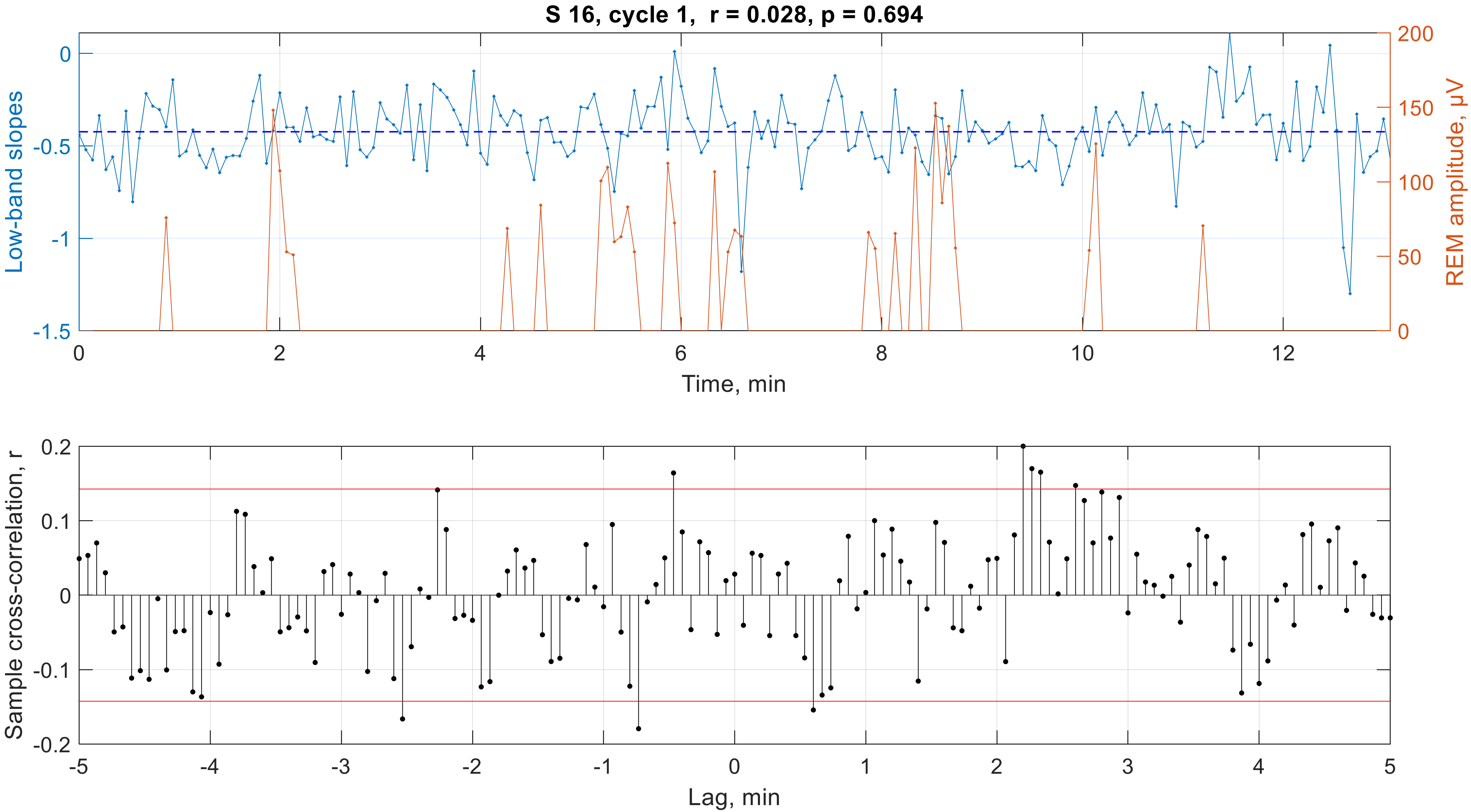

S 16, cycle 1,  r = 0.028, p = 0.694
Low-band slopes
REM amplitude, μV
Time, min
Sample cross-correlation, r
Lag, min

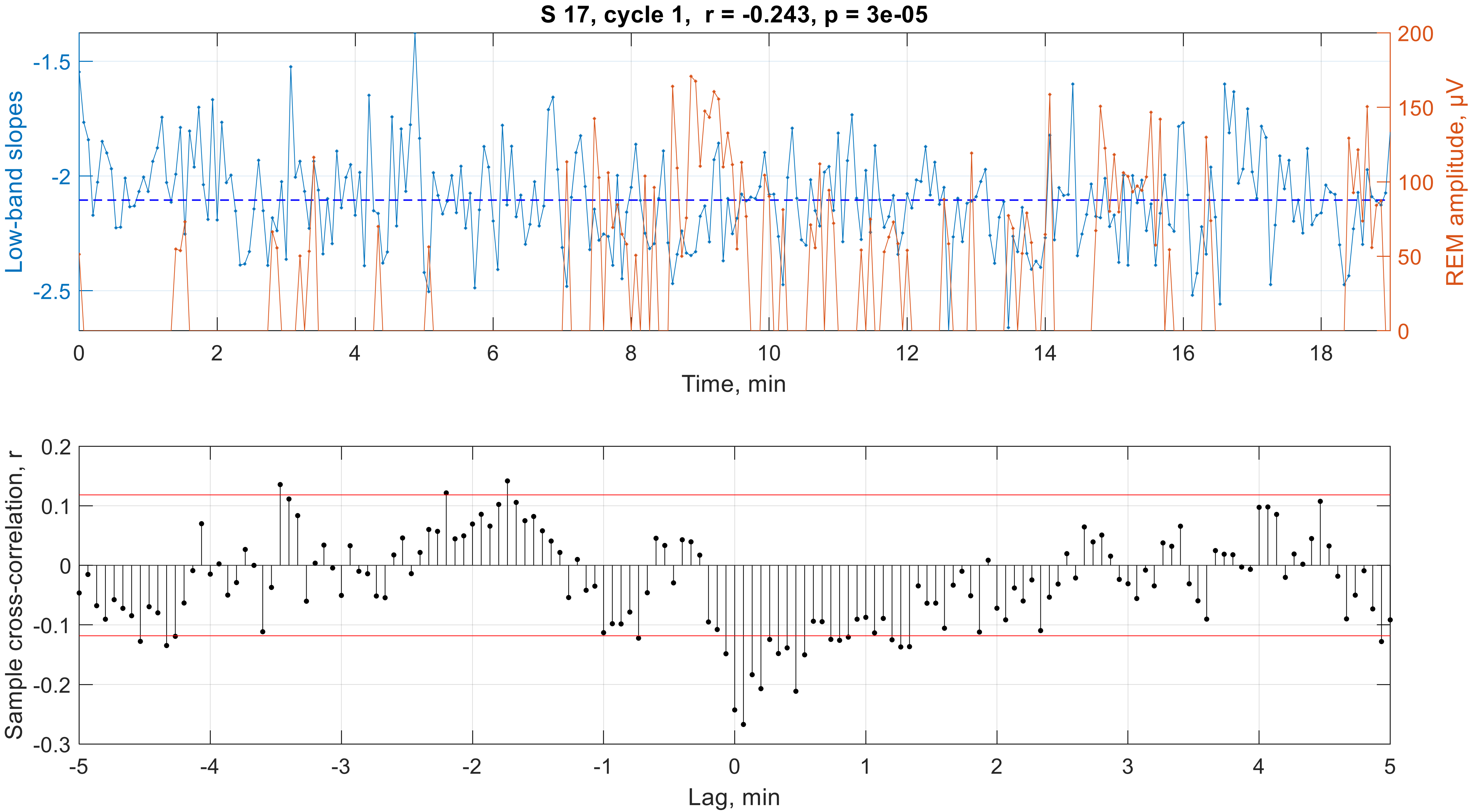

S 17, cycle 1, r = -0.243, p = 3e-05
Low-band slopes
-1.5
-2
-2.5
REM amplitude, µV
200
150
100
50
0
0
2
4
6
8
10
12
14
16
18
Time, min
Sample cross-correlation, r
0.2
0.1
0
-0.1
-0.2
-0.3
-5
-4
-3
-2
-1
0
1
2
3
4
5
Lag, min

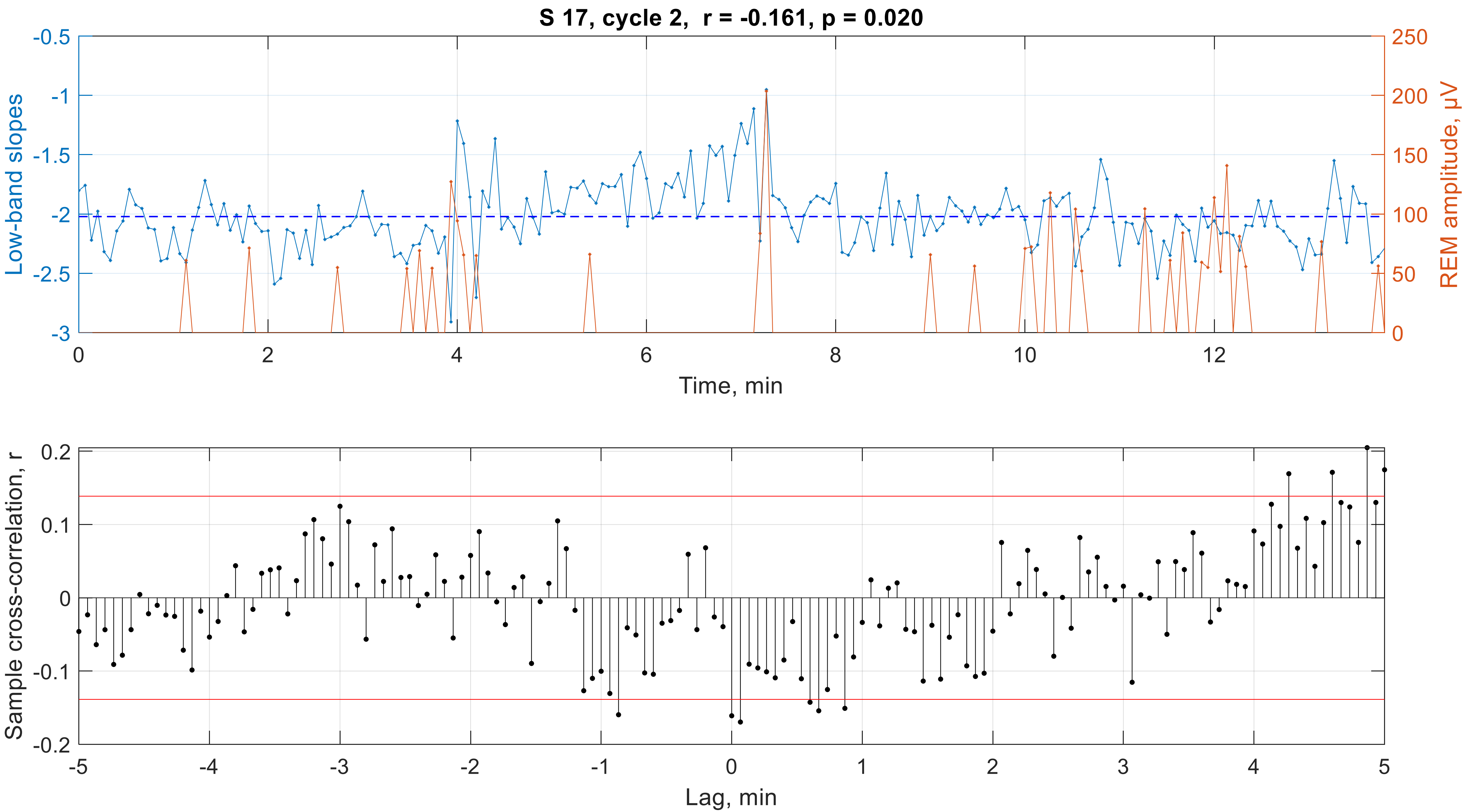
S 17, cycle 2, r = -0.161, p = 0.020
Low-band slopes
REM amplitude, μV
Time, min
Sample cross-correlation, r
Lag, min

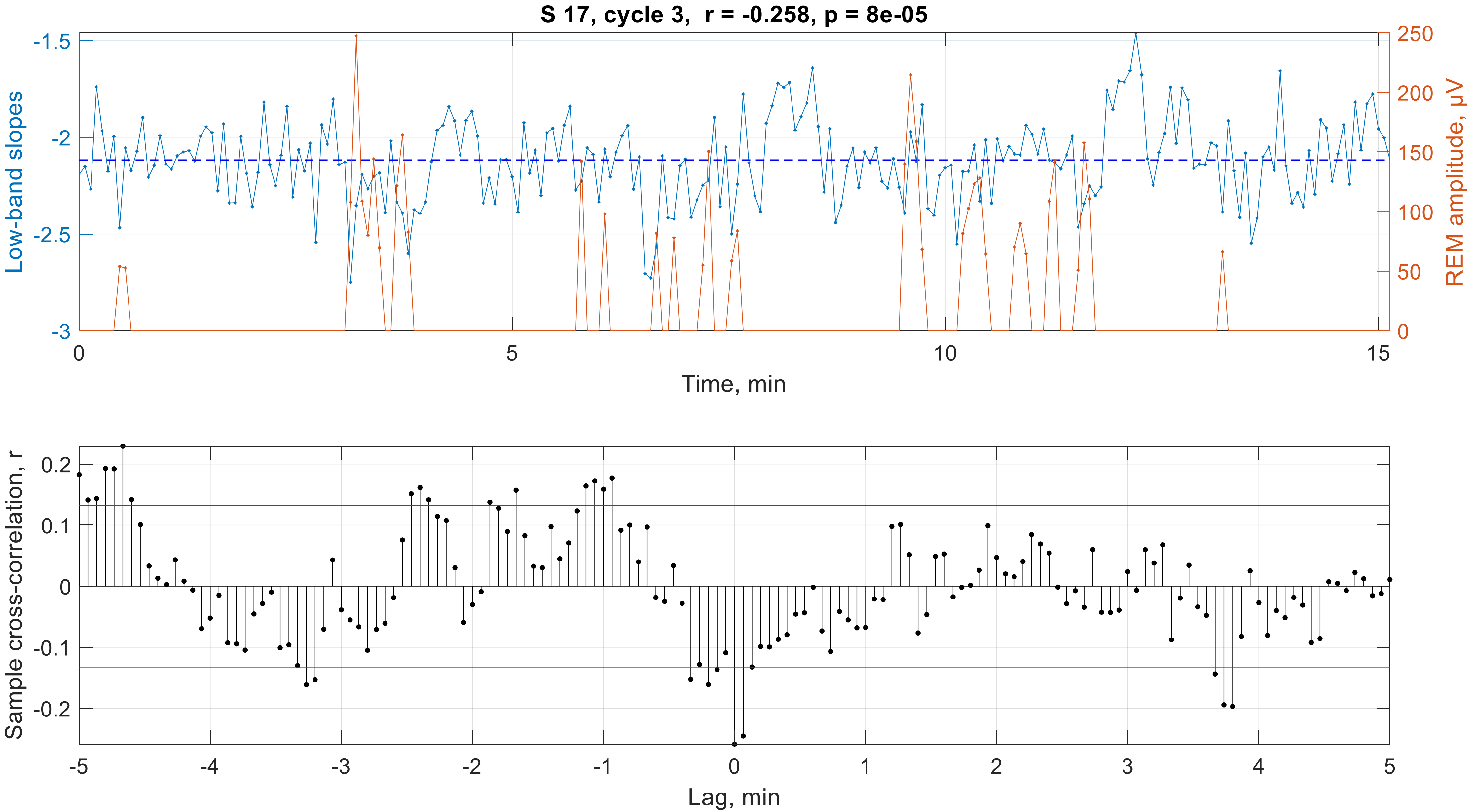
S 17, cycle 3, r = -0.258, p = 8e-05
Low-band slopes
REM amplitude, μV
Time, min
-1.5
-2
-2.5
-3
250
200
150
100
50
0
0
5
10
15
Sample cross-correlation, r
Lag, min
0.2
0.1
0
-0.1
-0.2
-5
-4
-3
-2
-1
0
1
2
3
4
5

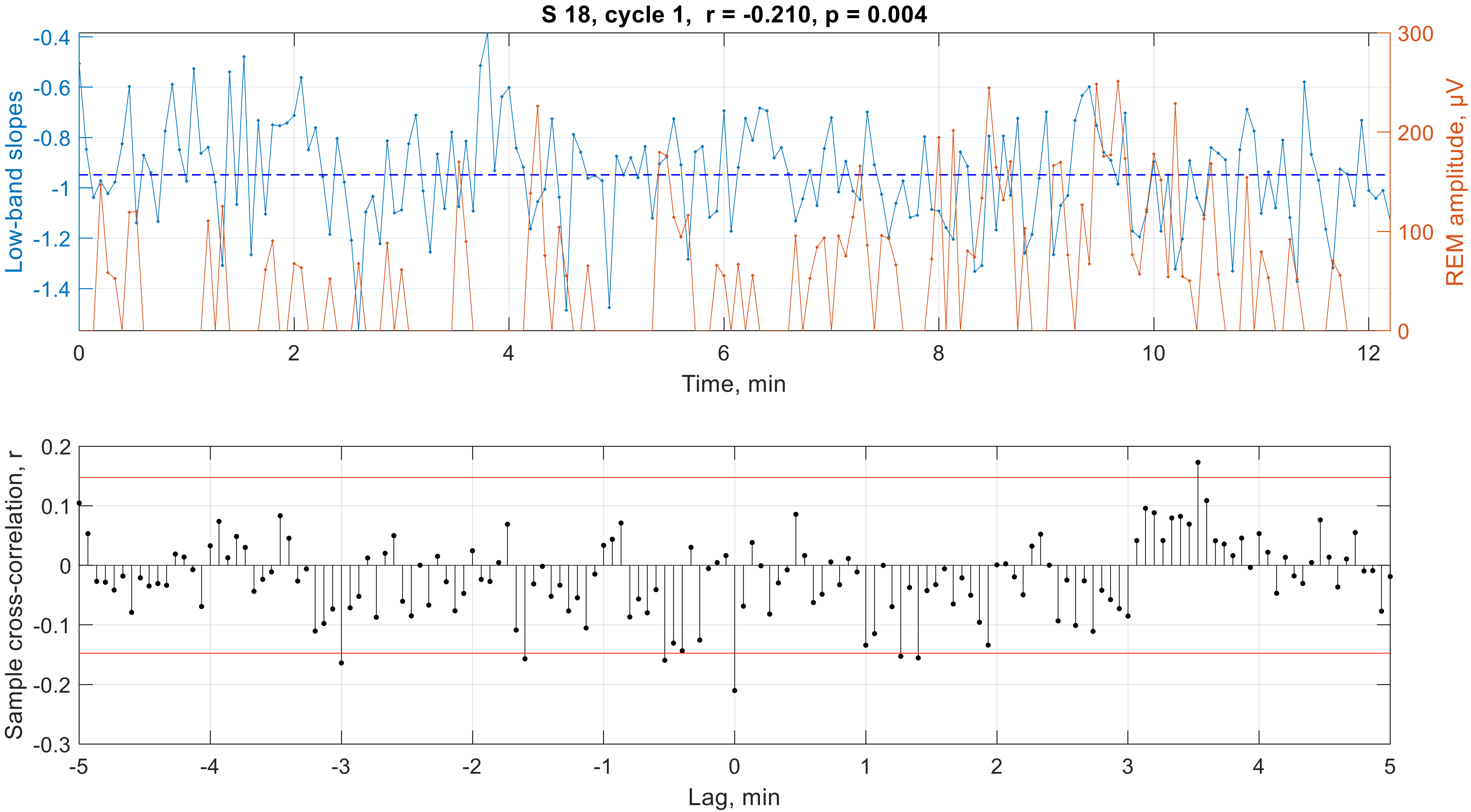

S 18, cycle 1, r = -0.210, p = 0.004
Low-band slopes
-0.4
-0.6
-0.8
-1
-1.2
-1.4
REM amplitude, μV
300
200
100
0
0
2
4
6
8
10
12
Time, min
Sample cross-correlation, r
0.2
0.1
0
-0.1
-0.2
-0.3
-5
-4
-3
-2
-1
0
1
2
3
4
5
Lag, min

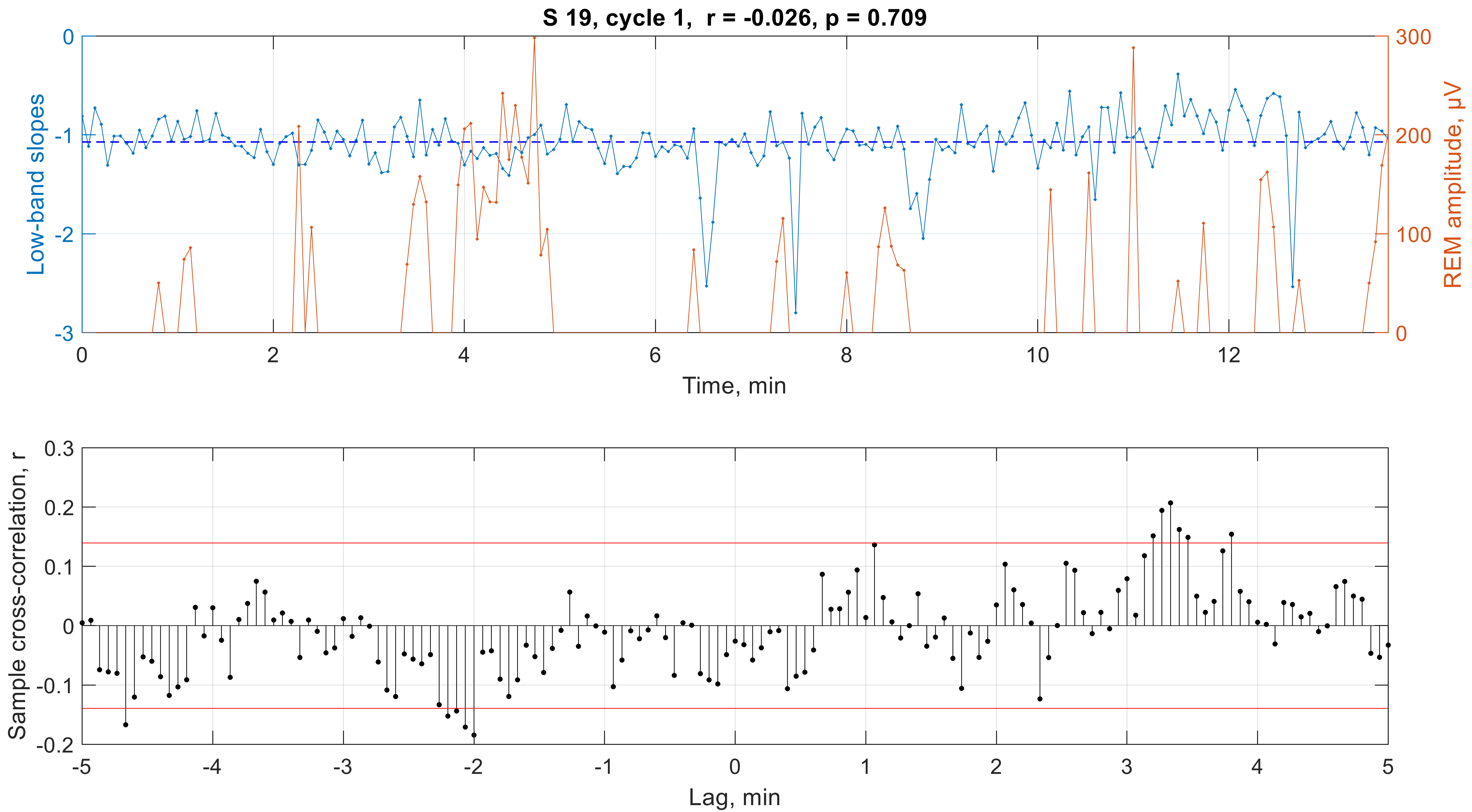

S 19, cycle 1, r = -0.026, p = 0.709
Low-band slopes
REM amplitude, µV
Time, min
0
-1
-2
-3
300
200
100
0
0
2
4
6
8
10
12
Sample cross-correlation, r
Lag, min
0.3
0.2
0.1
0
-0.1
-0.2
-5
-4
-3
-2
-1
0
1
2
3
4
5

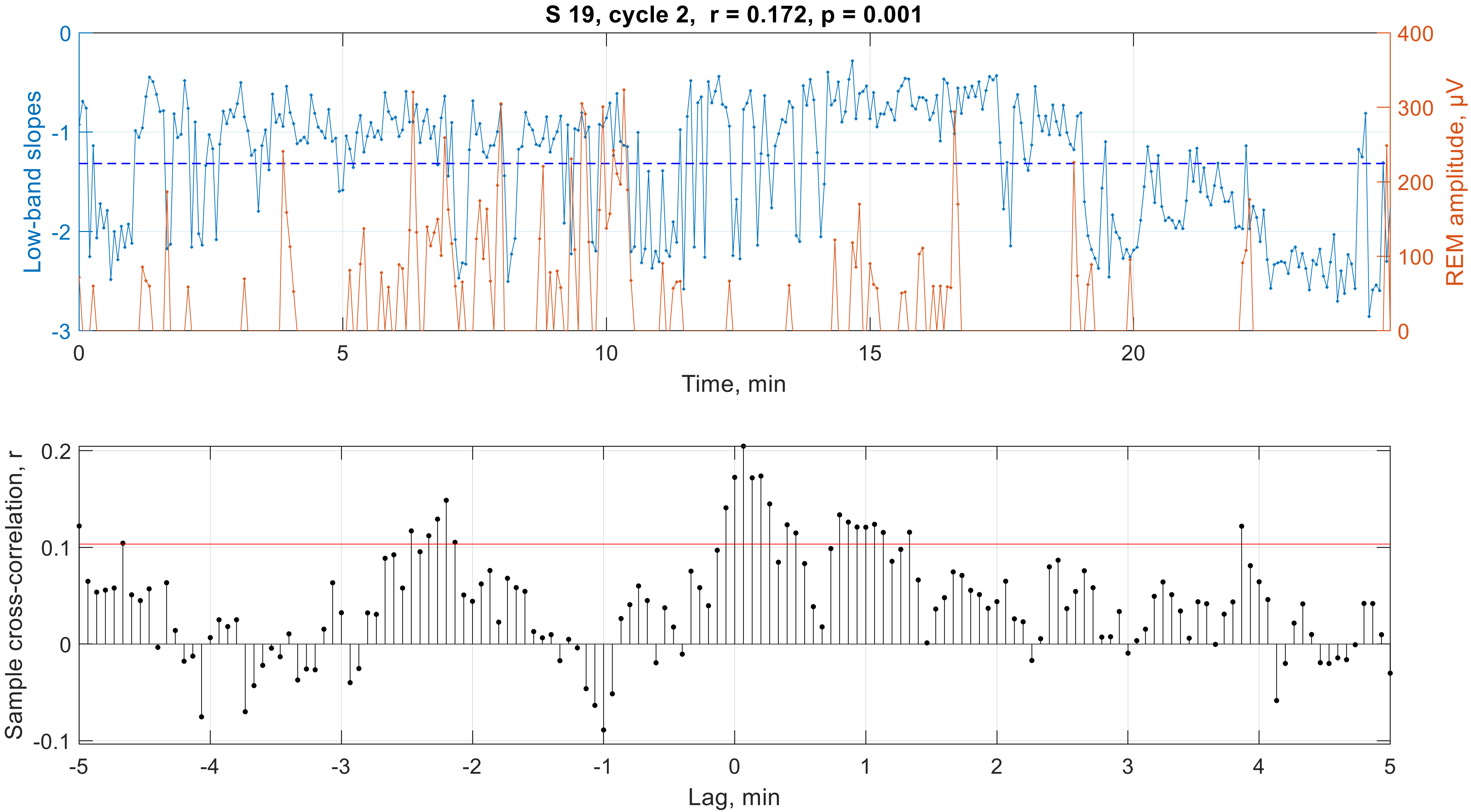

S 19, cycle 2, r = 0.172, p = 0.001
Low-band slopes
REM amplitude, μV
0
-1
-2
-3
400
300
200
100
0
0
5
10
15
20
Time, min
Sample cross-correlation, r
0.2
0.1
0
-0.1
-5
-4
-3
-2
-1
0
1
2
3
4
5
Lag, min

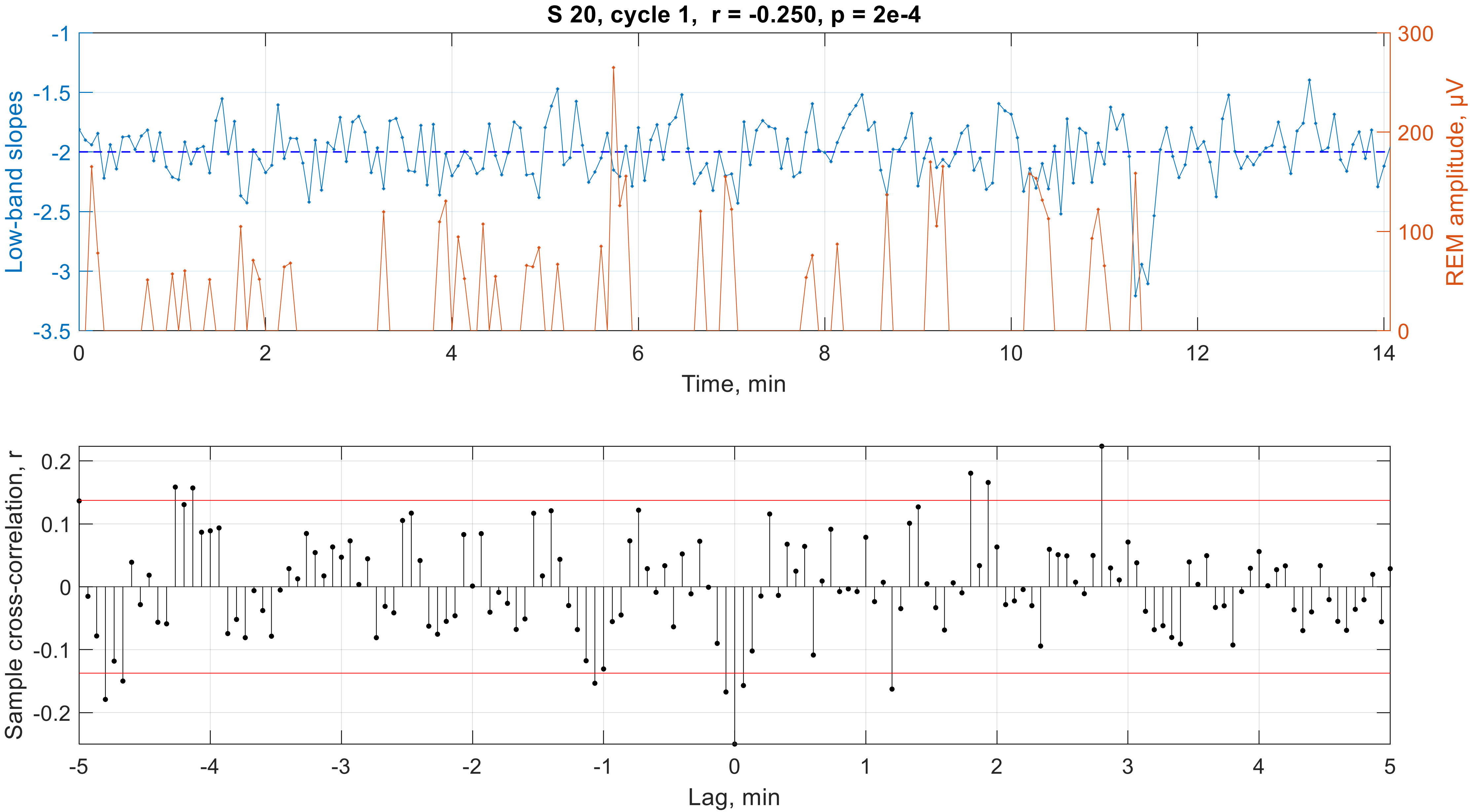

S 20, cycle 1, r = -0.250, p = 2e-4
Low-band slopes
REM amplitude, μV
Time, min
-1
-1.5
-2
-2.5
-3
-3.5
300
200
100
0
0
2
4
6
8
10
12
14
Sample cross-correlation, r
Lag, min
0.2
0.1
0
-0.1
-0.2
-5
-4
-3
-2
-1
0
1
2
3
4
5

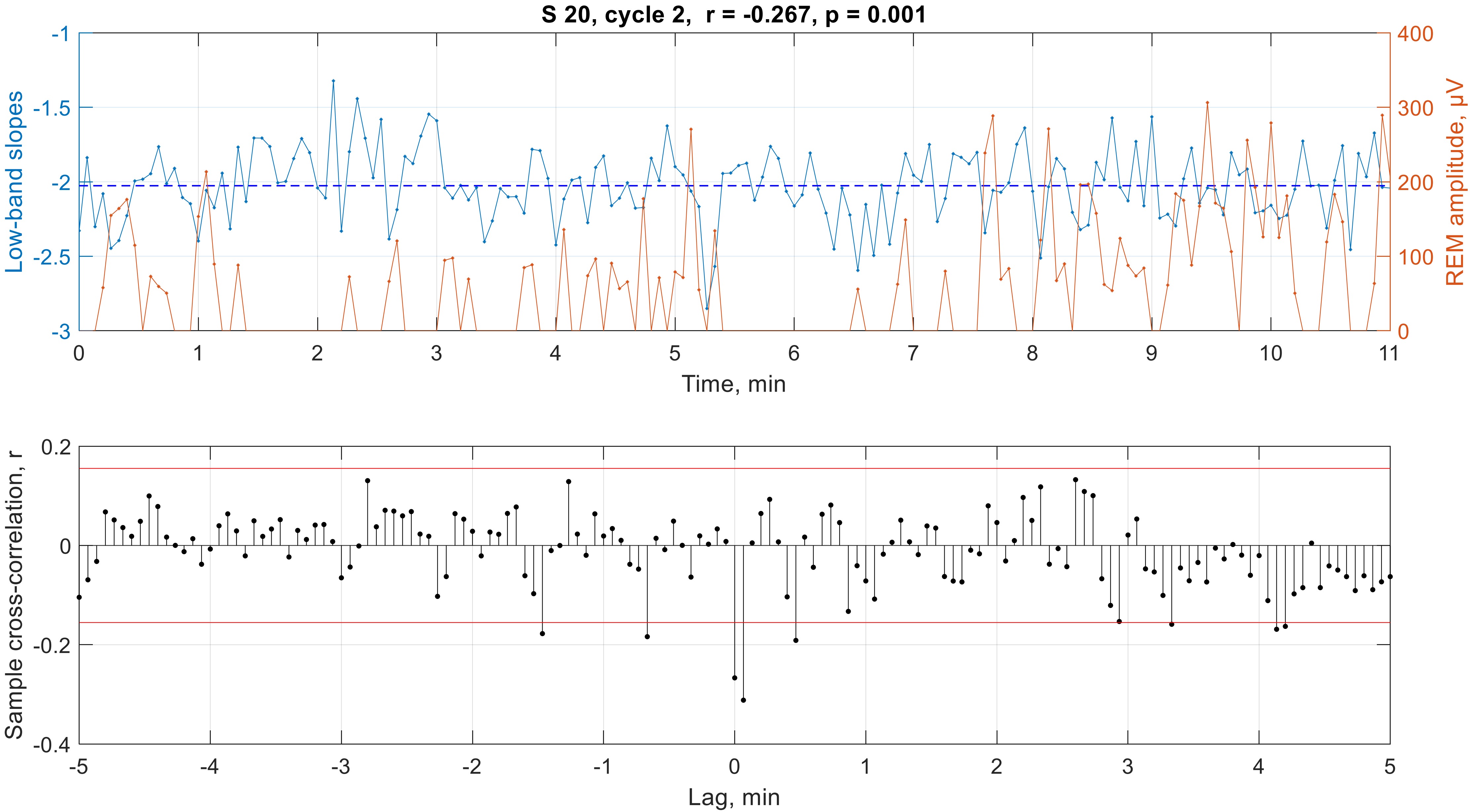

S 20, cycle 2, r = -0.267, p = 0.001
Low-band slopes
REM amplitude, μV
Time, min
Sample cross-correlation, r
Lag, min
